# The demonstration of a single origin for nodule evolution, with nodule engineering in a non-nodulating species

**DOI:** 10.64898/2026.05.31.728964

**Authors:** Anindya Kundu, R. Jordan Price, Esther Rosales Sanchez, Ivan Reynas-Llorens, Min-Yao Jhu, Jin-Peng Gao, Thiago Alexandre Moraes, Federico Marangelli, Cyril Libourel, Jean Keller, Jack Brooks, Wendy Harwood, Emma J. Wallington, Pierre-Marc Delaux, Richard J. Harrison, Giles E.D. Oldroyd

## Abstract

The nitrogen-fixing root-nodule symbiosis provides a sustainable source of nitrogen for plants within the Nitrogen (N)-fixing clade (NFC). A debate has raged over whether nodulation evolved once, with many losses or multiple times following a predisposition event. Here we demonstrate that nodule-organogenesis is fully conserved between an actinorhizal nodulator *Datisca glomerata* and the legume *Medicago truncatula*, showing entirely conserved programmes for *Nodule INception (NIN)-*controlled development leading to nodule emergence. Convergent losses of N-fixation within the NFC is associated with loss of *NIN* and we show the engineering of nodule-like development into strawberry, a non-nodulating member of the NFC, through the repair of *NIN-*functionality. Similar *NIN-*engineering resulted in altered-root developmental responses in barley. Our work is consistent with the single-gain hypothesis, where repair of NIN can recapitulate nodules in species within the NFC, demonstrating that understanding the ancestral state of nodulation facilitates its engineering.

## INTRODUCTION

The root nodule symbiosis (RNS) is a complex trait that has undergone gradual diversification, with it established in its most prolific form in legumes^1^. However, nitrogen-fixation, in the context of nodulation, is not limited to legumes, with nitrogen-fixing species also present within the Rosales, Fagales and Cucurbitales, which together with legumes form the monophyletic nitrogen-fixing clade (NFC) of plants^2^. In the Rosales, Fabales and Cucurbitales clades nodulating plants associate with a filamentous actinobacteria called *Frankia*, whereas legumes associate with single-celled rhizobia^3^. While the NFC contains all nitrogen-fixing plant species that nodulate, many members of this clade do not fix nitrogen, which has led to the debate: did nodulation evolve once, followed by many losses (the single gain hypothesis)^4,5^, or did a predisposition event(s) facilitate the emergence of nodulation on multiple independent occasions^2,6^?

Our understanding of nodulation is dominated by work in legumes where genetic model systems allowed ease of research. There, nodule organogenesis is initiated following recognition of nitrogen-fixing bacteria at the root surface, via the symbiosis signalling pathway, that promotes cytokinin signalling in the root cortex, inducing expression of the *Nodule Inception* (*NIN*) transcription factor^7^. *NIN* activates nodule organogenesis through the induction of *LBD16*, that also acts in lateral root development and promotes local auxin biosynthesis, and by the parallel induction of the *NF-Y* CCAAT-transcription factors, together promoting cell divisions in the pericycle and root cortex^8,9^. Nodule-specific functionality, associated with accommodation of N-fixing bacteria, is achieved by *NIN*-induction of *LSH* and *NOOT* transcription factors, two transcription factors that promote and sustain nodule identity^10^.

Here we demonstrate that nodulation must have evolved once, since its development and the genetic basis that controls it are highly conserved between two very diverse nodulating species: the actinorhizal nodulator *Datisca glomerata* (*D. glomerata*) belonging to Cucurbitales and the legume nodulator *Medicago truncatula* (*M. truncatula*) belonging to Fabales, which would represent homologous states in a single gain scenario and convergent states in a multiple gains scenario. The single gain scenario presumes loss of nodulation, through the loss of *NIN* ^4,5^, and we show repair of *NIN* in strawberry, whose ancestors lost nitrogen fixing ability, transfers nodule-like development. Comparable engineering in barley, a monocot outside the NFC, enhances proliferative root development. This work provides definitive proof of a single-gain hypothesis and demonstrates the feasibility for transfer to non-nodulating species within the N-fixing clade.

## RESULTS

### A conserved role of NIN in actinorhizal nodule organogenesis

*Datisca glomerata* (Cucurbitales) and *Dryas drummondii* (Rosales) represent actinorhizal nodulators that are intercellularly infected by clade 2 *Frankia* sp., defining the supposedly basal form of nodule development^11^. *D. glomerata* contains a rare duplication of *NIN* (Supplementary Fig. 1-2, Supplementary Table 1, Supplementary Dataset 1), with both copies showing expression in nodules (Fig. 1a, Supplementary Fig. 3), and *DatiscaNIN2* also having non-symbiotic expression in roots (Fig. 1a). *D. glomerata NIN* can partially complement the *M. truncatula nin-1* mutant, whereas *D. glomerata NIN2* cannot (Supplementary Fig. 4), implying possible neofunctionalization of this paralog. In contrast, *D. drummondii* encodes a single copy of *NIN*, the promoter of which shows nodulation-specific activity in *D. glomerata* and in *M. truncatula* (Fig. 1a, Supplementary Fig. 5), and can complement nodulation in *M. truncatula nin-1* (Supplementary Fig. 4). These results suggest that diverse actinorhizal *NINs* have comparable functions in nodulation as legume *NINs*. To determine whether the biological function of *NIN* is also conserved in actinorhizal species, we silenced *D. glomerata NIN* which leads to a complete loss of nodule organogenesis, with only occasional formation of proliferative root structures that do not get infected by *Frankia* Dg1 (Supplementary Fig. 3a-b). Similar loss of nodulation was previously observed for another actinorhizal species, *Casuarina glauca*^12^. Ectopic expression of *NIN* in *M. truncatula* results in the formation of pseudonodules: nodule like-structures with a large uninfected mass of cortical cells and peripheral vascular bundles induced in the absence of bacteria^13^. Comparably ectopic expression of both *DatiscaNIN/NIN2* and *DryasNIN* in *D. glomerata* resulted in the formation of highly proliferative pseudonodules (Fig. 1b-d, Supplementary Fig. 6). These pseudonodules showed upregulation of most *NIN*-dependent genes previously reported for nodule organogenesis in legumes and upregulated by ectopic expression of *MtNIN* in *M. truncatula*, including *LSH* and *NOOT,* that drive nodule identity (Fig. 1e, Supplementary Table 2, Supplementary Dataset 2-3). These genes were also suppressed following *NIN* RNAi in *D. glomerata* justifying their strong *NIN*-dependence during actinorhizal nodule development (Supplementary Fig. 3g). We conclude that *NIN* function is conserved in legume and actinorhizal nodulators and further we see evidence for a highly conserved nodule organogenesis program across the NFC, providing very strong evidence for a single origin in the evolution of nodulation.

**Fig. 1.**
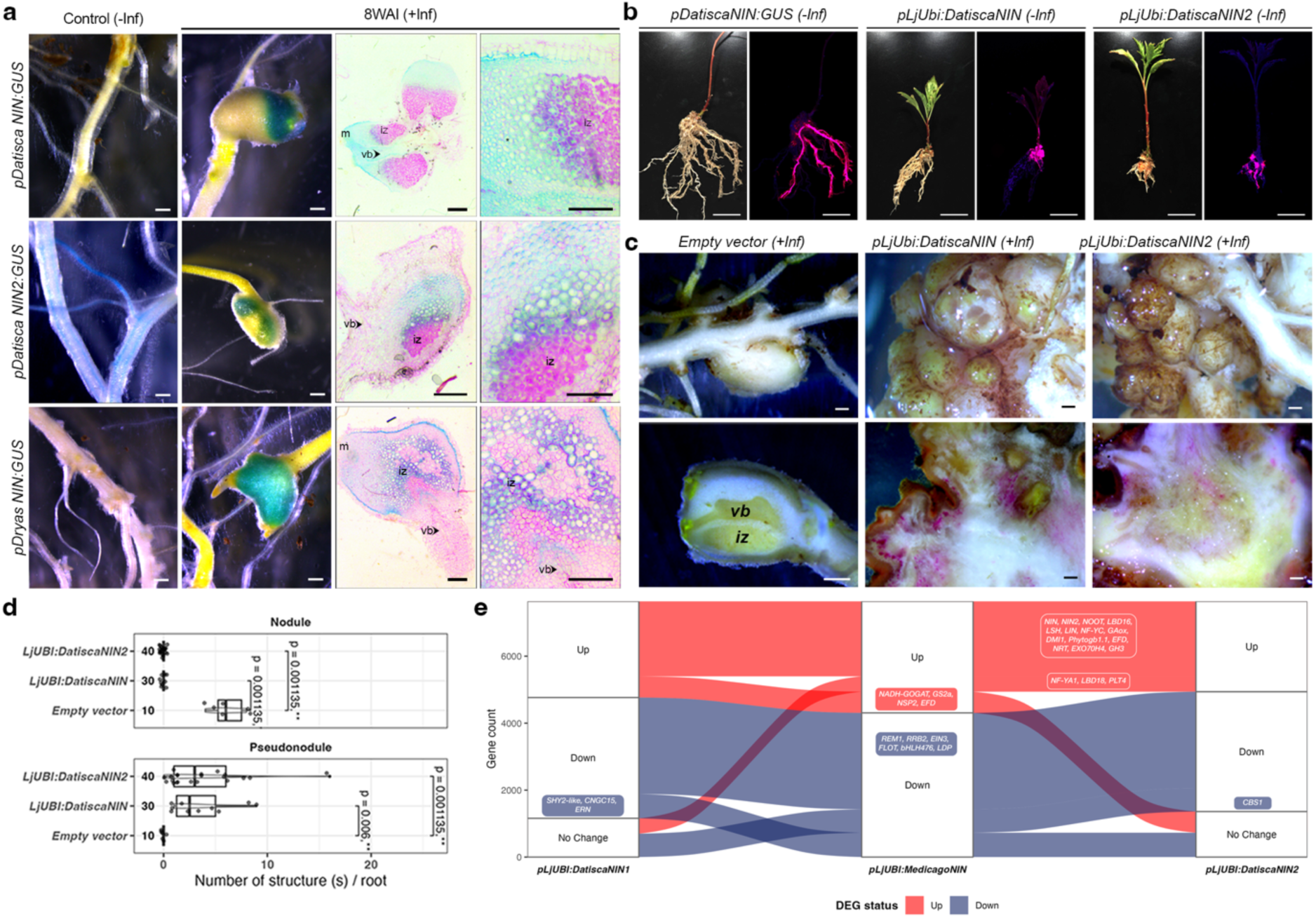
| A conserved role of NIN in actinorhizal nodule organogenesis. Transformed hairy roots of *D. glomerata* showing **a,** expression of *pDatiscaNIN:GUS*, *pDatiscaNIN2:GUS* and *pDryasNIN:GUS* in control (–Inf) and 8 weeks after infection (+Inf, 8WAI) with *Dg1 Frankia sp.*; ultrathin-sections of root nodules with GUS staining (blue) counter stained with ruthenium red (purple), Scale bar: 500 µm. **b,** Bright field and fluorescent images of composite plants transformed with *pDatiscaNIN:GUS*, *pLjUbi:DatiscaNIN* and *pLjUbi:DatiscaNIN2*, 8 weeks after transformation; mCherry (transformation marker, purple), Scale bar: 2 cm. **c,** Infected nodules on *D. glomerata* hairy-root transformed with empty vector and pseudonodules on roots transformed with *pLjUbi:DatiscaNIN* and *pLjUbi:DatiscaNIN2* (upper panel), with their corresponding half-sections (lower panel). Scale bar: 500µm. **d,** Boxplot showing the number of nodules and pseudonodules 8WAI with median (thick line) and second to third quartiles (box). Significance is indicated by p-values and asterisk derived from a Wilcoxon signed-rank test followed by Bonferroni correction, n = number of transformed roots analysed. **e,** Alluvial-plot illustrating the relationship between *pLjUBI:DatiscaNIN*, *pLjUBI:MedicagoNIN*^10,14^ and *pLjUbi:DatiscaNIN2* where the flow represents homologous gene expression patterns across three conditions, Up (> 0), Down (< 0) or No Change (0). The width of the flow indicates the number of DEGs (Differentially expressed genes) shared. m, meristem; vb, vascular bundle; iz, infection zone. See Supplementary Dataset 2-3.

### A basal role for auxin in regulating *NIN* expression during nodule development

Legumes show spatio-temporal control of *NIN* expression through *cis*-regulatory elements in the *NIN* promoter, controlled by CYCLOPS and cytokinin-dependent Response Regulators, that coordinate *NIN* expression with bacterial infection^7,15^. This coupling of cytokinin signalling to nodule organogenesis appears to be a unique adaptation in legumes^16,17^. Metanalysis of the promoter regions (–2.5kb) of *NIN-like proteins* (*NLPs*) within and outside the NFC showed the presence of putative auxin responsive element (*AuxRE*) pairs which were either in the form of direct, indirect or everted repeats, with inter-motif distances ranging from 1-50bps (Fig. 2a, Supplementary Fig. 7, Supplementary Dataset 4). While the presence of additional *AuxRE* beyond –2.5kb cannot be ruled out, most auxin responsive genes show the presence of functional *AuxRE* closer to their transcriptional start sites^18^. There was a strong positional bias for *AuxRE* in the nodulating *NIN* promoters near –900bp and at –2.4kb which were otherwise missing in non-nodulators (Fig. 2a, Supplementary Fig. 7). Exogenous treatment with the auxin analogue naphthaleneacetic acid (NAA) results in significant induction of *MtNIN* and *DgNIN/NIN2* expression in *M. truncatula* and *D. glomerata* respectively (Supplementary Fig. 8-10) and the spontaneous formation of pseudonodules (Supplementary Fig. 11), analogous to what has been previously reported^19^. *AuxREs* are also present in *NIN* promoters of legumes, including *M. truncatula,* and mutated forms of the *pMtNINsyn_*promoter in *AuxRE* showed attenuated complementation of *nin-1* with uninfected nodule primordia lacking progression into fully functional nodules (Fig. 2b, Supplementary Fig. 10, Supplementary Fig. 12), as well as attenuated induction following exogenous auxin (NAA) treatment (Supplementary Fig. 8a-d). *AuxRE* mutation resulted in reduced epidermal *pMtNIN* activation leading to loss of an auxin maxima affecting infection progression (Supplementary Fig. 12-13). Such *AuxRE-*mutated *pMtNINsyn* promoters also showed reduced induction by NIN in a transactivation assay in *N. benthamiana* (Supplementary Fig. 10c), implying a function for auxin in activating the *M. truncatula NIN* promoter in this heterologous system. We conclude that auxin contributes to the induction of *NIN,* through the presence of *AuxREs* in the promoters of *NINs* in legumes and actinorhizal nodulators. Where cytokinin responsive elements are missing^20^, auxin regulation of *NIN* may provide the main mode of *NIN* activation, such as in actinorhizal species, and this is supported by colocalization of *pDR5:GUS* and *pNIN:GUS* in *D. glomerata* (Fig. 1, Supplementary Fig. 14-15).

**Fig. 2.**
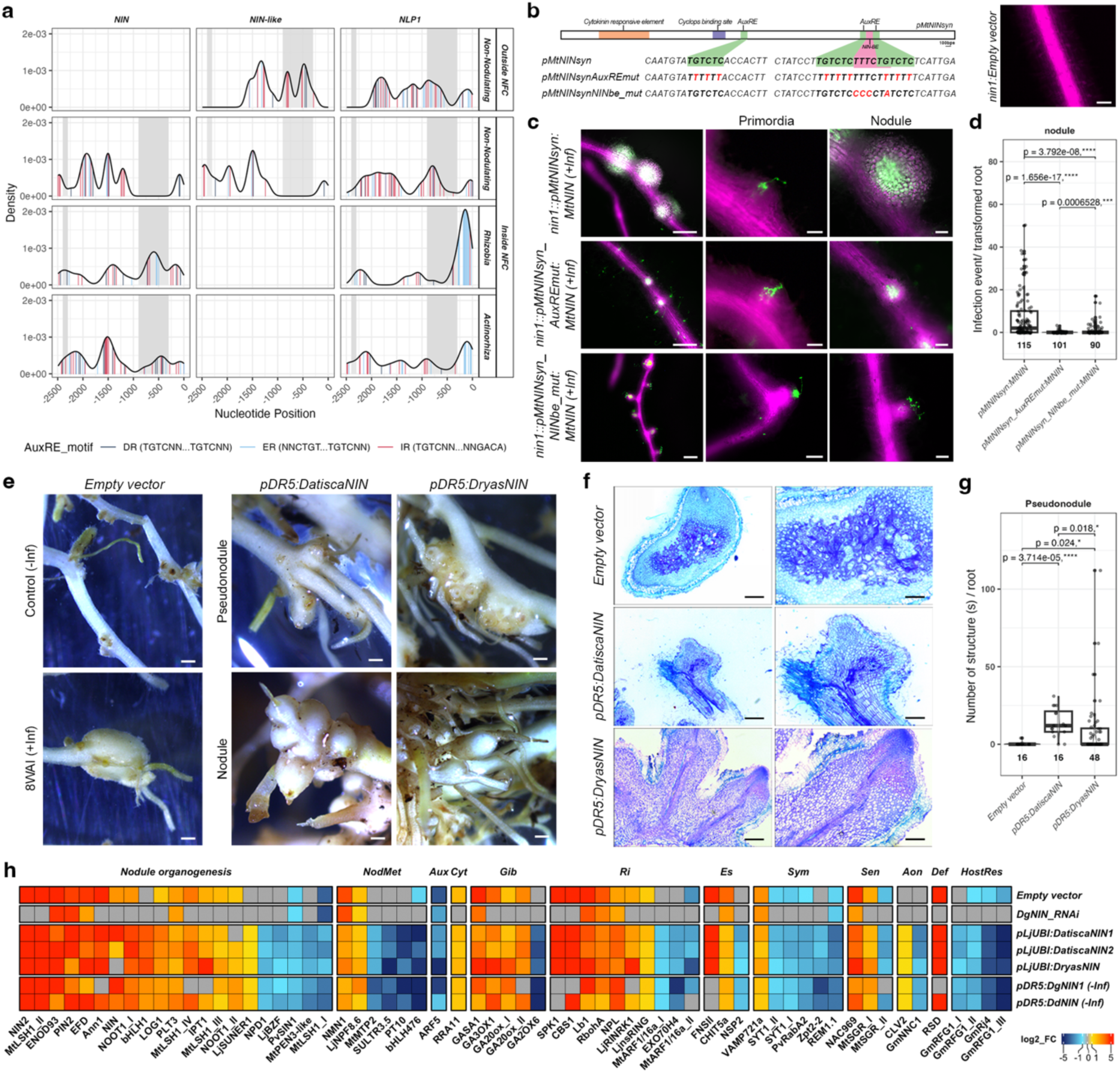
| Correlation between auxin and NIN function plays an essential role during nodule development. **a**, Density of *AuxRE* pairs in the promoters of *NIN*, *NIN-like* and *NLP1* from within and outside the NFC where colour of the lines indicates the orientation of *AuxRE* motifs with a p-value threshold <0.01 and grey boxes represent the unique presence of *AuxRE* in *NIN* promoters of nodulators. See Supplementary Dataset 4. **b,** Pictographic depiction of the synthetic *pMtNIN* and its mutant versions where the mutated nucleotides in the promoter region are represented in red and different *cis*-regulatory elements are highlighted with coloured boxes. A representative image showing *nin1* mutant transformed with empty vector. **c,** Representative images for *M. truncatula nin1* transformed with *pMtNINsyn:MtNIN*, *pMtNINsyn_AuxREmut:MtNIN* and *pMtNINsyn_NINbe_mut:MtNIN* showing primordia and nodule morphology 4WAI with *Sinorhizobium meliloti* 2011-GFP (green) where mCherry (magenta) is the transformation marker. Scale bar, 100 µm. **d,** The boxplot represents the number of nodules per *hairy-root* transformed root in *M. truncatula nin1* mutant. **e**, Hairy-root transformed *D. glomerata* plants transformed with empty vector, *pDR5:DatiscaNIN* and *pDR5:DryasNIN* under control (–Inf) and 8WAI (+Inf, *Dg1 Frankia sp.*). showing pseudonodule and nodule formation. Scale bar, 500 µm. **f,** Ultrathin sections of *D. glomerata* nodule and pseudonodules stained with toluidine blue showing ultrastructure. Scale bar, 500 µm (lower mag) and 250 µm (higher mag). **g,** Boxplot represents the number of pseudonodules per transformed root in *D. glomerata*. **h,** Heatmap of selected genes differentially regulated under control (empty vector), *DgNIN-RNAi*, *pDR5:DatiscaNIN* (–Inf), *pDR5:DryasNIN* (–Inf), *pLjUbi:DatiscaNIN* and *pLjUbi:DatiscaNIN2* where NodMet (Nodule metabolism), Aux (Auxin), Cyt (Cytokinin), Gib (Gibberellin), Ri (Rhizobial infection), Es (Early signaling), Sym (Symbiosome formation), Sen (Senescence), Aon (Autoregulation of Nodulation), Def (Defense) and HostRes (Host response). Gene functional nomenclature^22^. (D and G) In the boxplot, median (thick line) and second to third quartiles (box) and Wilcoxon signed-rank test followed by Bonferroni correction represents statistical significance where the p-value and the asterisk are represented over the brackets, n = number of transformed roots scored. See Supplementary Dataset 2.

To test the relevance of auxin regulation of *NIN* in actinorhizal species, we created a synthetic *NIN* driven only by an auxin-responsive promoter (*pDR5*). Hairy-root transformation of *D. glomerata* roots with auxin-regulated *DatiscaNIN* and *DryasNIN* led to the formation of spontaneous pseudonodules on primary roots (Fig. 2e), without any bacterial infection (Fig. 2f-g) which is a shift from the pseudonodules observed at the base of the stem upon *NIN* overexpression (Fig. 1b-c). Spontaneous pseudonodules were previously reported for the actinorhizal nodulators *Casuarina glauca* and *Discaria trinervis,* but here auto-active *CCaMK* triggered organogenesis^21^. Actinorhizal nodules share close homology to lateral roots, and hence it is difficult to differentiate between a modified lateral root and a pseudonodule. However, transcriptomic analysis of structures induced by *pDR5-NIN* showed many genes activated that show hallmarks of nodulation in legumes (Fig. 2h-i), and close similarity in transcriptomic responses between ectopic *NIN* expression and auxin-activated *NIN,* with a Spearman’s correlation score of 0.7-0.9. Unlike in *D. glomerata*, *pDR5:MtNIN* or *pDR5:DryasNIN* in *M. truncatula* failed to induce any organogenesis in *nin-1* (Supplementary Fig. 16). Since *pDR5:GUS* alone has attenuated activity in the *nin-1* mutant (Supplementary Fig. 17), and the first *pMtNINsyn(–5kb)* promoter region of *pMtNIN* contains several *cis*-regulatory elements required for infection progression^7^, we created a minimal promoter by fusing synthetic *pDR5* with *pMtNINsyn(–5kb)* (Supplementary Fig. 18). Expressing *Mt*NIN under *pDR5_ pMtNINsyn(–5kb)* in the cytokinin perception mutant *cre1-1* of *M. truncatula* resulted in the formation of mature nodules, indicating that auxin activation of *NIN* can bypass the cytokinin requirement for nodule organogenesis in legumes. We conclude that auxin provides the principal mode for activation of *NIN* leading to nodulation in actinorhizal species, and we see evidence that this function still exists in legumes, even with cytokinin principally superseding it.

### Synthetic auxin responsive *NIN* can induce nodule organogenesis in strawberry

Our work demonstrates deep conservation of *NIN* function in the NFC, strongly suggesting a single origin for the evolution of nodulation. Although non-nodulators within the NFC, like *Fragaria vesca* (strawberry), have lost NIN, they are expected to retain certain genetic frameworks similar to closely related nodulating relatives like *D. drummondii.* To test this, we attempted to repair *NIN-*functionality in strawberry. Ectopic expression of *DryasNIN* in strawberry was toxic, causing necrosis of callus without any plant regeneration (Supplementary Table 3), and this toxicity is not species-specific, as ectopic expression of *MtNIN*, *DatiscaNIN* and *DryasNIN* all led to senescence in transient leaf assays of *Nicotiana benthamiana* (Supplementary Fig. 19). Considering the ancestral function of auxin in regulating *NIN,* we attempted to generate stable lines of strawberry expressing *pDR5:DryasNIN,* for which, unlike ectopic *NIN*, transgenic lines regenerated normally. The presence of *NIN* in *F. vesca* dramatically enhanced the response to auxin treatment, showing formation of pseudonodules that were significantly higher in number and larger in size, as compared to lines only expressing the control vector (Fig. 3a-c). These pseudonodules were morphologically similar to nodules, showing a large ‘flattened’ top, containing multiple vascular strands, differentiating them from lateral root primordia and analogous to spontaneous pseudonodules in legumes, where similar peripheral vascular bundles were previously noted^13,23^. The induction of these pseudonodules was independent of nitrate-availability in the media (Supplementary Fig. 20, Supplementary Dataset 5). Despite the presence of the *AuxRE* in the *Dryas* and *Datisca NIN* promoters, neither *pDryasNIN:DryasNIN* or *pDatiscaNIN:DatiscaNIN* engineering in strawberry resulted in pseudonodules (Supplementary Fig. 21), alluding to the difficulty in cross species activation using native promoters due to mismatched *cis-* and *trans-* regulatory elements ^24^.

**Fig. 3.**
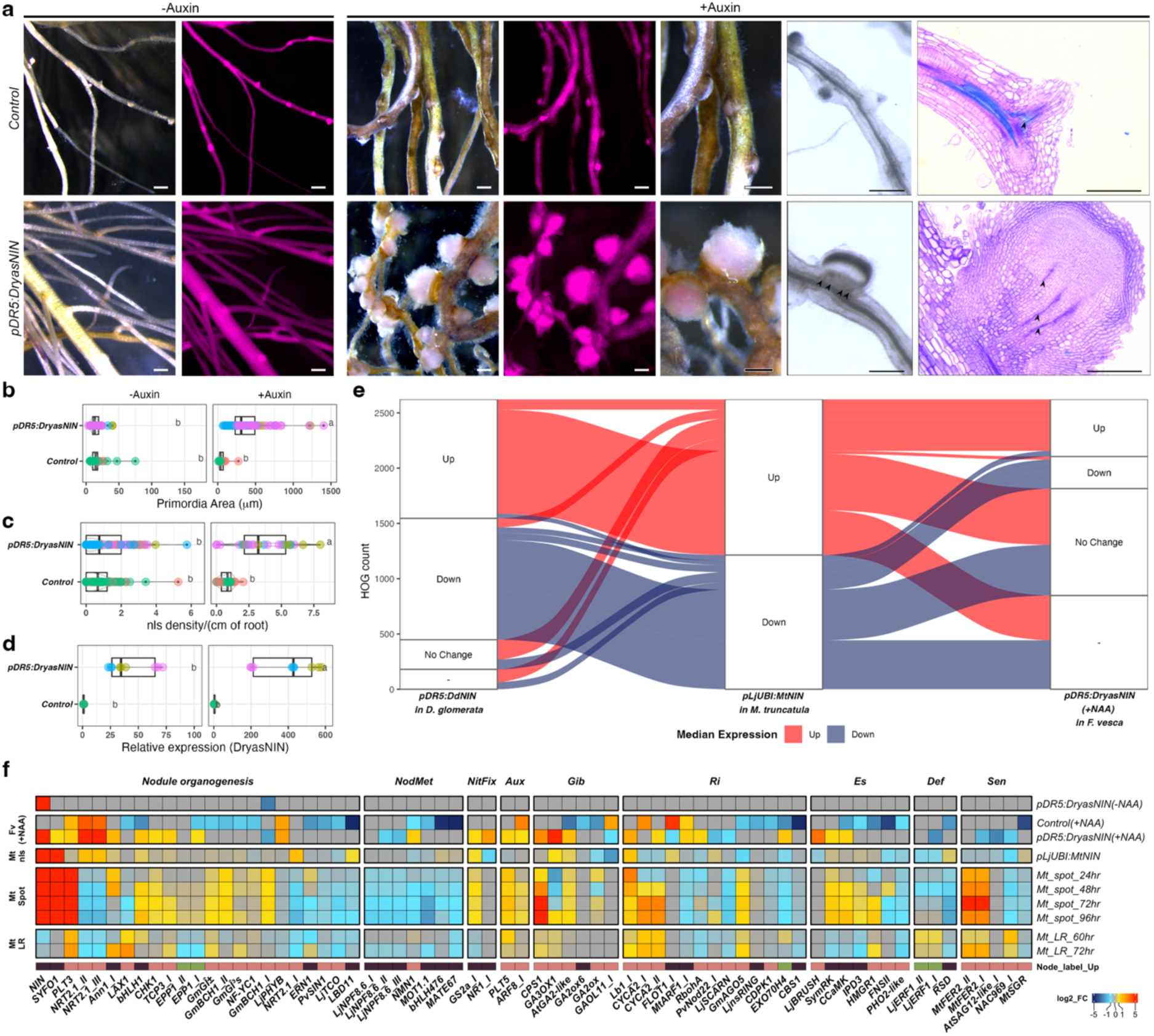
| Synthetic auxin responsive element driving NIN can induce nodule-like structures (nls) in *Fragaria vesca* (Fv, strawberry). **a**, Stereofluorescence images of *F. vesca* stable lines transformed with Control (*pDryasRPG:GUS*) and *pDR5:DryasNIN-pDryasRPG:GUS* treated under –Auxin and +Auxin (100 μM NAA, 1-Naphthaleneacetic acid) harvested 4 weeks post treatment (4 WPT) where mCherry (transformation marker, purple). Bright field and Toluidine blue stained ultrathin section of the root showing vascular bundle in the root and nodule-like structures, nls (arrowhead). Scale bar, 500 μm and 250 μm (toluidine blue sections). Boxplot represents the **b,** nls density **c,** Primordia area per transformed root and **d,** Expression of *DryasNIN* measured by quantitative reverse transcription polymerase chain reaction, qRT-PCR relative to reference gene *FvHistone4* respectively for mock and auxin treated roots, n = 9 where median (thick line), second to third quartiles (box) and independent lines (coloured dots). Single Letters represent statistically significant groupings analyzed by two-way ANOVA followed by TukeyHSD, p < 0.05. **e,** Alluvial-plot illustrating the relationship between median gene expression in each hierarchical orthogroup (HOG) for *pLjUBI:MtNIN* in *M. truncatula*, *pDR5:DryasNIN* in *D. glomerata* and *pDR5:DryasNIN* (+NAA) in *F. vesca* where the Up (>0) and Down (<0) HOGs in *M. truncatula* are compared against Up (>0) and Down (<0) HOGs in *D. glomerata* and *F. vesca pDR5:DryasNIN* datasets. No Change (= 0) and Orthologue absent (−). **f,** Heat-map of selected genes Up (>0) and Down (<0) regulated under *pDR5:DryasNIN* (+NAA) in *F. vesca* and *pLjUBI:MtNIN* in *M. truncatula*^10,14^, plotted with Control (+NAA), *pDR5:DryasNIN* (–NAA), *M. truncatula* spot inoculation and lateral root time-point dataset^9^, where NodMet (Nodule metabolism), NitFix (Nitrogen fixation), Aux (Auxin), Gib (Gibberellin), Ri (Rhizobial infection), Es (Early signalling), Def (Defence) and Sen (Senescence). Gene functional nomenclature^22^. See Supplementary Dataset 5 and Supplementary Fig. 20. The “Node label Up” is based on the ancestral HOG identity^26^, where coloured boxes represent Ancestral (dark brown), Legume specific (light brown), Actinorhizal specific (green) HOGs.

Previously, pseudonodules in legumes have largely been distinguished from lateral root primordia by their shape and the nature of their vasculature^8,13,25^. The transcriptome acts as a secondary marker for nodule-identity and therefore we compared the transcriptomes of conditions that promote pseudonodules in *M. truncatula, D. glomerata* and strawberry (Fig. 3e). Using the *NIN*-induced transcriptome in *M. truncatula* as the comparator, we see 82.9% of orthogroups in *D. glomerata* showing comparable *NIN*-induced expression and 30.7% showing comparable *NIN-*induced expression in strawberry. However, in strawberry many of the *NIN-*regulated genes from *M. truncatula* do not have orthogroups, many more than in *D. glomerata.* When we remove those genes lacking orthogroups from the comparison, then the DEGs comprises 45.5% of *NIN-*regulated genes in strawberry as compared to 89% in *D. glomerata*.

In a previous study, a metanalysis of transcriptomic data from 9 nodulators within the NFC allowed identification of hierarchical orthogroups (HOGs), progenitor genes that show comparable behaviours across the NFC, and thus act as core markers of nodule identity^26^. Among this group, many show differential expression in strawberry pseudonodules (Supplementary Fig. 20), including components of the symbiosis signalling pathway, *SymRK, EPP1* and *CCaMK,* genes activated by this pathway *Ann1/Ann2* and *SYFO1,* genes involved in nodule organogenesis *CHK1 (CRE1)*, *PLT3* and *NRT2.1* and genes associated with nitrogen fixation *GS2a* and *Lb1* (Fig. 3f, Supplementary Fig. 22). A correlation analysis for these genes further shows the presence of a significant overlap between the *M. truncatula* pseudonodules and strawberry pseudonodules, with a high correlation coefficient score (Supplementary Fig. 20, Supplementary Dataset 5). We conclude that a significant proportion of the *NIN*-regulon is conserved in strawberry and can be recruited by reintroducing the lost *NIN* to that lineage. However there remain components whose expression is not co-opted in these NIN-resurrected lines, such as *LSHs* and *NOOTs*^10^.

### Synthetic auxin responsive *NIN* enhances root organogenesis in barley

Outside of the NFC, cereal crops are strategically important for engineering nitrogen-fixation and we attempted to recapitulate the nodule-engineering in barley. Equivalent to what we previously observed in strawberry, attempts to engineer ectopic *NIN* expression into barley were unsuccessful, however we were able to regenerate transgenic lines of barley with *pDR5:DryasNIN.* These constructs leading to auxin-activation of *NIN* in these plants amplified the developmental responses to auxin alone, giving rise to the emergence of sizable lateral structures (Fig. 4a). These structures also showed multiple vasculature strands (Fig. 4b), but clearly lacked the same level of developmental organization that we saw in strawberry. Notably, the structures in barley had apparently friable tissues at their periphery (Fig. 4b). The transcriptomic comparison showed 6% of orthologous groups with comparable *NIN-*regulation in *M. truncatula* and barley, and if we remove the many genes that lack orthogroups, given that legumes and monocots are distantly related, the number is 15.5% (Fig. 4d, Supplementary Dataset 6). The HOGs that show comparable expression in barley and *M. truncatula* are principally genes associated with the auxin response (Fig. 4e, Supplementary Fig. 23) and lateral root development, with the notable exception of *GS2a,* a gene associated with nitrogen-fixation and recently shown to be associated with evolution of gene regulatory networks controlling the RNS^27^. Primarily, we found that *pDR5:DryasNIN* failed to induce many known targets in barley, which it could in strawberry. A recent report supports our observation, where *Medicago*NIN failed to induce the barley homolog of *SYFO1/2* in a transient *N. benthamiana* assay possibly due to missing *cis*-regulatory NIN binding elements in its promoter ^28^, which is upregulated in our strawberry dataset. We conclude that *NIN-*engineering into barley directs root development, but the extent of this impact is not as complete, as compared to strawberry, and lacks the nodule-like development that is apparent within plants in the NFC.

**Fig. 4.**
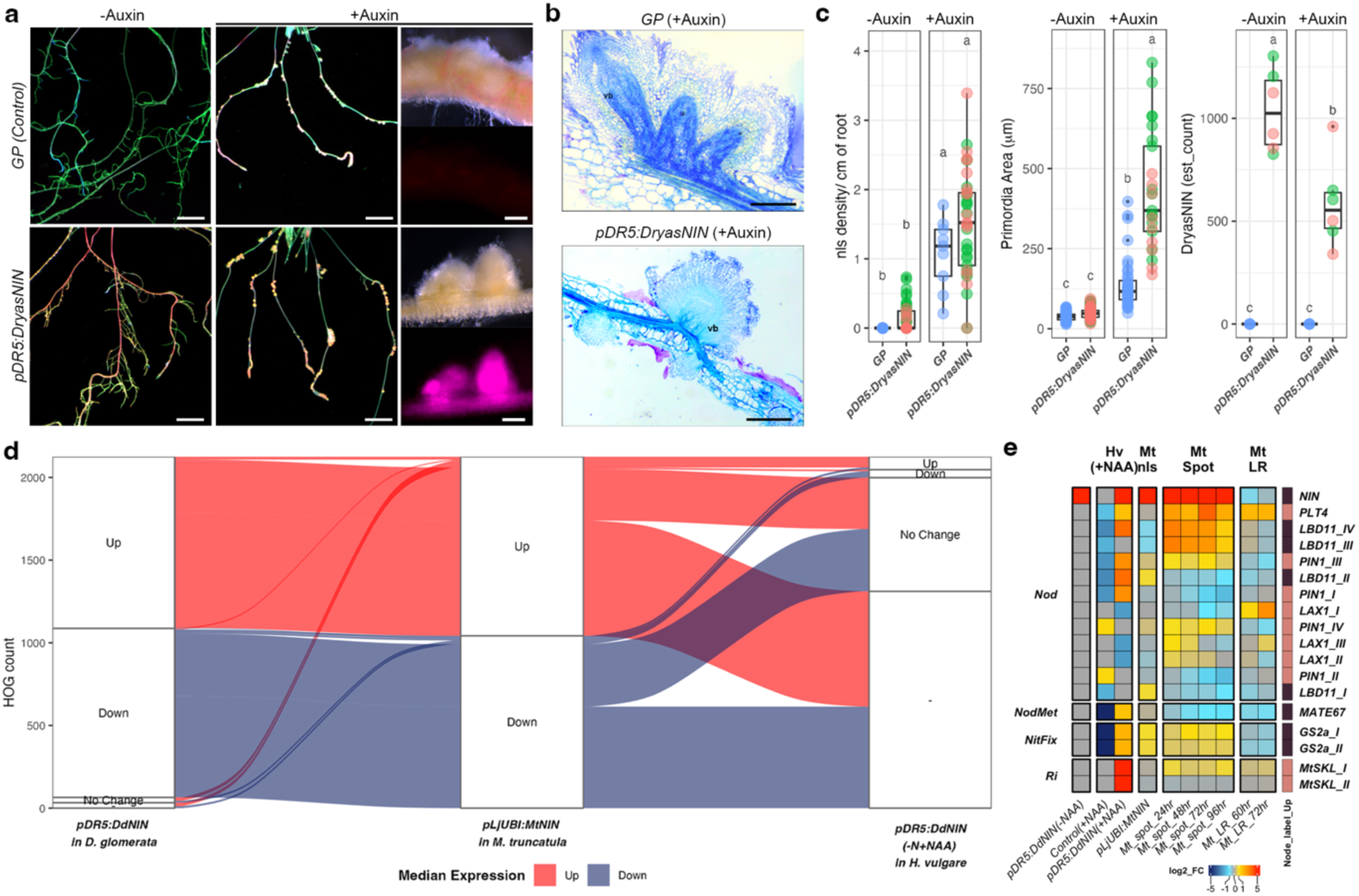
| Synthetic auxin responsive element driving *NIN* can enhance root organogenesis in *Hordeum vulgare* (*Hv*, barley). **a**, Imagequant fluorescence images of *H. vulgare* Golden promise, GP (Control) and stable lines transformed with *pDR5:DryasNIN* treated under –N-NAA (–Auxin) and – N+NAA (+Auxin, 50 μM NAA) harvested 3 weeks post treatment (3 WPT) where green, blue and red channel are merged. Scale bar, 2.4 cm. Stereofluorescence zoomed images of the nodule-like structures (nls) where mCherry (transformation marker, purple). Scale bar, 500 μm. **b,** Toluidine blue-stained ultrathin section of the nls showing vascular bundle (vb) for control (GP) and *pDR5:DryasNIN* roots. **c,** Boxplot represents nls density, primordia area per transformed root and *DryasNIN* estimated counts where median (thick line), second to third quartiles (box) and independent lines (coloured dots). Single Letters represent statistically significant groupings analyzed by two-way ANOVA followed by TukeyHSD, p < 0.05. **d,** Alluvial-plot illustrating the relationship between median gene expression in each heirarchical orthogroup (HOG) for *pLjUBI:MtNIN* in *M. truncatula*, *pDR5:DryasNIN* in *D. glomerata* and *pDR5:DryasNIN* (−N+NAA) in *H. vulgare* where the Up (>0) and Down (<0) HOGs in *M. truncatula* are compared against Up (>0) and Down (<0) HOGs in *D. glomerata* and *H. vulgare pDR5:DryasNIN* datasets. No Change (= 0) and Orthologue absent (−). **e,** Heat-map of selected genes Up (>0) and Down (<0) regulated under *pDR5:DryasNIN* (+NAA) in *H. vulgare* and *pLjUBI:MtNIN* in *M. truncatula*^10,14^, plotted with Control (+NAA), *pDR5:DryasNIN* (–NAA) under –N condition, *M. truncatula* spot inoculation and lateral root time-point dataset^9^, where Nod (Nodule organogenesis), NodMet (Nodule metabolism), NitFix (Nitrogen fixation) and Ri (Rhizobial infection). Gene functional nomenclature^22^. See Supplementary Dataset 6 and Supplementary Fig. 23. The “Node label Up” is based on the ancestral HOG identity^26^, where coloured box represent Ancestral (dark brown), and Legume specific (light brown) HOGs.

## DISCUSSION

There is little consensus on how nodulation evolved in angiosperms approximately 100mya: did it evolve many times, following a predisposition event, or did it emerge once, followed by many losses ^4,6,29,30^? Here we demonstrate that the central regulator of nodulation in legumes, *NIN,* has a conserved function in nodule organogenesis in two clades of nodulating species, namely *D. glomerata* from Cucurbitales and *D. drummondii* from Rosales, distantly related to legumes (Fig. 1, Supplementary Fig. 3). At the protein level, NIN function is sufficiently conserved to allow cross-complementation of *NIN* between Fagales ^12^, Rosales and Cucurbitales (Supplementary Fig. 4). Furthermore, both *NIN* function and its regulon is highly conserved across the NFC. Such an early emergence of *NIN* functionality, whose actions are alone sufficient to drive nodule-like organogenesis in legumes and in actinorhizal species, provides, we believe, compelling evidence for a single origin in the evolution of nodulation within the NFC.

Actinorhizal species, such as *D. glomerata* and *D. drummondii,* show nodules that are morphologically similar to lateral roots^11^. In a single origin hypothesis, because they are paraphyletic, the actinorhizal nodulators likely represent the basal state^1,30^, and as such it is a straight-forward summation that a modified lateral root represents the basal nodule. Such a proposal is consistent with genetic evidence in legumes showing conserved features of lateral root and nodule development^8,9^, and our observation of markers of lateral root development apparent in the *NIN-*regulon in all the species we studied. In this context, it is not surprising then to see that auxin acts as a principal initiator of *NIN* and nodulation in these tested nodulators, considering the central role for auxin in the initiation of lateral roots ^31^. What is however surprising is the absolute necessity of this modality of regulation to allow transfer of *NIN* across species (Fig. 2-4). We cannot yet explain the toxicity issue with *NIN* expression in shoots, but no doubt, as reported here, this issue has limited the success of *NIN* engineering to date. We demonstrate that when constrained to cells experiencing high auxin, NIN transfer is possible, and with it comes promotion of root development. In strawberry such development is nodule-like (Fig. 3), whereas in barley this is harder to argue (Fig. 4). But its transfer into strawberry combined with the recapitulation of nodule identity genes further supports a single-gain hypothesis where much of the framework for nodule development pre-exists needing a kick start by NIN. However, it is apparent that there are aspects of nodule development missing also in strawberry, for instance, we do not see induction of *LSH* or *NOOT*, key nodule regulators, both of which are activated in *D. glomerata* and *M. truncatula* nodules, and this likely reflects incremental secondary losses of pathway components or their modality of regulation, following loss of nodulation in the ancestors of strawberry. Accompanying a major loss, plants adapt to the newer environment through rewiring and neofunctionalization of pathways^32^. Nodule engineering by resurrecting *NIN* using ancestral auxin-regulation, represents a first step in transferring nodulation and with it N-fixation, using an evolution-guided approach. Although additional engineering steps are clearly still required, even in strawberry, this proof of concept offers a path to follow for future efforts. Our work implies that engineering N-fixation within the NFC is likely an easier first step, since it is resurrecting a pre-existing state, rather than transferring a new developmental state, as is the case in cereals.

## ACKNOWLEDGEMENTS

We want to acknowledge Prof. Katharina Pawlowski for providing the *D. glomerata* seeds and the batch of nodulated plants with clade 2 *Frankia* sp. Dg1. Prof. Rene Geurts and Dr. Rik Huisman for providing *Sinorhizobium meliloti Sm2011-GFP*. Kirsty McLeary for maintaining the strawberry lines. Eleni Soumpourou and her team for providing the vector backbones, assisting in DNA synthesis and genotyping the barley lines.

## FUNDING

This work was supported by the Bill and Melinda Gates Foundation and the UK Foreign, Commonwealth and Development Office (OPP1028264) and by Gates Ag One through Enabling Nutrient Symbioses in Agriculture (ENSA) project.

## AUTHOR CONTRIBUTIONS

AK, RJH and GEDO designed the experiments, analysed the data and wrote the manuscript. AK did the majority of experiments, prepared the figures and wrote the initial draft with inputs from RJH and GEDO. RJH and RJP acquired the seeds and plants to initiate the project. RJP and AK led genome sequencing. RJP, CL, JK analysed and annotated the newly sequenced genomes under the supervision of PMD. ERS and FM established the protocol for *D. glomerata* regeneration and AK did the *D. glomerata* hairy-root transformation. AK undertook the *M. truncatula* hairy-root transformation. AK, ERS and FM were responsible for raising the strawberry lines and performing the experiments. JB and EW subcultured and maintained the strawberry lines. TAM and AK analysed the RNAseq data and performed the *Nicotiana* experiments. IRL extracted the promoter regions for NLPs and did the initial *AuxRE* analysis. JG did the luciferase assay. MYJ performed the remapping of the *Medicago* transcriptomic data. WH generated the barley stable lines and AK conducted the barley experiments. EW, MYJ, TAM and PMD helped in editing. All the authors have read and approved the final draft.

## COMPETING INTERESTS

The work is filed for patent under the name Methods of genetically altering a plant nin-gene to be responsive to auxin. A Kundu, RJ Harrison – US Patent App. 18/816,990, 2025.

## SUPPLEMENTARY MATERIALS

Supplementary Fig. 1-23

Supplementary Tables 1-5

Supplementary Dataset 1-6

## Materials and Methods

### Plant material and growing condition for *Datisca glomerata*

In order to produce uniform plant materials through tissue culture, seeds were taken out from cold storage in 1.5ml microcentrifuge tubes and treated with 1ml solution A (70% ethanol and 0.05% SDS) for 1 min with gentle shaking. Following the treatment, the seeds were treated with 1 ml solution B (20% bleach and 0.1% SDS) for 5 mins with gentle shaking, then the seeds were thoroughly washed with sterile deionized water 3-4 times. The seeds in the microfuge tube are resuspended in water and covered with aluminium foil and kept in the dark for 7-10 days at 4°C. After the vernalization step, the seeds were plated on ½ MS (Murashige and Skoog) media with vitamins (Duchefa, Cat. No. M0222) at pH 5.8 and 0.8% agarose (Sigma, Cat. No. A7921) in 9cm Petri dishes. The plates were sealed with micropore tapes and kept at 25°C under lights with 16h-photoperiod till they germinate. Germinated seedlings were transferred to 12cm square plates till they are well established with 4-6 leaves (approx. 4-6 weeks after germination). Young fresh leaves were collected from the apical part of the plant and chopped into 4-5 mm pieces under sterile condition. The leaf pieces were transferred to Datisca Callusing Media (DCM) containing MS supplemented with 1.5 mg/L BAP (6-Benzylaminopurine; Duchefa, Cat. No. B0904), 0.1 mg/L of NAA (α-naphthaleneacetic acid; Sigma, Cat. No. N0640), 5 g/L of Agargel (Sigma, Cat. No. A3301) and 30 g/L of sucrose (Merck, Cat. No. S0389) and adjusted to pH 5.8. Leaf pieces were grown for 10-15 days before they are transferred to Datisca Regeneration Medium (DRM) petri dishes (MS-based medium supplemented with 1.5 mg/L of BAP, 0.1 mg/L of IBA (Sigma, Cat. No. I5386), 0.1 mg/L of GA3 (Duchefa, Cat. No. G0907), 5 g/L of Agargel and 30 g/L of sucrose and adjusted to pH 5.8. After 8-12 weeks on DRM, shoots regenerated from the callus starts forming shoots which were excised from the callus and moved into Datisca Rooting Medium (MS-based medium supplemented with 1 mg/L of IBA, 8 g/L of Phytoagar (Duchefa, Cat. No. P1003) containing 30 g/L of sucrose and adjusted to pH 5.8. The plants started rooting within next 3-4 weeks and matured plants could be harvested for hairy-root transformation.

### Hairy-root transformation of *D. glomerata*

For hairy root transformation, the cultures of *Agrobacterium rhizogenes QUA1* harbouring the plasmid of choice were grown overnight at 28°C in LB-media with appropriate antibiotics and shaken at 220 rpm. The cultures were centrifuged at 4000g and resuspended in fresh LB media before plating them on LB plates containing appropriate antibiotic selection. The plates were incubated overnight at 28°C to obtain a confluent lawn of the bacteria. Fresh *D. glomerata* regenerated plant shoots are harvested using a sharp sterile scalpel and inoculated with *A. rhizogenes* culture from the plate. The inoculated plantlets are then transferred to ½ MS plates supplemented with 10% sucrose, 0.5 gm/L MES (Sigma, Cat. No. 69889) containing 0.8% agar and adjusted to pH 5.8. The plates were sealed with micropore tapes and 90% covered with aluminium foil and kept at 22°C for 2 days with 16h-photoperiod. After 2 days the explants were cleaned of any excess bacterial culture using sterile tissue paper and moved to new ½ MS plates supplemented with 0.5 gm/L MES and adjusted to pH 5.8. The plants were maintained under 16h-photoperiod at 22°C till they start forming roots which are checked for fluorescence under the stereo-fluorescence microscope DX105 F (Leica). Once the plant had sufficient transformed roots with fluorescence, the non-fluorescent untransformed roots were excised to encourage transformed root growth. After ∼4-6 weeks, the plantlets were moved into autoclaved perlite in pots and grown for another 10 days under nitrogen deprived conditions under high humidity (closed cover in micro propagator) prior to inoculation.

### Nodulation assay for *D. glomerata*

Actively nodulating *Datisca glomerata* plants with *Frankia* clade 2 *Dg1*^33^ were grown in glasshouse under 16h-photoperiod at 24-22°C. The fresh nodules were harvested from the plants and thoroughly washed with sterile water before they are crushed with a mortar and pestle to form a consistent slurry (∼5 developed nodules per inoculated plant). The hairy-root transformed plants were inoculated by brief immersion into the nodule slurry. Plants are then moved onto a sterile soil mix prepared with perlite: low nutrient peat-based soil (Levington, BC0010) (1:1) and maintained 16h-photoperiod, 600mWm^-2^nm^-1^ at 24-22°C for 8-10 weeks prior to nodulation assessment.

### Growth condition for *Medicago truncatula*

Plant material used in the study: *M. truncatula* ecotype A17 was used as a wild-type background for hairy-root transformation and for comparison to *nin1*, *nin1::pENOD11:GUS* ^34^ and *cre1-1* ^35^.

### Hairy-root transformation of *M. truncatula*

Seeds were scarified with concentrated sulphuric acid (∼99%) in a microfuge tube for 1-2 mins. Following the scarification, the sulphuric acid is thoroughly washed with sterile water 4-5 times to remove any trace of the acid. Scarified seeds were surface sterilized using 15% (v/v) bleach solution and 1% (v/v) Tween-20 (Merck, Cat. No. P2287) for 15 mins. The seeds are thoroughly washed with sterile water and kept at 4°C overnight in dark. Next day the seeds were plated on water droplets and allowed to germinate in dark at 22°C. *Agrobacterium rhizogenes* harboring the plasmid of choice are grown as described above. The hairy root transformation is done according to a previously published protocol^36^. After the transformation, the plants were transferred to FPmod plates^9^. The plants were maintained at 22°C with 16h-photoperiod, 600mWm^-2^nm^-1^ till they start forming transformed hairy-roots which are checked under a stereo-fluorescence microscope DX105 F (Leica).

### Nodulation assay for *M. truncatula*

*Sinorhizobium melliloti 2011* expressing constitutive GFP^7^ was a kind gift from Rene Geurts, Wageningen University, Netherlands. The *S. meliloti 2011* was grown in YEM media for 2 days at 28°C and diluted to a final concentration of 0.02 OD 600nm using buffered nodulation media, BNM^37^. After maintaining the hairy-root transformed plants in nutrient deprived condition for 3-4 days in perlite, the plants were inoculated with the *S. meliloti* culture (2ml of 0.02OD 600nm) and maintained in perlite for up to 4 weeks for nodule quantification and histochemical staining.

### Spot inoculation for *M. truncatula*

Hairy-root transformed plants with sufficient transformed roots (with visible mCherry expression, selected under stereofluorescence microscope) were maintained on BNM-N+P plates with 1μM AVG for 4 days. The transformed roots were spot inoculated with 1μl *S. meliloti* culture (0.02OD 600nm) mixed with 3μM luteolin according to Schiessl et. al., 2019^9^.

### Stable transformation of *Fragaria vesca* (woodland strawberry)

#### Plant materials and *in vitro* propagation

Transformation of *F. vesca cv. Hawaii 4* with *Agrobacterium tumefaciens* EHA105 carrying appropriate binary vector was done as previously described^38^.Regenerated plants were grown on media containing selective agent and assessed under the stereofluorescence microscope for consistent expression of a fluorescence marker for transformation, to avoid chimeric events. After 4 weeks in SMM tubes, plants were mature enough to be moved to sterile honey jars (5 plants per jar). Honey jars with SMM medium or Strawberry Medium for Rooting (SMR) were alternated at 4–6-week intervals. Plants were maintained at 22°C with 16h-photoperiod, 600mWm^-2^nm^-1^ throughout.

#### Genotyping of transgenic lines

DNA were extracted from 50-100mg of leaf tissue using an in-house protocol described subsequently. The frozen leaf tissues were ground with metal ball-bearing balls (IG100_5/32_PK1000; Simply Bearings Ltd., Leigh, UK) using a mechanical pulveriser (MiniG from Spex) at 1200 rpm for 30 seconds. 500µl of the extraction buffer (1.25% sodium dodecyl sulphate; 100 mM Tris HCl pH 8.0; 50 mM EDTA pH 8.0 and 25 mg PVP) were added to the disrupted tissue. The samples were mixed and incubated at 65°C for 30 min, inverting the tubes each 5 min. Samples were cooled on ice for 5 min and then 250 µl of chilled 5M NaCl added to the tube and incubated for15 min on ice. The samples were centrifuged for 10 min at 20,000 g. Supernatant was transferred into a new tube containing 360 µl isopropanol. Samples were vortexed and incubated for 30 minutes or overnight at –20°C for DNA precipitation. The samples were centrifuged for 20 min at 15,700 g. Supernatant was discarded and pellet was washed in 500 µl of 70% ethanol twice. DNA pellet was resuspended in 50 µl TE buffer (10 mM Tris HCl pH 8.0; 1 mM EDTA pH 8.0). PCR amplification was performed using gene specific primers and PCRBIO Taq Mix Red (PCR Biosystems) following the manufacturer’s guidelines.

### Stable transformation of barley (*Hordeum vulgare* L. cv. Golden Promise)

#### Donor plant growth

Barley seeds were treated with 70% ethanol for 2 minutes, followed with three times wash in sterile water. The seeds were then surfaced sterilized by 5% sodium hypochlorite solution for 4 minutes and rinsed with sterile water 4–5 times. The sterilized seeds were plated on 1% water agar plates and imbibed at 4 °C for 3 days and then germinated in the dark at 22 °C for 2–3 days^39^.

#### Barley transformation

Barley (*Hordeum vulgare* L. cv. Golden Promise) lines were raised using *A. tumifaciens* mediated transformation ^40,41^. Briefly, the constructs were transformed into *A. tumefaciens* strain AGL1. Immature barley seeds (embryo size ∼ 1.5mm in diameter) were harvested and sterilized with 70% ethanol for 2 minutes followed by 3 washes in sterile water. The seeds were then sterilized with 5% sodium hypochlorite solution for 4 minutes followed by rinsing in sterile water for 4-5 times. Immature embryos were isolated using fine forceps in a laminar flow hood and the embryonic axis removed from the scutellum and discarded. They were then cultured on Callus induction (CI) plates at 23-24°C in the dark overnight ^41^. The embryos were treated with *Agrobacterium* inoculum and transferred to new CI plates for co-cultivation. After 3 days, the embryos were transferred to CI plates with antibiotics: Timentin, to kill the *Agrobacterium* and hygromycin, the selection of transgenic cells. The tissue culture were maintained at 23-24°C in the dark for 2 weeks, embryos were transferred to fresh CI plates and the cultured under the same conditions for another 4 weeks to induce callus formation, with subcultures to fresh plates after 2 weeks. Calli were then transferred to Transition medium under low light for 2 weeks^41^. Calli undergoing active shooting were transferred to regeneration plates until small plantlets formed. Individual plantlets were separated and transferred to rooting tubes. Once the plantlets developed good root systems, they were transferred to soil for growth to maturity.

### Constructs production

The Golden Gate modular cloning system were used to prepare plasmids for the transformation experiments ^42^. All the Level 0, Level 1 and Level 2 are listed in Table S8. Sequences were domesticated, synthesized, and cloned into pMS (GeneArt, Thermo Fisher Scientific, Waltham, USA). Sequence information for *MtNIN* (*M. truncatula* Mt5.0v1 genome via Phytozome (https://phytozome.jgi.doe.gov)) and *DgNIN* and *DgNIN2* (*D. glomerata* (ASM325502v1)) and *DryasNIN* (*D. drummondii* (ASM325486v1) from NCBI^4^. For *DgNIN-RNAi* construct the fragment was amplified using forward primers 5’-ccgggattccgctatggtcagcaggttgatg-3’ with BamH1 overhang and reverse primer 5’-cgcctcgagcagaatgcttgtaacacccag-3’ with Xho1 overhang and cloned into entry vector *pENTR3c* dual (Invitrogen) and sequence confirmed. The entry clone was recombined into the destination vector *pK7GWIWG2-7F2.1* (VIB, Ghent) using LR clonase Gateway cloning kit (Invitrogen) (see Supplementary Table 4).

### Gene Expression analysis

RNA extraction from *D. glomerata* and *F. vesca*: Individual plant root system is harvested forming a biological replicate weighing 100-150mg. RNA was extracted using a modified extraction buffer containing 2%SDS (v/v), 100mM Tris-HCl pH 8.0, 1.4M NaCl, 20mM EDTA. The samples were crushed with tungsten-carbide beads (Qiagen) in a mechanical pulveriser (Genogrinder) in the presence of 12ul/ml β-mercaptoethanol (Sigma) and 50mg PVPP (Sigma). The powder is resuspended with the extraction buffer followed by treatment with chloroform:iso-amyl alcohol (v/v24:1) (Melford) and chloroform (Sigma). The RNA was precipitated using 7.5M Lithium Chloride solution. The precipitated RNA was washed with 75% ethanol followed by air drying and re-suspension in deionised water. The extracted RNA was quantified using Nanodrop (Thermo) and then treated for DNAse contamination using TurboDNAse free kit (Thermo) using the manufacturer protocol.

RNA extraction from *M. truncatula:* Approximately 100-150mg (fresh weight) tissue from individual plant root system was harvested for RNA extraction using MN Nucleospin Plants RNA extraction kit following the manufacturer guidelines. The eluted RNA was resuspended in deionised water and quantified using Nanodrop (Thermo).

RNA extraction from *H. vulgare:* Approximately 100-150mg (fresh weight) tissue from individual plant root system was harvested for RNA extraction using QIAGEN RNeasy plant RNA extraction kit following the manufacturer guidelines. The eluted RNA was resuspended in deionised water and quantified using Nanodrop (Thermo) and then treated for DNAse digestion using TurboDNAse free kit (Thermo) using the manufacturer’s protocol.

### Quantitative realtime-PCR

500ng of the treated RNA was then reverse transcribed to make cDNA using Superscript III (Invitrogen) using manufacturer’s protocol. Quantitative real-time polymerase chain reaction (qRT-PCR) were performed in technical replicates using SyGreen mix Lo-Rox (PCR Biosystems) using Quantiflex 7 system (ABI) in a total volume of 10ul. The primer pairs for gene expression analysis are listed in Supplementary Table 5.

### RNAseq analysis

For RNAseq analysis the integrity of the extracted DNAse treated RNA were checked using Agilent Tapestation following manufacturer protocol. Library preparation and paired-end RNA sequencing were performed by Novogene (Cambridge, UK) on an Illumina Novaseq 6000 or Novaseq Xplus platform, generating high-quality 150 bp paired-end reads. Raw sequencing data were first subjected to quality assessment and adapter trimming using TrimGalore (v0.6.10), which integrates Cutadapt and FastQC for removal of low-quality bases (Phred score < 20) and sequencing adapters^43^. Only reads passing the default quality filters were retained for downstream analyses. Following trimming, transcript abundance was estimated using Kallisto (v0.50.1), a pseudo-alignment-based quantification tool that enables rapid and accurate transcript-level abundance estimation^44^. A reference transcriptome index was built from the [species name/version] reference transcriptome (*F. vesca* v4.0.a1; *H. vulgare* v1r1_Apollo) with concatenated target sequence (*DryasNIN*) using a k-mer size of 21. Multiple paired end read files per sample were automatically merged prior to quantification, and transcript abundance was reported in transcripts per million (TPM) and estimated counts for each transcript. Transcript-level count matrices generated by Kallisto were imported into R (v4.5.1) using the Tximport package to summarize abundances at the gene level^45^. Differential expression analysis was performed with DESeq2 (v3.17)^46^. Genes with low expression (total counts <10 across all samples) were filtered out prior to statistical testing. Normalization was carried out using DESeq2’s median-of-ratios method, and differential expression between experimental conditions was determined using the Wald test. Genes with an adjusted p-value < 0.05 and absolute log₂ fold-change ≥ 1 were considered significantly differentially expressed. All computational analyses were performed in RStudio. Visualization of differential expression and quality metrics, including PCA plots, volcano plots, and heatmaps, were generated using ggplot2, ComplexUpset and ComplexHeatmap packages^47–49^.

### Remapping of *M. truncatula* RNAse<u>q</u> datasets

RNA-seq datasets from *Medicago truncatula*, including independent time-course datasets of rhizobial spot inoculation and lateral root development, as well as NIN overexpression data, were re-analysed following remapping to the *M. truncatula* genome version 5^50^. The time-course datasets and the NIN overexpression dataset were generated and processed in separate previous studies; experimental details are described in the corresponding publications for timecourse nodule and lateral root (GEO accession GSE133612) and NIN overexpression (GEO accession GSE178119)^9,10,14^. Raw FASTQ files were quality assessed using FastQC and aligned to the *M. truncatula* v5 reference genome using STAR v2.7.11b, generating coordinate-sorted BAM files. Gene-level counts were calculated from aligned reads and imported into R for downstream analysis. Data processing and statistical analyses were performed in R using DESeq2 for normalization and differential expression modelling. At least three biological replicates were included for each condition. Variance-stabilised expression values were used for principal component analysis to assess sample clustering. Lowly expressed genes were filtered prior to differential expression analysis using counts per million (CPM) values calculated with edgeR^51^, together with a minimum raw count threshold (CPM < 0.35 and counts < 10 removed). Differentially expressed genes (DEGs) were identified by pairwise comparisons between experimental and control conditions using DESeq2, with a threshold of false discovery rate (FDR) < 0.05 and absolute log_2_ fold change ≥ 1.

### Histochemical assays and staining

For GUS staining the roots were thoroughly cleaned in sterile water and assessed under the stereo-fluorescent microscope so that only the transformed roots were harvested for the staining purpose. The roots were fixed in 90% acetone on ice for 1h. Subsequently, the acetone is replaced with 50mM phosphate buffer pH7.2 (Sigma) and washed thoroughly to remove any trace of acetone. The roots were placed in staining solution (0.1M Phosphate buffer, 0.25mM K_3_Fe(CN)_6_ (potassium ferricyanide), 0.25 mM K_4_Fe(CN)_6_ (potassium ferrocyanide), 0.25%(v/v) Triton X-100 and 1mM 5-bromo-4-chloro-3-indolyl-beta-D-glucuronide (X-Gluc, Thermo)). The samples were vacuum infiltrated for 30mins and incubated at 37°C untill the blue staining was visible. After the staining the tissues were cleared and stored in 50% ethanol for imaging under the stereomicroscope DX105 F (Leica). For ruthenium red staining, a few milcrolitres of (0.1% (w/v) ruthenium red and pippeted over each tissue section and cleared using sterile deionised water. For toluidine blue staining, a few milcrolitres of (0.1% (w/v) toluidine blue staining solution was pippeted over each section and cleared using sterile deionized water.

### Histology

Roots and nodules were fixed with 2.5% glutaraldehyde (Sigma, G5882) in 1xPBS overnight. The fixed material was dehydrated in an ethanol series and subsequently embedded in Technovit 7100 (Kulzer Technik, Wehrheim, Germany) according to the manufacturer’s protocol. Embedded tissues were sectioned (10-15μm) using a Leica microtome (Histocore autocut R) and mounted on glass slide followed by staining with ruthenium red solution or toluidine blue staining solution. Imaged using stereomicroscope DX105 F (Leica) using LAS 3.4 software.

### Microscopy

*M. truncatula* hairy-root: To quantify the total number of curled root hair, extended root hairs, infection patches, each transgenic root was considered as an independent system, and all the structures were scored in random sections of ∼1cm as previously described ^7^. Nodules were counted in all the transformed roots and imaged by stereomicroscope DX105 F (Leica) using LAS 3.4 software and DFC7000T (Leica) using LAS X software.

*D. glomerata* hairy-root: Whole plant fluorescent images were captured using ImageQuant 800 system (GE Healthcare, Chicago, IL, USA) using proprietary software. The higher magnification images are captured using stereomicroscope DX105 F (Leica) using LAS 3.4 software.

*F. vesca* stable lines: Images captured using stereomicroscope DX105 F (Leica) using LAS 3.4 software.

*H. vulgare* stable lines: Whole plant fluorescent images were captured using ImageQuant800 system (GE Healthcare, Chicago, IL, USA) using proprietary software. The higher magnification images are captured using stereomicroscope DX105 F (Leica) using LAS 3.4 software.

### Hormone treatment

*D. glomerata:* Seeds were germinated on ½ MS plates supplemented with 0.5gm/L MES (Merck), 0.8% agarose (Sigma) and adjusted to pH 5.8. After 4 weeks, seedlings were transferred to BNM –N+P plates supplemented with NAA and maintained for the desired time point under 16h-photperiod, 600mWm^-2^nm^-1^ at 22°C before harvesting for RNA extraction and GUS staining.

*M. truncatula:* Seeds were germinated on water agar plates and grown for 4 days before moving them to BNM –N+P plates supplemented with NAA and maintained for the desired time point under 16h-photoperiod, 600mWm^-2^nm^-1^ at 22°C before harvesting for RNA extraction and GUS staining.

*F. vesca:* Tissue culture plants growing in SMR (approx. 4 weeks after transfer) were moved to ½ Hoagland’s media plates supplemented with 100µM NAA, 0.8% agar (Sigma). The plants were maintained for 4 weeks under 16h-photoperiod, 600mWm^-2^nm^-1^ at 24-22°C before harvesting them for phenotyping and RNA extraction.

*H. vulgare:* Germinated seedlings (7 days after germination) were moved to ½ Hoagland’s media plates supplemented with 50µM NAA, 0.8% agarose (Sigma). The plants were maintained for 3 weeks under 16h-photoperiod, 600mWm^-2^nm^-1^ at 24-22°C before harvesting them for phenotyping and RNA extraction.

### Promoter analysis

Promoter region of variable length ranging from 2000bp to 5000bp upstream from the transcriptional start site (TSS) were extracted for the individual NLP gene (Supplementary Table 4) and scanned for the *AuxRE* motif TGTCNN and NNGACA. A position specific background model with nucleotide frequency A = 0.275, C = 0.225, G = 0.225, T = 0.275 was used. Motif occurrences were identified using FIMO (MEME Suite v5.5.7) with a p-value cutoff of 0.01 ^52^. Data visualizations were performed in RStudio (v4.5.1) using ggridges (v0.5.7) and ggseqlogo (v0.2) packages.

### Genomic DNA Extraction and Sequencing

High molecular weight (HMW) genomic DNA was isolated from leaf tissue harvested from a single plant, following a specialised HMW extraction protocol^53^. For long-read sequencing, libraries were constructed from approximately 1μg HMW DNA using the Oxford Nanopore Technologies (ONT) SQK-LSK108 Ligation Sequencing Kit, in accordance with the manufacturer’s instructions. These libraries were sequenced on an ONT GridION X5 platform using R9.4.1 flow cells (FLO-MIN106), with base calling performed using high accuracy settings. Short-read sequencing was outsourced to Novogene (Cambridge, UK); this involved the generation of 150 bp paired-end reads on an Illumina HiSeq 2500 system.

### Genome Assembly and Polishing

Genome size estimations were determined using the K-mer Analysis Toolkit (KAT)^54^. Long-read quality was evaluated with NanoPlot v1.30.1^55^, and adapter sequences were removed using Porechop v0.2.4 (https://github.com/rrwick/Porechop) with default parameters. To refine the dataset, reads were filtered via Filtlong v0.2.1 (https://github.com/rrwick/Filtlong) to exclude sequences shorter than 1 kb or those with a quality score below Q7. The *de novo* assembly of long reads was performed using NECAT v0.0.1_update20200803 ^56^, incorporating the estimated genome sizes while maintaining all other default settings. Following the primary assembly, Purge_Dups v1.0.1 was used to identify and remove redundant heterozygous contigs and overlapping sequences^57^. The assembly then underwent error correction; long reads were aligned to the contigs using Minimap2 v2.17-r941 ^58^ to inform one round of Racon v1.4.20^59^, followed by a single iteration of Medaka v1.5.0 (https://github.com/nanoporetech/medaka) using the r941_min_high_g360 model. For short-read processing, Illumina data quality was verified with FastQC v0.11.9 (https://www.bioinformatics.babraham.ac.uk/projects/fastqc/), and Trimmomatic v0.39^60^ was used to prune adapters and low-quality regions. These processed short reads were mapped to the corrected long-read assembly using Bowtie2 v2.4.4^61^, with alignment sorting and indexing performed with Samtools v1.12^62^. This alignment facilitated three successive rounds of polishing with Pilon v1.24^63^. Final assembly statistics were calculated using a custom Python script, and assembly completeness was quantified via BUSCO v5.2.2 using the eudicots_odb10 database^64^.

### RNA Isolation and Sequencing for genome annotation

When available, tissue samples were collected from the leaves, stems, and roots. These samples were immediately snap frozen in liquid nitrogen and maintained at –80°C until required. Total RNA was isolated from inoculated root material using a methodology adapted from Yu et al.^65^. Briefly, tissues were powdered under liquid nitrogen using a mortar and pestle that had been decontaminated with RNaseZap and baked for 2 hours at 230°C to eliminate RNase activity. The resulting powder was transferred to a pre-warmed extraction buffer containing β-mercaptoethanol and 1% w/w PVPP. The protocol then followed the stages described by Yu et al.^65^, with the exception that RNA was eluted in 60 μL of DEPC-treated water. RNA integrity was assessed, and only samples with a RIN value exceeding 7.0 were submitted to Novogene (Cambridge, UK) for 150 bp paired-end sequencing on the Illumina HiSeq 2500 platform.

### Genome Annotation

Repetitive and low complexity genomic regions were identified and soft-masked using Red (version 05/22/2015)^66^. Quality control for RNA-seq data was performed with FastQC v0.11.9, followed by adapter and quality trimming using Trimmomatic v0.39^60^. The assemblies were indexed, and the trimmed RNA-seq reads were aligned using HISAT2 v2.2.1 under default setting ^67^. Gene prediction was conducted with BRAKER2 v2.1.6 in –etpmode, using both the RNA-seq alignments and the eudicot protein database as evidence^68^. Finally, annotation completeness was evaluated using BUSCO v5.2.2 against the eudicots_odb10 database^64^.

### Orthogroup analysis

The hierarchical orthologous groups (HOGs) annotating the ancestral nodes used for the correlation analysis is based on previous study^26^. The orthogroups reconstruction was done using 50 genomes ^4,5,50,69–88^, including newly sequenced and annotated genomes for 11 nodulators (*Allocasuarina nana*, *Ceanothus thyrsiflorus*, *Cercocarpus montanus*, *Colletia paradoxa*, *Colletia ulicina*, *Coriaria japonica*, *Datisca cannabina*, *Elaeagnus angustifolia*, *Hippophae salicifolia*, *Myrica cerifera* and *Shepherdia argentea*), 2 re-sequenced nodulator (*Datisca glomerata* and *Dryas drummondii*) and 7 non-nodulator (*Cornycarpus laevigatus*, *Dryas octopetala*, *Filipendula ulmaria*, *Pomaderris vaccinifolia*, *Purshia tridentata*, *Rubus ellipticus*, *Sagretia theezans*) (table S3) with OrthoFinder v3.1.0 using the ultra-sensitive Diamond mode (–S diamond_ultra_sens option)^89^. The inferred tree from OrthoFinder was then checked for consistency with known species trees. The HOGs for *M. truncatula*, *D. glomerata* v3, *F. vesca* and *H. vulgare* were inferred from the tree.

### Orthogroup based inter-species gene expression matrix construction

As Orthogroups differ in gene number between the species, normalization were performed at the orthogroup level by calculating the median log2 fold-change in gene expression for each species. Median aggregation was chosen to minimize the influence of outlier expression values ensuring comparable expression summaries which was converted to categorical expression status namely Up (>0), Down (<0) and No Change (0). The data were plotted using ggalluvial (v0.12.5) package in R^90^, where flow represent each orthogroup status. Polychoric correlations were computed to estimate the association between ordered categorical expression states (Up, No Change, Down) across the three species. Pairwise polychoric correlation coefficients calculated from the logical annotations derived from median log₂ fold-change values within each orthogroup using Polycor (v0.8.2) package and bootstrap calculated using the boot (v1.3.32) to extract the asymptotic p-values in R. Positive correlations indicate concordant directional expression patterns between species, whereas negative correlations reflect divergent transcriptional responses.

### Nicotiana benthamiana transient assay

For *Nicotiana benthamiana* leaf transient assays, *Agrobacterium tumefaciens* strain EHA105 carrying various constructs, were mixed with *A. tumefaciens* EHA105 strain carrying *p19* before infiltration, as described ^91^. Leaves were harvested for analysis three days post infiltration. Images of the whole leaves were acquired under bright filed, *mCherry*, UV channels by ImageQuant 800 system (GE Healthcare, Chicago, IL, USA). The images were analysed using ImageJ for assessing the infiltrated area (mCherry channel) and senescent area (UV channel).

### Dual luciferase assay

The dual luciferase assay was performed as previously described ^92^. *N. benthamiana* leaf discs were harvested 2 days post transfection and ground into a fine powder in 2 mL microcentrifuge tubes. The samples were incubated with 200 µL of Passive Lysis Buffer (E1910, Promega, Madison, WI, USA). The crude leaf extract was centrifuged at 15,000×g for 2 min at room temperature. An aliquot of the supernatant was used for the assay following the manufacturer’s protocol (E1910, Promega, Madison, WI, USA). Chemiluminescence was measured using a plate reader (CLARIOstar, BMG Labtech, Ortenberg, Germany). For each reporter construct, the promoter of interest was cloned into the pGreenII-0800 vector via recombination reactions (E2611S, NEB, Ipswich, MA, USA) to fuse with the Firefly luciferase gene, while the constitutively expressed Renilla luciferase from the same vector was used for normalization. The LUC/REN ratio was calculated and plotted using RStudio (v4.5.1). At least four biological replicates from two independent experiments were recorded.

### Phylogenetic tree analysis

Phylogenetic analysis were conducted using IQ-TREE2 where maximum likelihood trees were inferred using the substitution model Q.mammal+F+R5. Branch support was assessed using 1000 replicates of the SH-like appropriate likelihood ratio test (SH-aLRT) and 1000 ultrafast bootstrap replicates^93^. Default parameters were used for all other settings. The resulting phylogenetic trees were visualized and annotated using iTOL^94^.

### Statistical analysis

Statistical analyses were done using RStudio (v4.5.1). Test applied for each plot are stated in the respective figure legends.

## Notes

### Competing Interest Statement

The authors have declared no competing interest.

## REFERENCE

1. Kundu, A., Moraes, T. A., Price, R. J., Harrison, R. J. & Oldroyd, G. E. D. Getting to the Route: The Evolution of Nitrogen-Fixing Nodules. Annual Review of Cell and Developmental Biology Downloaded from www.annualreviews.org. Cambridge University 98, (2025).

2. Kates, H. R. et al. Shifts in evolutionary lability underlie independent gains and losses of root-nodule symbiosis in a single clade of plants. Nat. Commun. 15, 4262 (2024).

3. Shen, D. et al. A homeotic mutation changes legume nodule ontogeny into actinorhizal-type ontogeny. Plant Cell 32, 1868–1885 (2020).

4. Griesmann, M. et al. Phylogenomics reveals multiple losses of nitrogen-fixing root nodule symbiosis. Science (1979). 361, (2018).

5. van Velzen, R. et al. Comparative genomics of the nonlegume *Parasponia* reveals insights into evolution of nitrogen-fixing rhizobium symbioses. Proceedings of the National Academy of Sciences 115, (2018).

6. Soltis, D. E. et al. Chloroplast gene sequence data suggest a single origin of the predisposition for symbiotic nitrogen fixation in angiosperms. Proceedings of the National Academy of Sciences 92, 2647–2651 (1995).

7. Liu, J. et al. A remote cis-regulatory region is required for nin expression in the pericycle to initiate nodule primordium formation in medicago truncatula. Plant Cell 31, 68–83 (2019).

8. Soyano, T., Shimoda, Y., Kawaguchi, M. & Hayashi, M. A shared gene drives lateral root development and root nodule symbiosis pathways in Lotus. Science (1979). 366, 1021–1023 (2019).

9. Schiessl, K. et al. NODULE INCEPTION Recruits the Lateral Root Developmental Program for Symbiotic Nodule Organogenesis in Medicago truncatula. Current Biology 29, 3657–3668.e5 (2019).

10. Lee, T. et al. Light-sensitive short hypocotyl genes confer symbiotic nodule identity in the legume Medicago truncatula. Current Biology 34, 825–840.e7 (2024).

11. Pawlowski, K. & Demchenko, K. N. The diversity of actinorhizal symbiosis. Protoplasma 249, 967–979 (2012).

12. Clavijo, F. et al. The Casuarina NIN gene is transcriptionally activated throughout Frankia root infection as well as in response to bacterial diffusible signals. New Phytologist 208, 887–903 (2015).

13. Vernié, T. et al. The NIN transcription factor coordinates diverse nodulation programs in different tissues of the medicago truncatula root. Plant Cell 27, 3410–3424 (2015).

14. Feng, J., Lee, T., Schiessl, K. & Oldroyd, G. E. D. Processing of NODULE INCEPTION controls the transition to nitrogen fixation in root nodules. Science (1979). 374, 629–632 (2021).

15. Singh, S., Katzer, K., Lambert, J., Cerri, M. & Parniske, M. CYCLOPS, A DNA-binding transcriptional activator, orchestrates symbiotic root nodule development. Cell Host Microbe 15, 139–152 (2014).

16. Liu, J. & Bisseling, T. Evolution of NIN and NIN-like Genes in Relation to Nodule Symbiosis. Genes (Basel). 11, 777 (2020).

17. Gauthier-Coles, C., White, R. G. & Mathesius, U. Nodulating legumes are distinguished by a sensitivity to cytokinin in the root cortex leading to pseudonodule development. Front. Plant Sci. 9, (2019).

18. Galli, M. et al. The DNA binding landscape of the maize AUXIN RESPONSE FACTOR family. Nat. Commun. 9, (2018).

19. Rightmyer, A. P. & Long, S. R. Pseudonodule Formation by Wild-Type and Symbiotic Mutant Medicago truncatula in Response to Auxin Transport Inhibitors. Mol. Plant. Microbe. Interact. 24, 1372–1384 (2011).

20. Zhang, Y. et al. Comparative phylogenomics and phylotranscriptomics provide insights into the genetic complexity of nitrogen-fixing root-nodule symbiosis. Plant Commun. 5, (2024).

21. Svistoonoff, S. et al. The Independent Acquisition of Plant Root Nitrogen-Fixing Symbiosis in Fabids Recruited the Same Genetic Pathway for Nodule Organogenesis. PLoS One 8, e64515 (2013).

22. Roy, S. et al. Celebrating 20 Years of Genetic Discoveries in Legume Nodulation and Symbiotic Nitrogen Fixation. Plant Cell 32, 15–41 (2020).

23. Rübsam, H. et al. Nanobody-driven signaling reveals the core receptor complex in root nodule symbiosis. Science (1979). 379, 272–277 (2023).

24. Kim, S. & Wysocka, J. Deciphering the multi-scale, quantitative cis-regulatory code. Molecular Cell vol. 83 373–392 Preprint at 10.1016/j.molcel.2022.12.032 (2023).

25. Op den Camp, R. et al. LysM-Type Mycorrhizal Receptor Recruited for Rhizobium Symbiosis in Nonlegume Parasponia. Science (1979). 331, 909–912 (2011).

26. Libourel, C. et al. Comparative phylotranscriptomics reveals ancestral and derived root nodule symbiosis programmes. Nat. Plants 9, 1067–1080 (2023).

27. Liu, H. et al. Evolution of root nodule symbiosis via paleopolyploidy and modular pathway rewiring. Cell Host Microbe https://doi.org/10.1016/j.chom.2026.01.001 (2026) doi:10.1016/j.chom.2026.01.001.

28. Qiao, L. et al. Nanodomain-localized formin gates symbiotic microbial entry in legume and solanaceous plants. Science (1979). 391, 1036–1045 (2026).

29. Doyle, J. J., Ren, J., Pawlowski, K., James, E. K. & Gao, Y. One versus many independent assemblies of symbiotic nitrogen fixation in flowering plants. Nature Communications 16, (2025).

30. van Velzen, R., Doyle, J. J. & Geurts, R. A Resurrected Scenario: Single Gain and Massive Loss of Nitrogen-Fixing Nodulation. Trends Plant Sci. 24, 49–57 (2019).

31. Cavallari, N., Artner, C. & Benkova, E. Auxin-regulated lateral root organogenesis. Cold Spring Harb. Perspect. Biol. 13, (2021).

32. Albalat, R. & Cañestro, C. Evolution by gene loss. Nature Reviews Genetics vol. 17 379–391 Preprint at 10.1038/nrg.2016.39 (2016).

## MATERIALS AND METHODS REFERENCES

33. Persson, T. et al. Candidatus Frankia datiscae Dg1, the Actinobacterial microsymbiont of datisca glomerata, expresses the canonical nod genes NodABC in symbiosis with its host plant. PLoS One 10, (2015).

34. Marsh, J. F. et al. Medicago truncatula NIN is essential for rhizobial-independent nodule organogenesis induced by autoactive calcium/calmodulin-dependent protein kinase. Plant Physiol. 144, 324–335 (2007).

35. Plet, J. et al. MtCRE1-dependent cytokinin signaling integrates bacterial and plant cues to coordinate symbiotic nodule organogenesis in Medicago truncatula. Plant Journal 65, 622–633 (2011).

36. Boisson-Dernier, A. et al. *Agrobacterium rhizogenes* –Transformed Roots of *Medicago truncatula* for the Study of Nitrogen-Fixing and Endomycorrhizal Symbiotic Associations. Molecular Plant-Microbe Interactions® 14, 695–700 (2001).

37. Ehrhardt, D., Atkinson, E. & Long. Depolarization of alfalfa root hair membrane potential by Rhizobium meliloti Nod factors. Science (1979). 256, 998–1000 (1992).

38. Sanchez, E. R. et al. Overexpression of Vitis GRF4-GIF1 improves regeneration efficiency in diploid Fragaria vesca Hawaii 4. Plant Methods 20, (2024).

39. Li, X. R. et al. Nutrient regulation of lipochitooligosaccharide recognition in plants via NSP1 and NSP2. Nat. Commun. 13, (2022).

40. Bartlett, J. G., Alves, S. C., Smedley, M., Snape, J. W. & Harwood, W. A. High-throughput Agrobacterium-mediated barley transformation. Plant Methods 4, (2008).

41. Bartlett, J. G., Snape, J. W. & Harwood, W. A. Intron-mediated enhancement as a method for increasing transgene expression levels in barley. Plant Biotechnol. J. 7, 856–866 (2009).

42. Weber, E., Engler, C., Gruetzner, R., Werner, S. & Marillonnet, S. A modular cloning system for standardized assembly of multigene constructs. PLoS One 6, (2011).

43. Martin, M. Cutadapt removes adapter sequences from high-throughput sequencing reads. EMBnet. J. 17, 10 (2011).

44. Bray, N. L., Pimentel, H., Melsted, P. & Pachter, L. Near-optimal probabilistic RNA-seq quantification. Nat. Biotechnol. 34, 525–527 (2016).

45. Soneson, C., Love, M. I. & Robinson, M. D. Differential analyses for RNA-seq: transcript-level estimates improve gene-level inferences. F1000Res. 4, 1521 (2015).

46. Love, M. I., Huber, W. & Anders, S. Moderated estimation of fold change and dispersion for RNA-seq data with DESeq2. Genome Biol. 15, (2014).

47. Gu, Z., Eils, R. & Schlesner, M. Complex heatmaps reveal patterns and correlations in multidimensional genomic data. Bioinformatics 32, 2847–2849 (2016).

48. Lex, A., Gehlenborg, N., Strobelt, H., Vuillemot, R. & Pfister, H. UpSet: Visualization of intersecting sets. IEEE Trans. Vis. Comput. Graph. 20, 1983–1992 (2014).

49. Krassowski, M., Arts, M., Lagger, C. & Max. Complex-upset: v1.3.5. Preprint at 10.5281/zenodo.3700590 (2022).

50. Pecrix, Y. et al. Whole-genome landscape of Medicago truncatula symbiotic genes. Nature Plants vol. 4 1017–1025 Preprint at 10.1038/s41477-018-0286-7 (2018).

51. Robinson, M. D., McCarthy, D. J. & Smyth, G. K. edgeR: A Bioconductor package for differential expression analysis of digital gene expression data. Bioinformatics 26, 139–140 (2009).

52. Grant, C. E., Bailey, T. L. & Noble, W. S. FIMO: Scanning for occurrences of a given motif. Bioinformatics 27, 1017–1018 (2011).

53. Schalamun, M. et al. Harnessing the MinION: An example of how to establish long-read sequencing in a laboratory using challenging plant tissue from Eucalyptus pauciflora. Mol. Ecol. Resour. 19, 77–89 (2019).

54. Mapleson, D., Accinelli, G. G., Kettleborough, G., Wright, J. & Clavijo, B. J. KAT: A K-mer analysis toolkit to quality control NGS datasets and genome assemblies. Bioinformatics 33, 574–576 (2017).

55. De Coster, W., D’Hert, S., Schultz, D. T., Cruts, M. & Van Broeckhoven, C. NanoPack: Visualizing and processing long-read sequencing data. Bioinformatics 34, 2666–2669 (2018).

56. Chen, Y. et al. Efficient assembly of nanopore reads via highly accurate and intact error correction. Nat. Commun. 12, (2021).

57. Guan, D. et al. Identifying and removing haplotypic duplication in primary genome assemblies. Bioinformatics 36, 2896–2898 (2020).

58. Li, H. Minimap2: Pairwise alignment for nucleotide sequences. Bioinformatics 34, 3094–3100 (2018).

59. Vaser, R., Soviæ, I., Nagarajan, N. & Šikiæ, M. Fast and accurate de novo genome assembly from long uncorrected reads. Genome Res. 27, 737–746 (2017).

60. Bolger, A. M., Lohse, M. & Usadel, B. Trimmomatic: A flexible trimmer for Illumina sequence data. Bioinformatics 30, 2114–2120 (2014).

61. Langmead, B. & Salzberg, S. L. Fast gapped-read alignment with Bowtie 2. Nat. Methods 9, 357–359 (2012).

62. Li, H. et al. The Sequence Alignment/Map format and SAMtools. Bioinformatics 25, 2078–2079 (2009).

63. Walker, B. J. et al. Pilon: An integrated tool for comprehensive microbial variant detection and genome assembly improvement. PLoS One 9, (2014).

64. Simão, F. A., Waterhouse, R. M., Ioannidis, P., Kriventseva, E. V. & Zdobnov, E. M. BUSCO: Assessing genome assembly and annotation completeness with single-copy orthologs. Bioinformatics 31, 3210–3212 (2015).

65. Yu, D. et al. Comparison and Improvement of Different Methods of RNA Isolation from Strawberry (Fragria * ananassa). Journal of Agricultural Science 4, (2012).

66. Girgis, H. Z. Red: An intelligent, rapid, accurate tool for detecting repeats de-novo on the genomic scale. BMC Bioinformatics 16, (2015).

67. Kim, D., Paggi, J. M., Park, C., Bennett, C. & Salzberg, S. L. Graph-based genome alignment and genotyping with HISAT2 and HISAT-genotype. Nat. Biotechnol. 37, 907–915 (2019).

68. Brùna, T., Hoff, K. J., Lomsadze, A., Stanke, M. & Borodovsky, M. BRAKER2: Automatic eukaryotic genome annotation with GeneMark-EP+ and AUGUSTUS supported by a protein database. NAR Genom. Bioinform. 3, 1–11 (2021).

69. Buti, M. et al. The genome sequence and transcriptome of Potentilla micrantha and their comparison to Fragaria vesca (the woodland strawberry). Gigascience 7, 1–14 (2018).

70. Chagné, D. et al. The draft genome sequence of European pear (Pyrus communis L. ‘Bartlett’). PLoS One 9, (2014).

71. VanBuren, R. et al. The genome of black raspberry (Rubus occidentalis). Plant J. 87, 535–547 (2016).

72. Edger, P. P. et al. Single-molecule sequencing and optical mapping yields an improved genome of woodland strawberry (Fragaria vesca) with chromosome-scale contiguity. GigaScience vol. 7 Preprint at 10.1093/gigascience/gix124 (2018).

73. Edger, P. P. et al. Origin and evolution of the octoploid strawberry genome. Nat. Genet. 51, 541–547 (2019).

74. Lamesch, P. et al. The Arabidopsis Information Resource (TAIR): Improved gene annotation and new tools. Nucleic Acids Res. 40, (2012).

75. Li, Q. et al. A chromosome-scale genome assembly of cucumber (Cucumis sativus L.). Gigascience 8, (2019).

76. Li, H., Jiang, F., Wu, P., Wang, K. & Cao, Y. A high-quality genome sequence of model legume lotus japonicus (Mg-20) provides insights into the evolution of root nodule symbiosis. Genes (Basel). 11, (2020).

77. Liu, M. J. et al. The complex jujube genome provides insights into fruit tree biology. Nat. Commun. 5, (2014).

78. Raymond, O. et al. The Rosa genome provides new insights into the domestication of modern roses. Nat. Genet. 50, 772–777 (2018).

79. Schreiber, M. et al. A genome assembly of the barley ‘transformation reference’ cultivar golden promise. G3: Genes, Genomes, Genetics 10, 1823–1827 (2020).

80. Sun, H. et al. Karyotype Stability and Unbiased Fractionation in the Paleo-Allotetraploid Cucurbita Genomes. Mol. Plant 10, 1293–1306 (2017).

81. Tang, H. et al. An improved genome release (version Mt4.0) for the model legume Medicago truncatula. BMC Genomics 15, 312 (2014).

82. Valliyodan, B. et al. Construction and comparison of three reference-quality genome assemblies for soybean. Plant Journal 100, 1066–1082 (2019).

83. Verde, I. et al. The Peach v2.0 release: High-resolution linkage mapping and deep resequencing improve chromosome-scale assembly and contiguity. BMC Genomics 18, (2017).

84. Wang, J. et al. Construction of pseudomolecules for the chinese chestnut (Castanea mollissima) genome. G3: Genes, Genomes, Genetics 10, 3565–3574 (2020).

85. Wu, Z. et al. Genome of Hippophae rhamnoides provides insights into a conserved molecular mechanism in actinorhizal and rhizobial symbioses. New Phytologist 235, 276–291 (2022).

86. Li, Y., Pi, M., Gao, Q., Liu, Z. & Kang, C. Updated annotation of the wild strawberry Fragaria vesca V4 genome. Hortic. Res. 6, (2019).

87. Young, N. D. et al. The Medicago genome provides insight into the evolution of rhizobial symbioses. Nature 480, 520–524 (2011).

88. Zhang, Q. et al. The genome of Prunus mume. Nat. Commun. 3, (2012).

89. Emms, D. M. & Kelly, S. OrthoFinder: Phylogenetic orthology inference for comparative genomics. Genome Biol. 20, (2019).

90. Rosvall, M. & Bergstrom, C. T. Mapping change in large networks. PLoS One 5, (2010).

91. Geddes, B. A. et al. Engineering transkingdom signalling in plants to control gene expression in rhizosphere bacteria. Nat. Commun. 10, (2019).

92. Gao, J. P. et al. An NSP2-MYB module orchestrates flavonoid biosynthesis and nodule symbiosis. Current Biology https://doi.org/10.1016/j.cub.2026.01.013 (2026) doi:10.1016/j.cub.2026.01.013.

93. Nguyen, L.-T., Schmidt, H. A., von Haeseler, A. & Minh, B. Q. IQ-TREE: A Fast and Effective Stochastic Algorithm for Estimating Maximum-Likelihood Phylogenies. Mol. Biol. Evol. 32, 268–274 (2015).

94. Letunic, I. & Bork, P. Interactive Tree of Life (iTOL) v6: recent updates to the phylogenetic tree display and annotation tool. Nucleic Acids Res. 52, W78–W82 (2024).

