## Supplementary Figures and Tables for "The demonstration of a single origin for nodule evolution, with nodule engineering in a non-nodulating species"

**The file includes:**

Supplementary Fig. 1-23
Supplementary Table 1-5
Supplementary Dataset 1-6

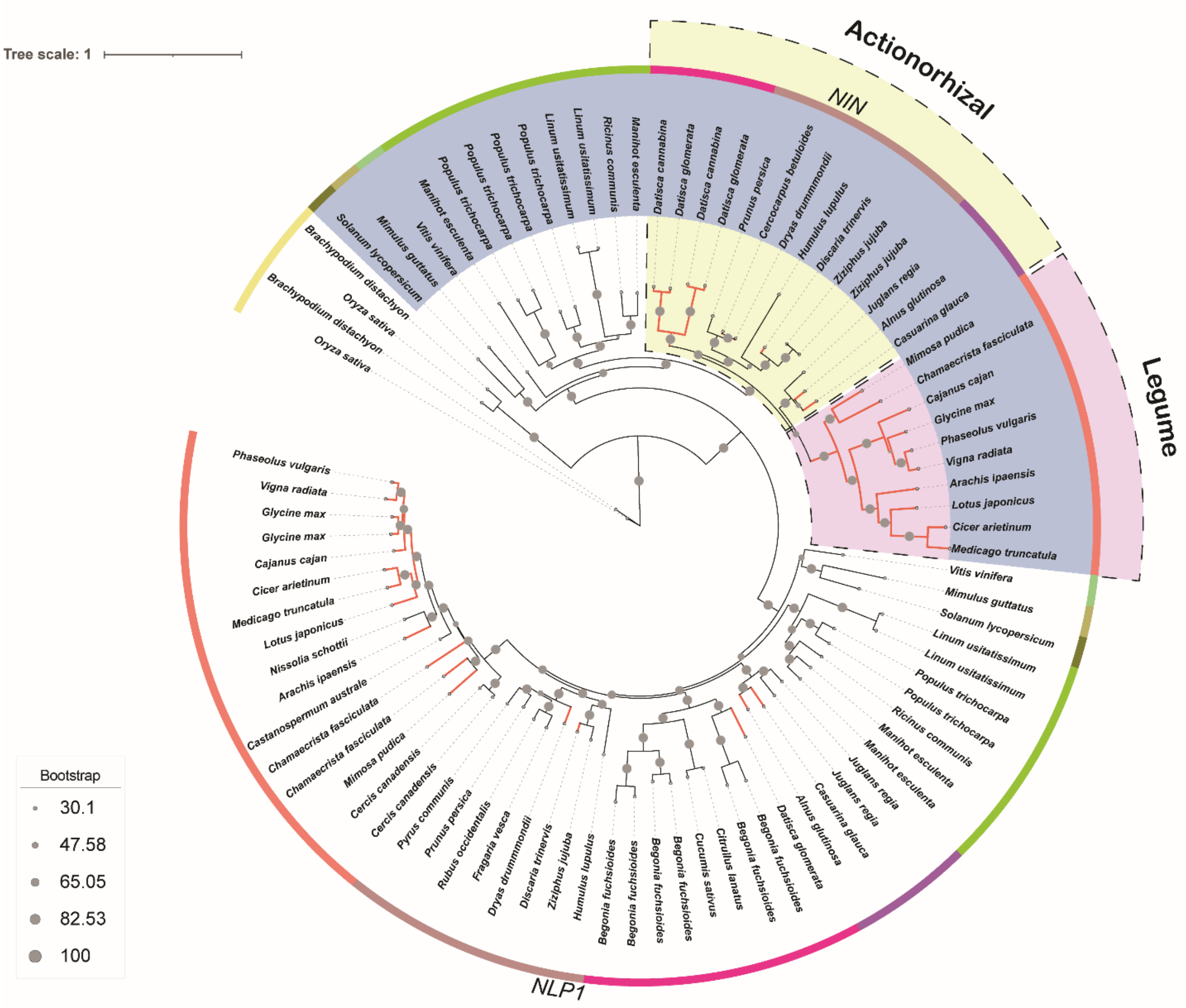


**Supplementary Fig. 1| Phylogenetic tree of the NLP1 clade.** Maximum-likelihood phylogenetic tree inferred with IQ-TREE2 based on the amino acid sequences of the NLP1 and NIN from within and outside of the NFC (table S1, data S1). The tree is created using Q.mammal+F+R5 substitution model. The branch support was assessed with 1000 bootstrap replicates. The tree is visualized using iTOL where NIN clade (steel grey highlight), legumes (orange), actinorhizal clades (Cucurbitales, purple; Rosales, brown; Fagales, violet), outside of NFC (shades of green) and nodulators (red branch).

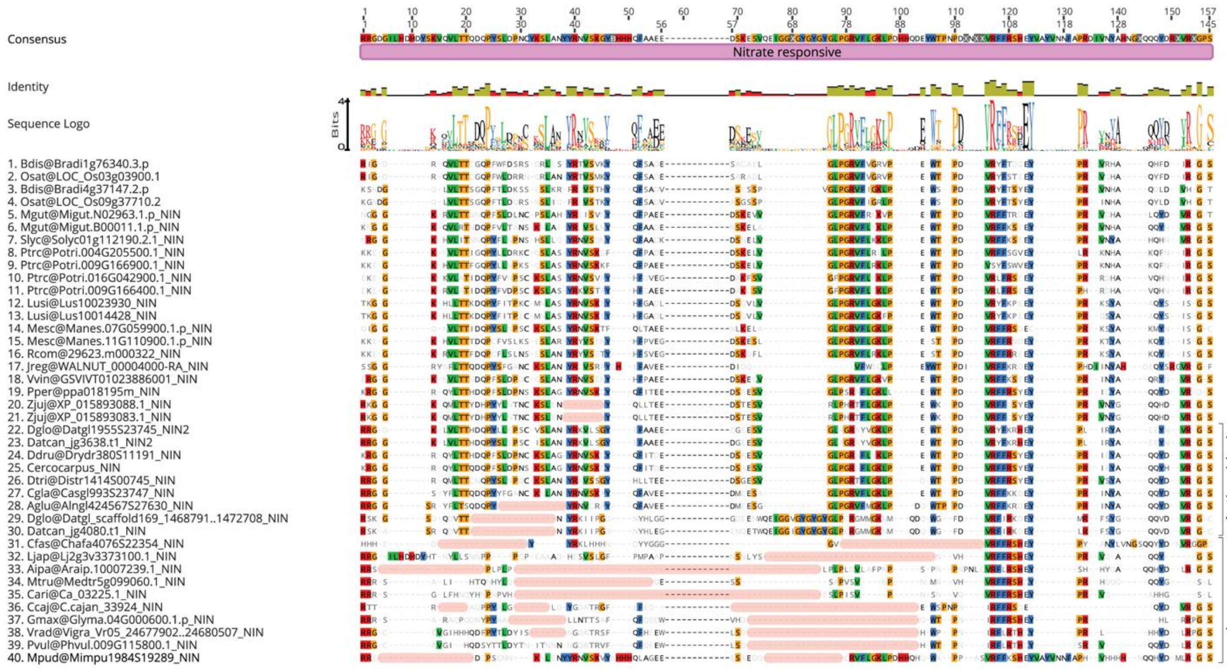


**Supplementary Fig. 2| Alignment of the NIN protein sequence showing the N-terminal Nitrate responsive domain.** The amino acid that matches the consensus are highlighted and the non-consensus are blurred. Sequence-logo represent the consensus sequence of amino acid residues. Pink bars indicate the deletions. See Supplementary Dataset 1.

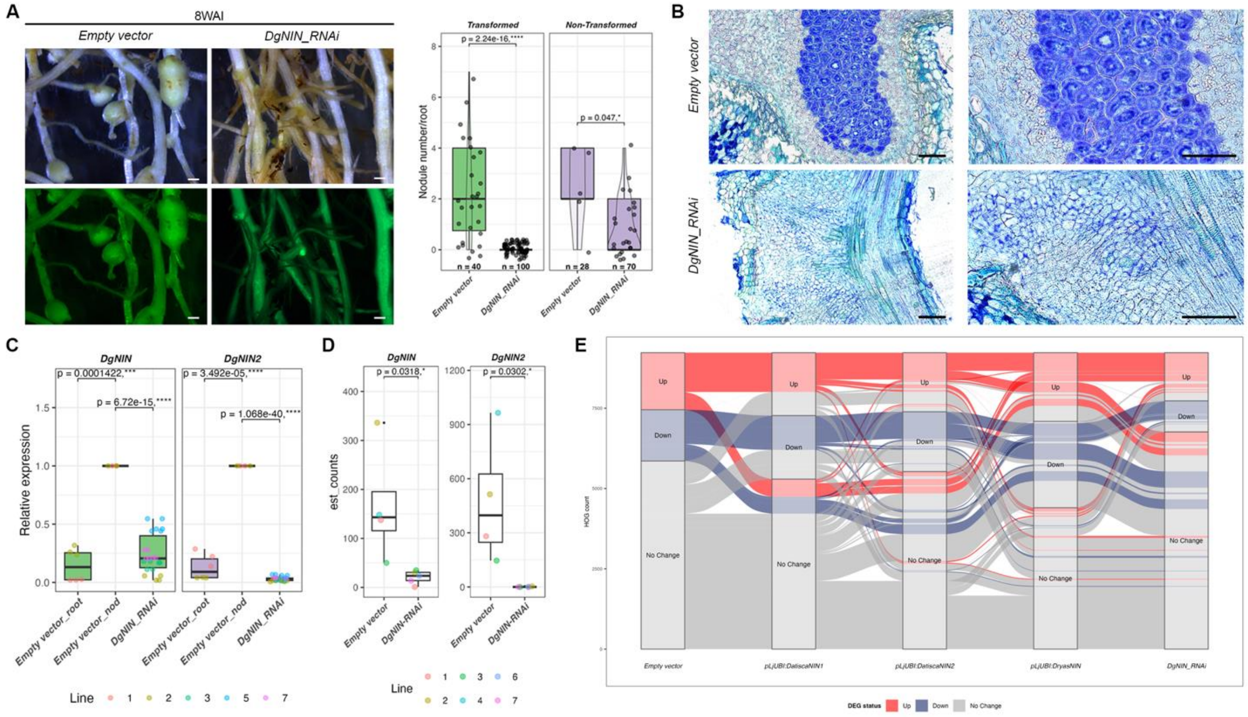


**Supplementary Fig. 3| Downregulation of *DgNIN* affected nodule development.** Hairy-root transformed *Datisca glomerata* plants transformed with Empty vector and *DgNIN_RNAi* harvested 8WAI with Dg1 clade 2 *Frankia* sp. showing (**A**) macroscopic view of the root. Scale bar, 500 µm. Boxplot represents the number of nodules per transformed root where median (thick line), second to third quartiles (box) and independent transformation experiment (coloured dots). Wilcoxon signed-rank test followed by Bonferroni correction represents statistical significance where the p value and the asterisk are represented over the brackets, n = number of transformed roots scored. (**B**) ultrastructure of root nodule (Empty vector) and root (*DgNIN_RNAi*) stained with toluidine blue. Scale bar, 100 µm. Boxplot represents the (**C**) Expression of *DgNIN* and *DgNIN2* determined by qRT-PCR relative to the endogenous control *DgActin* (**D**) Estimated counts of the individual transcript quantified from the RNAseq data relative to Empty vector transformed roots where median (thick line), second to third quartiles (box) and independent lines (coloured dots). (C and D) Student's t-test represents statistical significance where the p value and asterisk represented over the brackets (**E**) Alluvial-plot illustrating the relationship between *Empty vector*, *pLjUBI:DatiscaNIN*, *pLjUBI:DatiscaNIN2*, *pLjUBI:DryasNIN* and *DgNIN_RNAi* where the flow represents homologous gene expression pattern across three conditions, Up (> 0), Down (< 0) or No Change (0). The width of the flow indicates the number of Genes shared. See Supplementary Dataset 2.

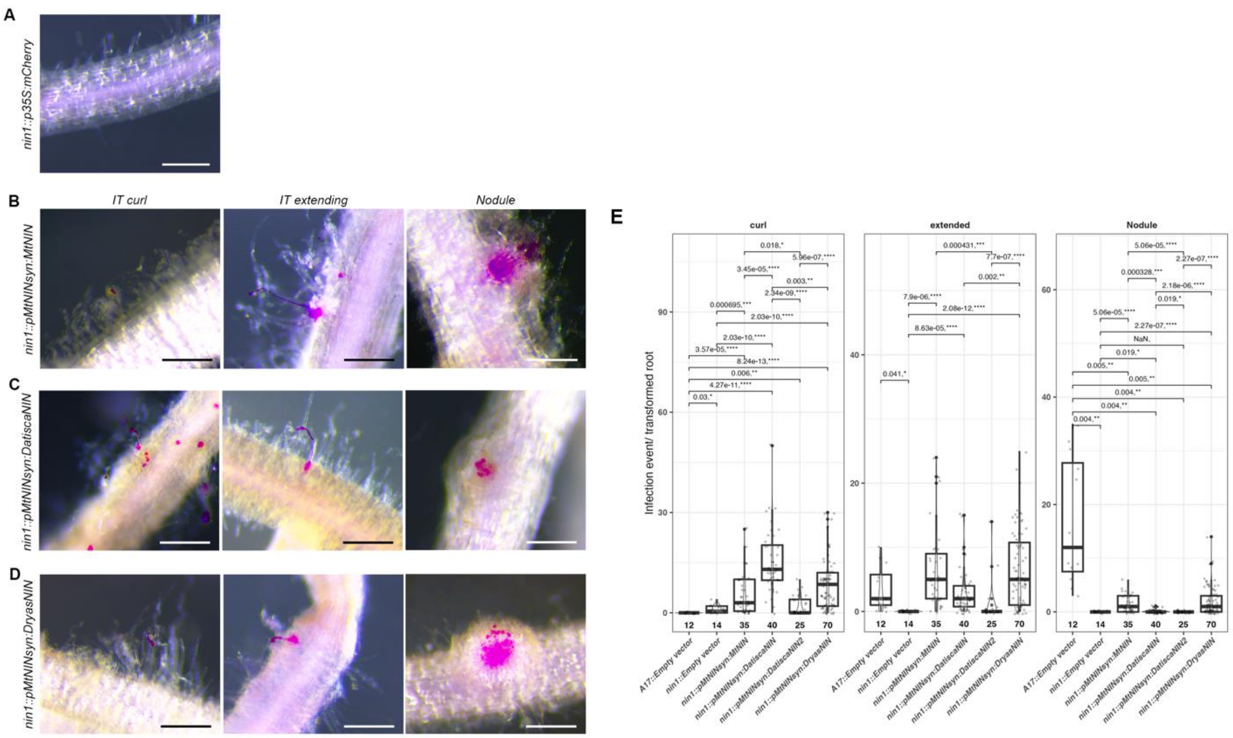


**Supplementary Fig. 4| Actinorhizal NIN can differentially rescue nodule development in *M. truncatula nin1* mutant.** Hairy-root transformed *M. truncatula nin1* with (**A**) empty vector (*p35S:mCherry*), (**B**) *pMtNINsyn:MtNIN*, (**C**) *pMtNINsyn:DatiscaNIN* and (**D**) *pMtNINsyn:DryasNIN* showing representative images for different infection phenotype namely IT (Infection Thread) curl, extending and nodule on transformed roots harvested 3 weeks after infection (WAI) with *S. meliloti* 2011-gfp (purple). Scale bar, 250 µm. (**E**) Boxplot represents the number of infection event per transformed root where median (thick line), second to third quartiles (box). Student's t-test represents statistical significance where the p value and the asterisk are represented over the brackets, n = number of transformed roots scored.

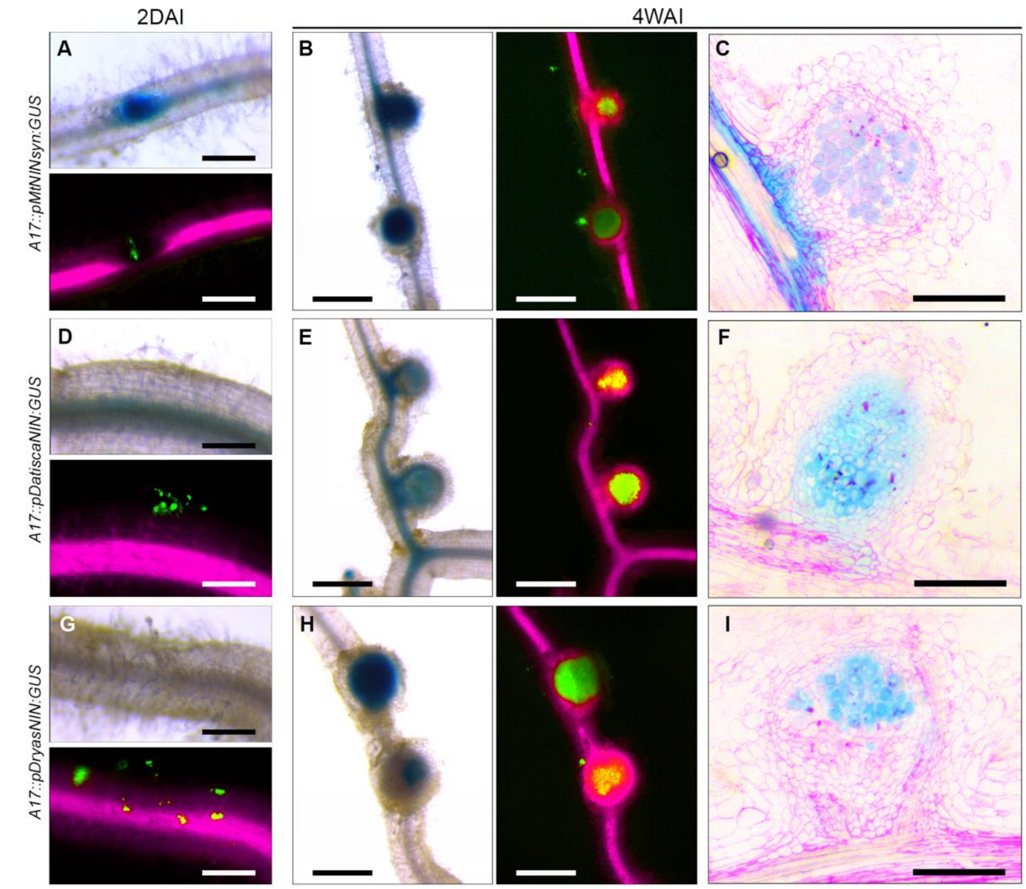


**Supplementary Fig. 5| Actinorhizal NIN promoter is activated during *M. truncatula* nodule development.** Hairy-root transformed *M. truncatula* A17 (wild-type) with (**A-C**) *pMtNINsyn:GUS*, (**D-F**) *pDatiscaNIN:GUS* and (**G-I**) *pDryasNIN:GUS* showing representative images for (**A**, **D** and **G**) early infection (2 day post infection, DPI) and (**B**, **E** and **H**) nodule (4WAI) where GUS stain (blue), transformation marker (mCherry, purple) and *S. meliloti* 2011-gfp (green). Scale bar, 250 μm (**A**, **D** and **G**); 500 μm (**B**, **E** and **H**). (**C**, **F** and **I**) Ultrathin section of nodule showing GUS stain (blue) and nodule ultrastructure (ruthenium red, purple). Scale bar, 250 μm.

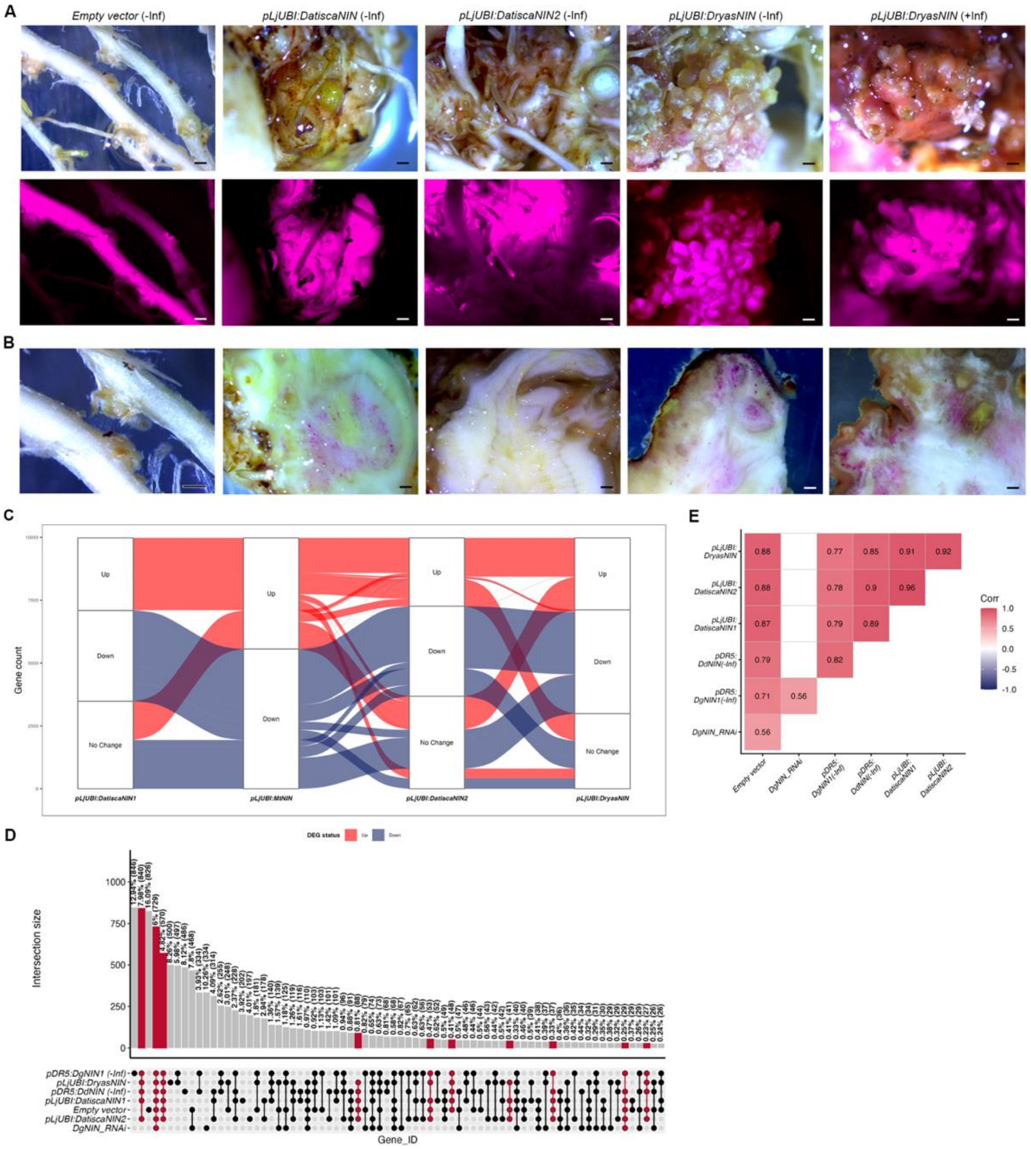


**Supplementary Fig. 6| Overexpression of *DryasNIN* in *D. glomerata* leads to formation of pseudonodules.** Macroscopic view of the (**A**) whole root showing spontaneous nodule like structure formation in *D. glomerata* hairy-root transformed with *Empty vector*, *pLjUBI:DatiscaNIN*, *pLjUBI:DatiscaNIN2* without infection (-Inf), *pLjUBI:DryasNIN* without (-Inf) and with infection (+Inf) where mCherry is transformation marker. Scale bar, 500 µm. (**B**) Half section of spontaneous nodule except for *Empty vector*. Scale bar, 500 µm. (**C**) Alluvial-plot illustrating the relationship between *pLjUBI:DatiscaNIN*, *MtNIN_OE* [1], *pLjUBI:DatiscaNIN2* and *pLjUBI:DryasNIN* where the flow represents homologous gene expression pattern across three conditions, Up (> 0), Down (< 0) or No Change (0). The width of the flow indicates the number of Genes shared and boxes shows representative gene name. m, meristem; vb, vascular bundle; iz, infection zone. (**D**) Upset plot showing all intersections for the Gene_IDs shared under different condition where the highlighted ones indicate unions shared between majority of the datasets. (**E**) Correlation plot is based on the gene expression from the highlighted unions where values represent Spearman's correlation coefficient (ρ), significance was tested using a two-sided t-test against ρ = 0 with significant values is based on FDR-adjusted p-value, with a threshold of $p < 0.0001$. Correlation that does not meet this significance threshold were left blank. See **Supplementary** Dataset 2.

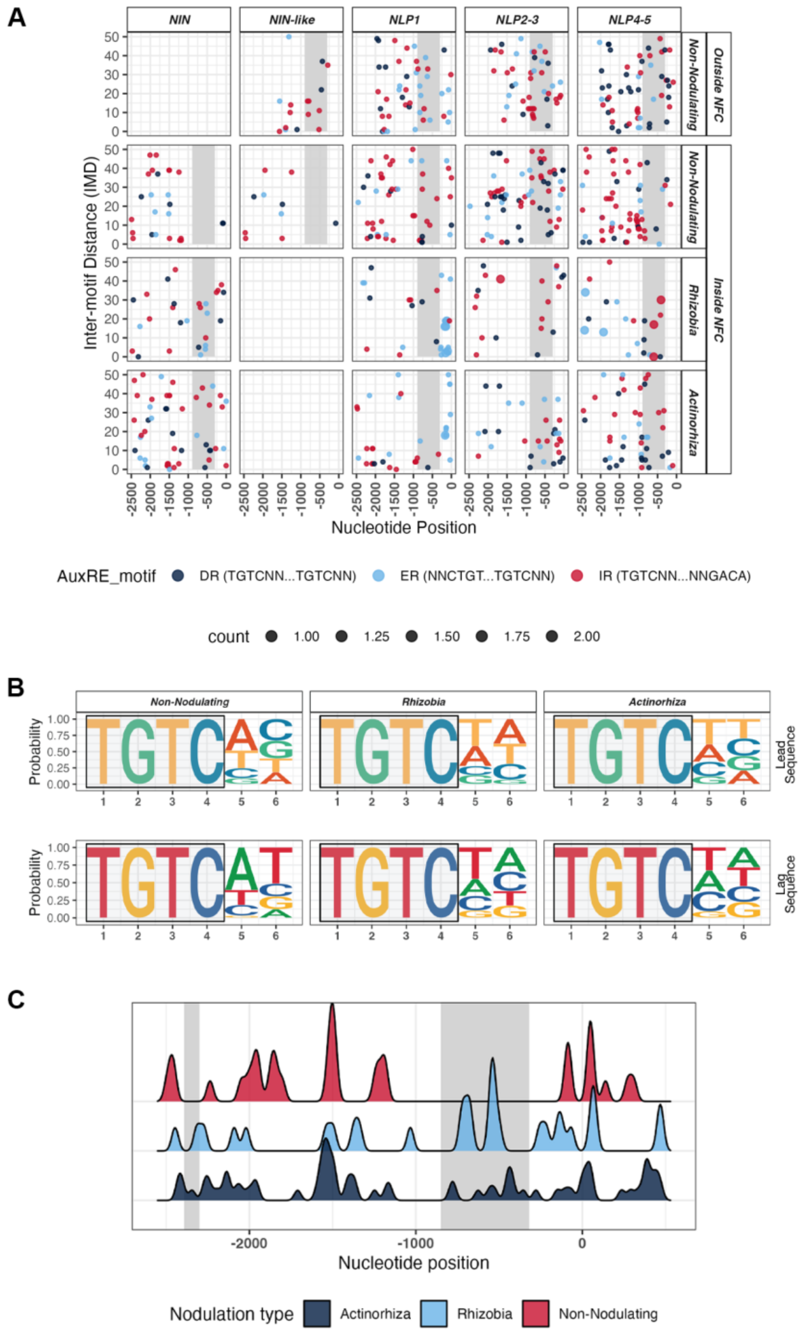


**Supplementary Fig. 7| Auxin responsive elements (*AuxRE*) in the NLP promoter.** (**A**) Dotplot showing Inter-motif distance (IMD) between the AuxRE pair present in the promoter of *NIN-like*, *NLP1*, *NLP2-3* and *NLP4-5* from within and outside of the NFC (Nitrogen fixing clade) mapped between -2.5kb and 500bp from the TSS (transcriptional start site, 0bp) where colour represents the relative cardinality of the *AuxRE* motifs with a p-value threshold <0.01. (**B**) Weblogo showing the nucleotide sequence of the leading and lagging *AuxRE* sequence. (**C**) Density plot of the NIN promoter region between non-nodulating, rhizobia and actinorhizal nodulators. (A and C) Gray highlighted region indicate the unique position of *AuxRE* in nodulating NIN. See Supplementary Table 2 and Supplementary Dataset 3.

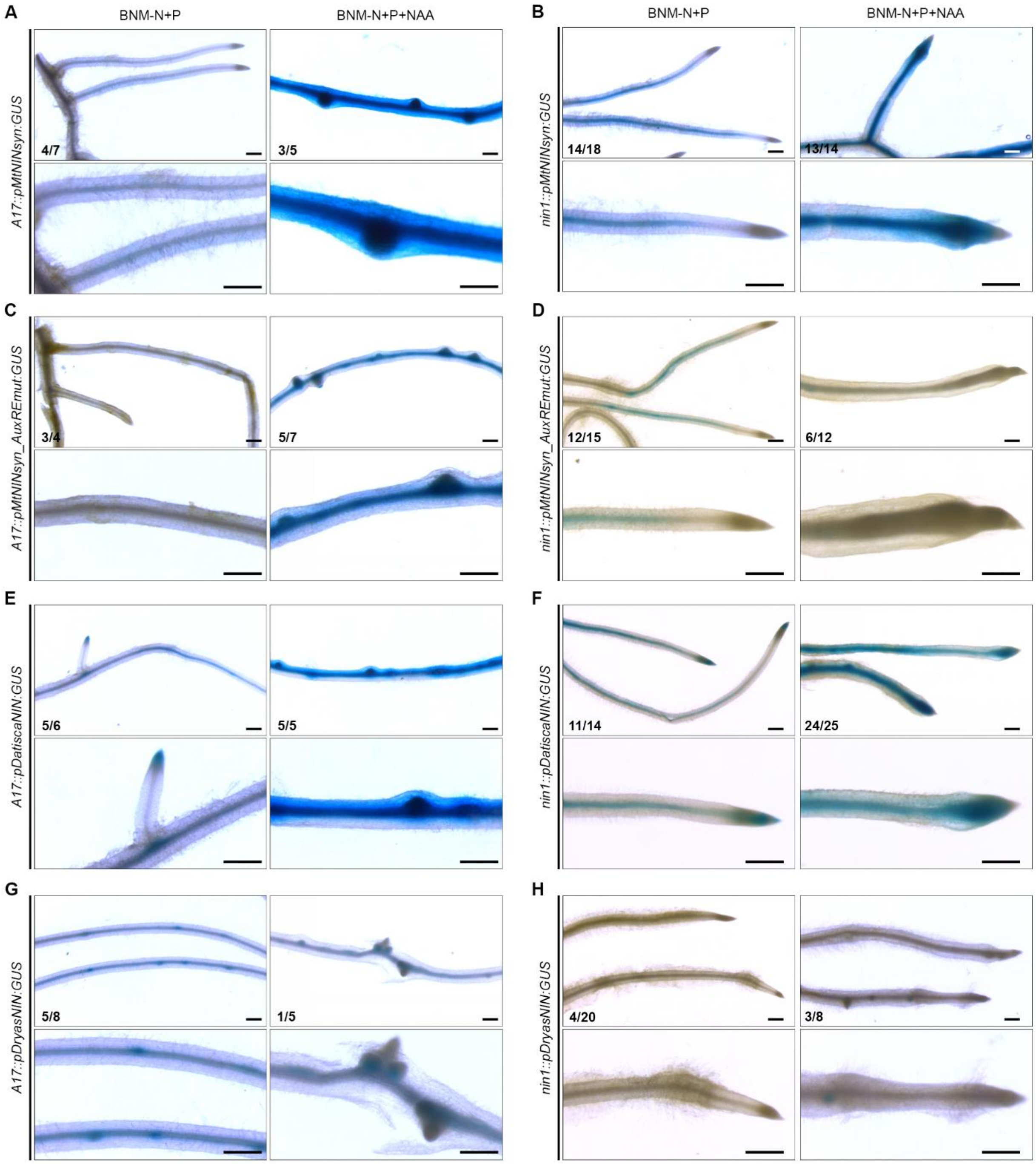


**Supplementary Fig. 8| Exogenous auxin mediated activation of *pNIN* in *Medicago truncatula*.** Hairy-root transformed *M. truncatula* (**A**, **C**, **E** and **G**) A17 and (**B**, **D**, **F** and **H**) *nin1* mutant transformed with *pMtNINsyn:GUS*, *pMtNINsyn_AuxREmut:MtNIN*, *pDatiscaNIN:GUS* and *pDryasNIN:GUS* treated under control (BNM-N+P) and 10 µM NAA (BNM-N+P+NAA) harvested 2 days post treatment (DPT) where GUS staining (blue) indicates promoter activation. The number at the bottom left indicates stained / transformed roots studied. Scale bar, 500 µm.

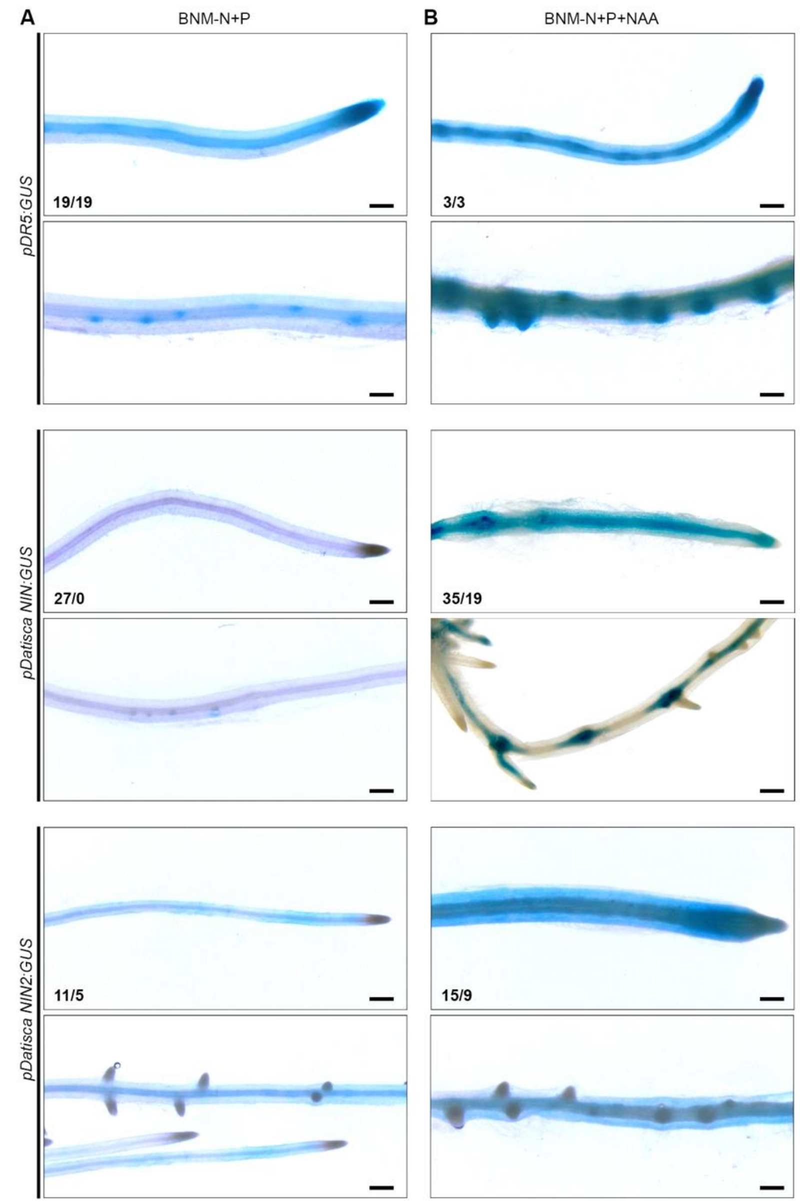


**Supplementary Fig. 9| Exogenous auxin mediated activation of *pNIN* in *Datisca glomerata*.** Hairy-root transformed *D. glomerata* transformed with *pDR5:GUS*, *pDatiscaNIN:GUS* and *pDatiscaNIN2:GUS* treated under (**A**) control (BNM-N+P) and (**B**) 10 µM NAA (BNM-N+P+NAA) harvested 3 days post treatment (DPT) where GUS staining (blue) indicates promoter activation. The number at the bottom left indicates stained / transformed roots studied. Scale bar, 500 µm.

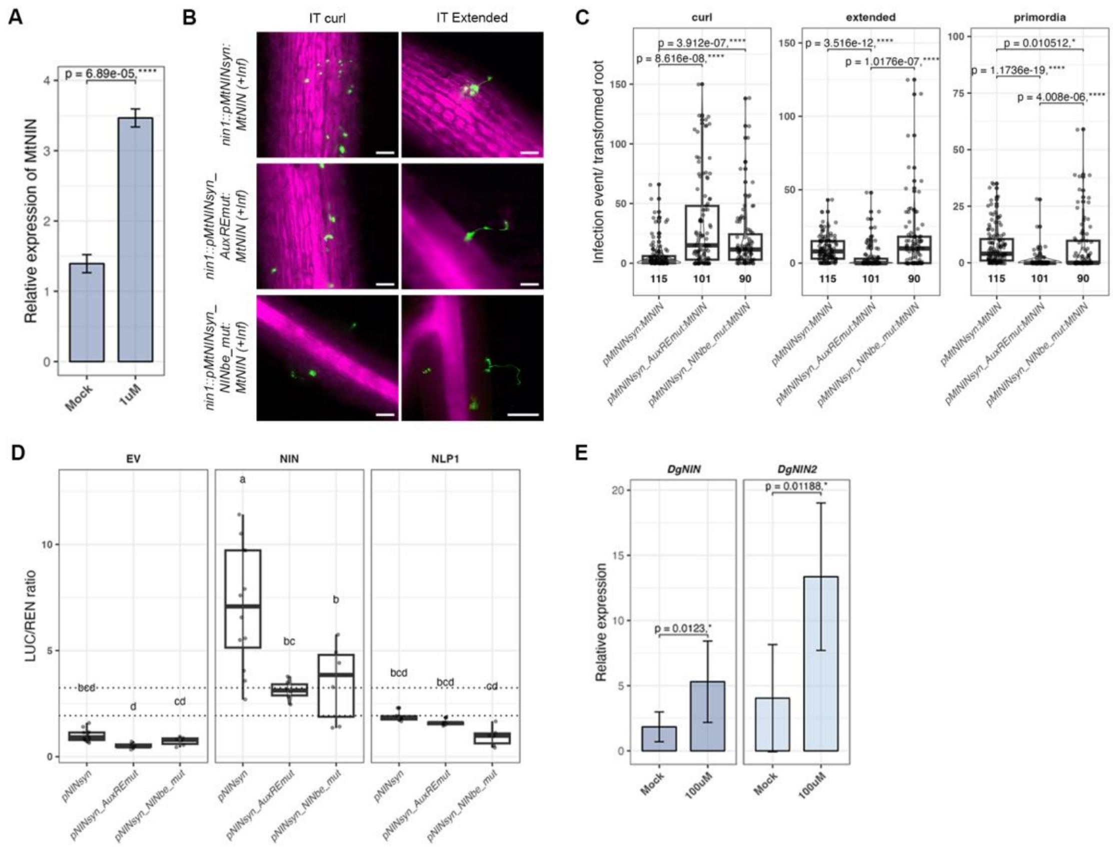


**Supplementary Fig. 10| Auxin responsive elements (*AuxRE*) in the NIN promoter are essential for nodule development.** The boxplot represents **(A)** expression as measured by qRT-PCR for *MtNIN* relative to reference gene *MtActin*, (B) Representative images for *M. truncatula nin1* transformed with *pMtNINsyn:MtNIN*, *pMtNINsyn_AuxREmut:MtNIN* and *pMtNINsyn_NINbe_mut:MtNIN* showing infection events 4WAI with *Sinorhizobium meliloti* 2011-GFP (green) where mCherry (magenta) is the transformation marker. Scale bar, 100 µm. (**C**) the number of infection events per transformed root where median (thick line), second to third quartiles (box) and independent transformation experiment (coloured dots). Wilcoxon signed-rank test represents statistical significance where the p value and the asterisk are represented over the brackets, n = number of transformed roots scored. Scale bar, 100 mm, (**D**) transactivation assay in *Nicotiana benthamiana*, *LUC (firefly luciferase)*; *REN (Renilla luciferase)*. For the ratio, median (thick line), second to third quartiles (box). Single letters represent statistically significant groupings analysed by two-way ANOVA followed by TukeyHSD, $p < 0.05$ and the pair of dotted lines represent the upper and lower limit of the confidence interval, (**E**) expression as measured by qRT-PCR for *DgNIN* and *DgNIN2* relative to reference gene *DgActin* where Error bars represent SD and means were compared using Student's t-test where the p value and the asterisk represent over the brackets represent statistical significance. (A and D) NAA (naphthalene acetic acid) concentration 1µM and 100 µM.

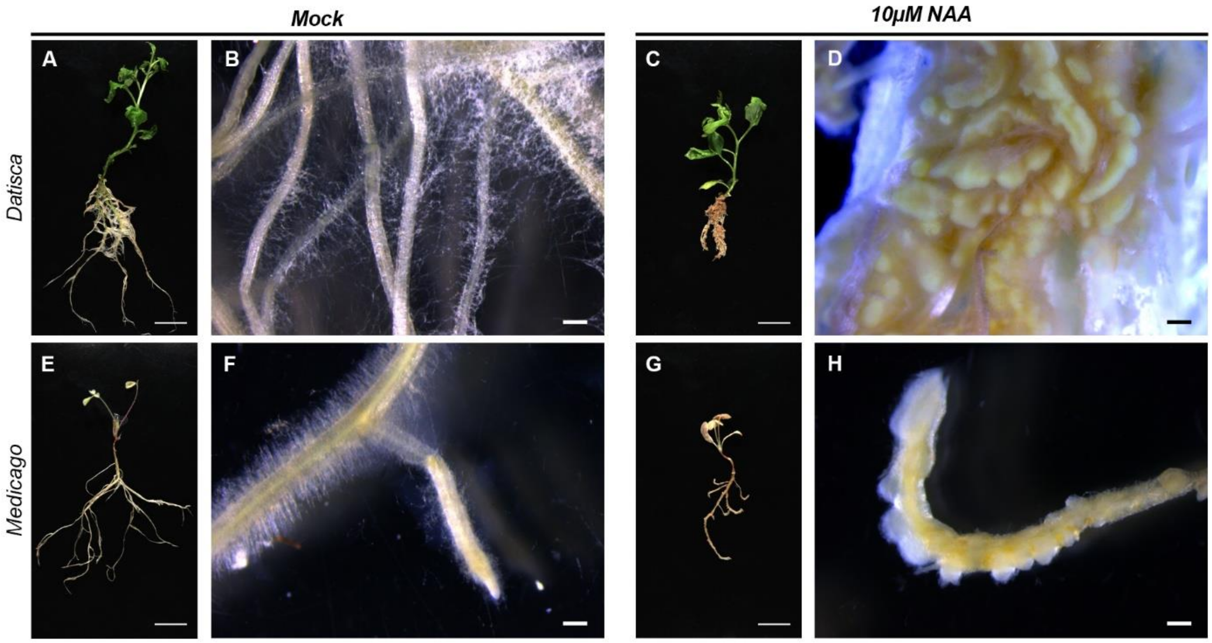


**Supplementary Fig. 11| Exogenous auxin treatment leads to spontaneous nodule-like structures.** Macroscopic images of (**A-D**) *Datisca glomerata* and (**E-H**) *Medicago truncatula* A17 seedling treated by (**A-B** and **E-F**) Mock and (**C-D** and **G-H**) 10 μM NAA (Naphthaleneacetic acid) harvested 2 weeks post treatment. Scale bar, 2.4cm (A, C, E and G); 500 μm (B, D, F and H).

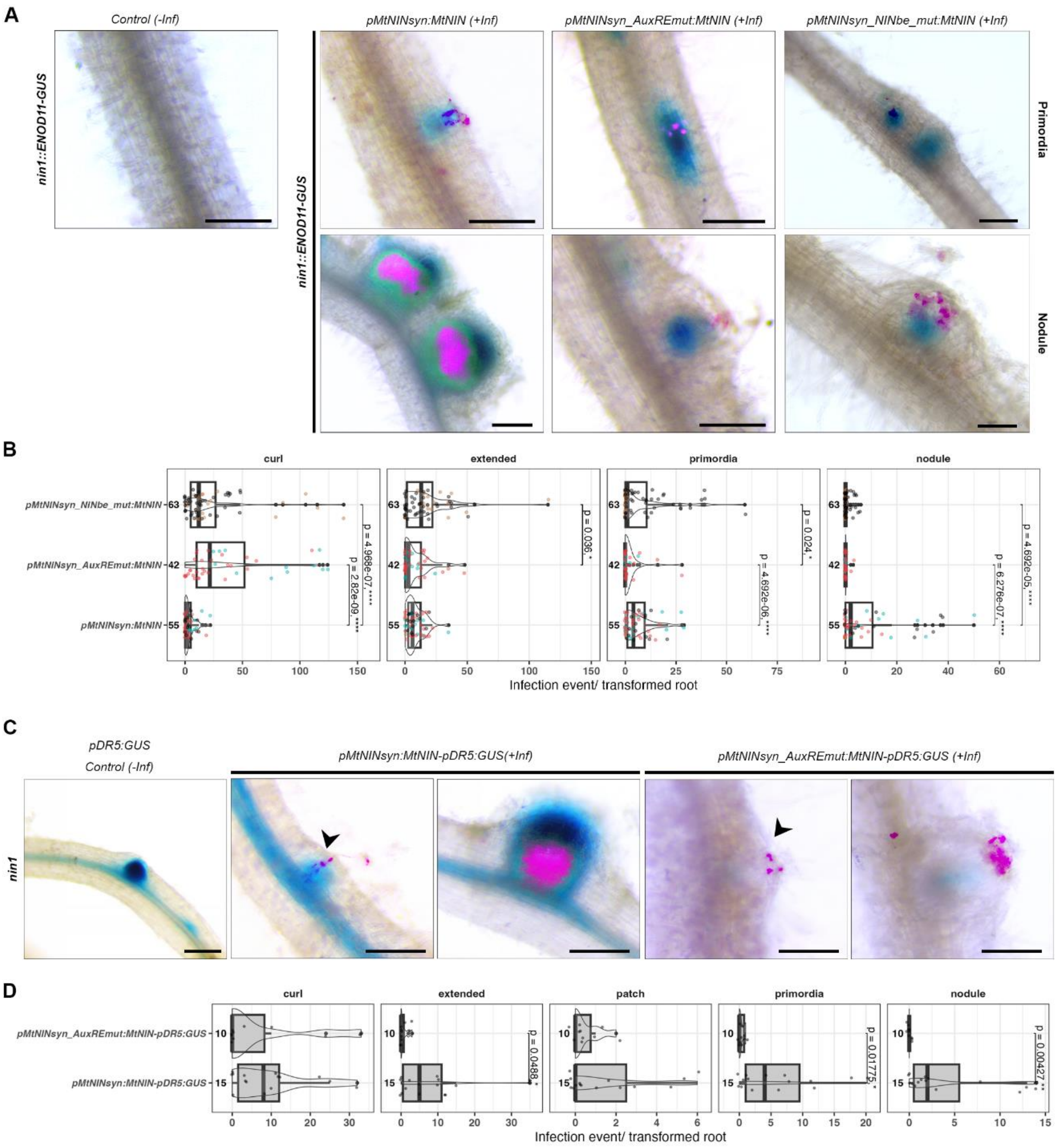


**Supplementary Fig. 12| *AuxRE* in *pMtNINsyn* is responsible for NIN activation.** Representative images for *Medicago truncatula* (**A**) *nin1::ENOD11:GUS* mutant hairy-root transformed with *pMtNINsyn:MtNIN*, *pMtNINsyn_AuxREmut:MtNIN* and *pMtNINsyn_NINbe_mut:MtNIN* showing primordia and nodule morphology where GUS staining (blue) shows *pENOD11* activation, (**C**) *nin1* mutant hairy-root transformed with *pMtNINsyn:MtNIN-pDR5:GUS* and *pMtNINsyn_AuxREmut:MtNIN-pDR5:GUS* where GUS staining (blue) shows *pDR5* activation. Transformed roots are harvested 4WAI with *Sinorhizobium meliloti* 2011-gfp (purple) Scale bar, 250 μm. (**B** and **D**) The boxplot represents the number of infection events per transformed root where median (thick line), second to third quartiles (box) and independent transformation experiment (coloured dots). Wilcoxon signed-rank test followed by Bonferroni correction represents statistical significance where the p value and the asterisk are represented over the brackets, n = number of transformed roots scored.

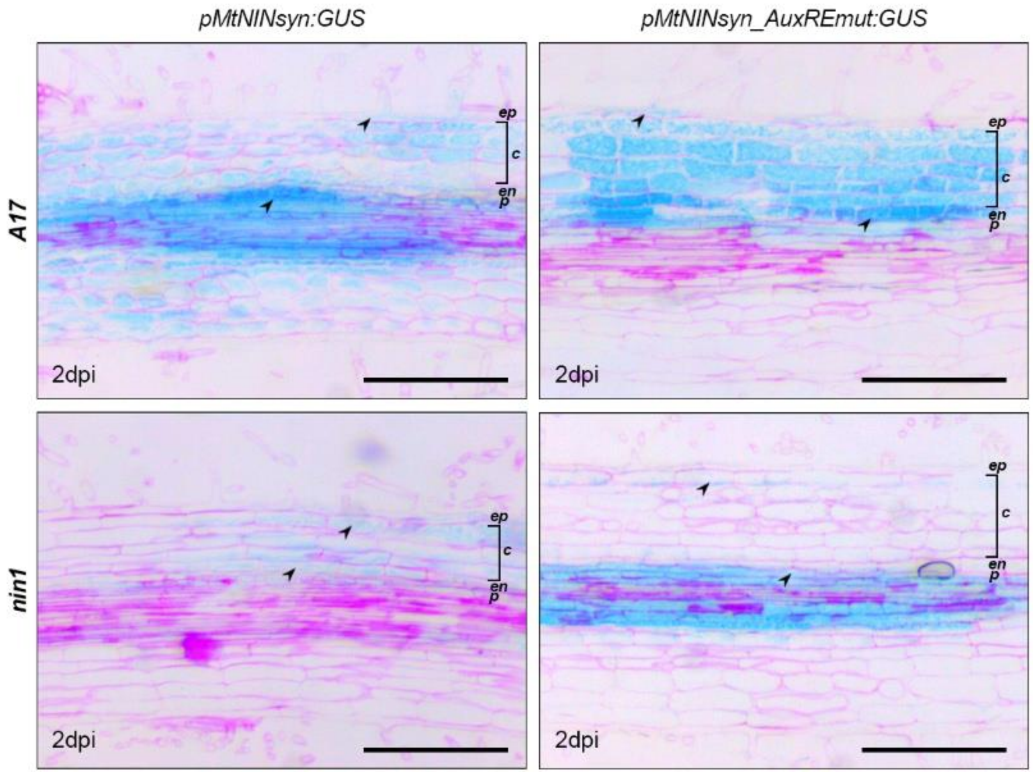


**Supplementary Fig. 13| *AuxRE* in *pMtNINsyn* is responsible for early activation during infection.** Representative images of ultrathin section of *Medicago truncatula* A17 and *nin1* mutant hairy-root transformed with *pMtNINsyn:GUS* and *pMtNINsyn_AuxREmut:GUS* spot inoculated with *Sinorhizobium meliloti* 2011-gfp harvested 2 days post inoculation (2dpi). The sections show promoter activity (arrowhead) counter stained with ruthenium red where ep (epidermis), c (cortex), en (endodermis) and p (pericycle). Scale bar, 250 μm.

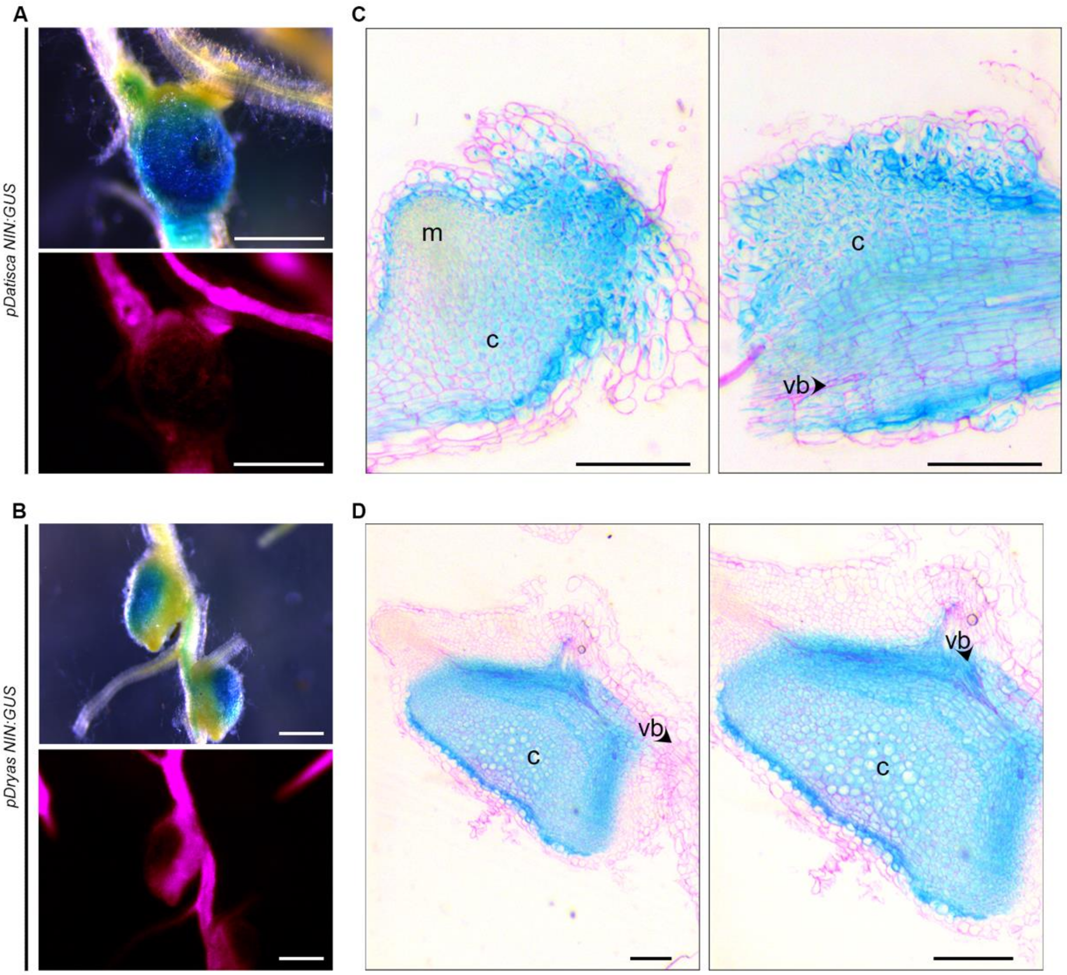


**Supplementary Fig. 14| Actinorhizal *NIN* express early during nodule development.** Representative images for hairy-root transformed *D. glomerata* transformed with *pDatiscaNIN:GUS* and *pDryasNIN:GUS* harvested 8 weeks after infection (8WAI) with clade 2 *Frankia* Dg1 showing (**A-B**) macroscopic view of early primordia and (**C-D**) ultrathin section. GUS stain (blue) represents promoter activity, mCherry transformation marker (purple) and ruthenium red (C-D). m (meristem), c (cortex) and vb (vascular bundle). Scale bar, 500 µm (A-B); 250 µm (C-D).

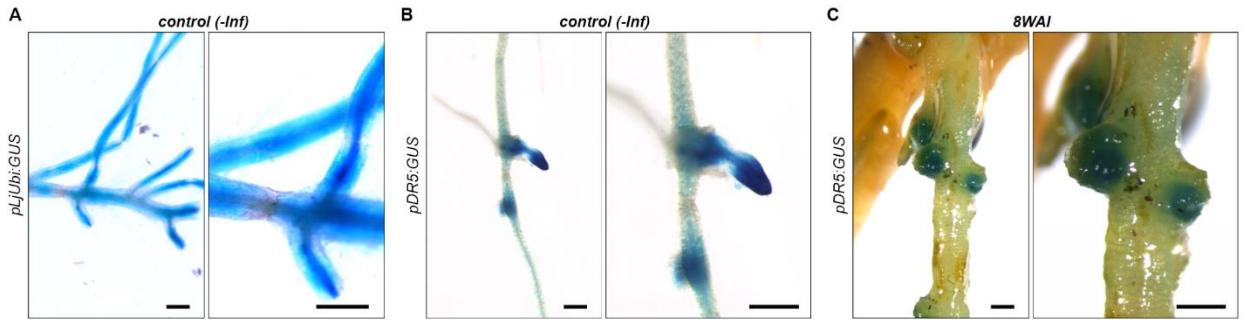


**Supplementary Fig. 15| Spatial difference in expression level for *pDR5* in *Datisca glomerata*.** Representative images for hairy-root transformed *D. glomerata* transformed with (**A**) *pLjUbi:GUS* and (**B-C**) *pDR5:GUS* under control (-Inf) and 8 weeks after infection (8WAI) with clade 2 *Frankia* Dg1. GUS stain (blue) represents promoter activity. Scale bar, 500 μm.

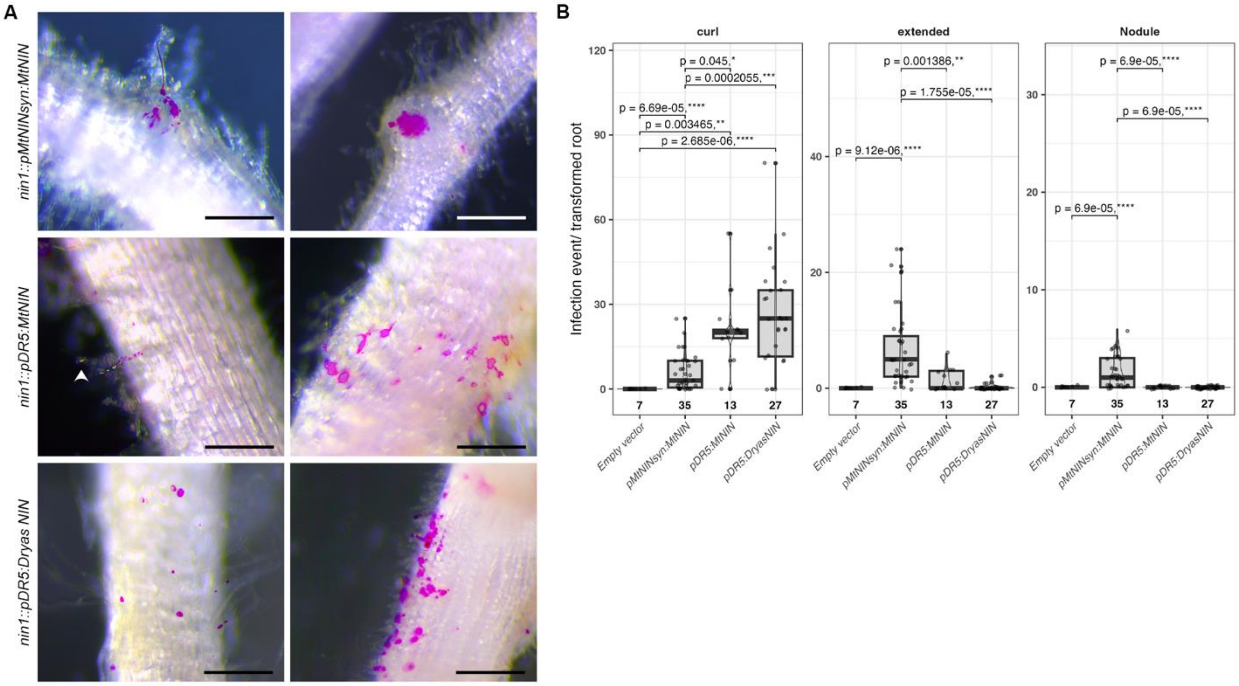


**Supplementary Fig. 16| *pDR5* is not sufficient to rescue nodulation in *Medicago truncatula*.** Representative images for *Medicago truncatula nin1* mutant transformed with (**A**) *pMtNINsyn:MtNIN*, (**B**) *pDryasNIN:DryasNIN*, (**C**) *pDR5:MtNIN* and (**D**) *pDR5:DryasNIN* harvested 4 weeks after infection (WAI) with *Sinorhizobium meliloti* 2011-gfp (purple). Arrowhead: Infection curl; Scale bar, 250 μm. The boxplot represents the number of (**E**) infection events and (**F**) nodules per transformed root where median (thick line), second to third quartiles (box) and independent transformation experiment (coloured dots). Student's t-test represents statistical significance where the p-value and the asterisk are represented over the brackets, n = number of transformed roots scored.

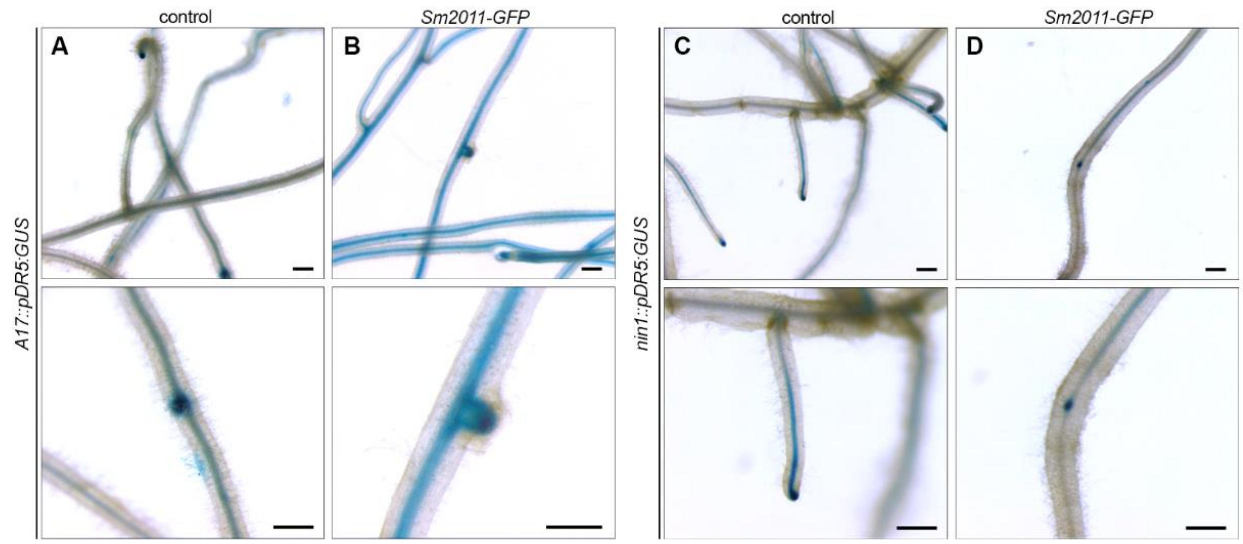


**Supplementary Fig. 17| *pDR5* expression in *Medicago truncatula*.** Representative images of hairy-root transformed *M. truncatula* roots for (**A-B**) A17 and (**C-D**) *nin1* transformed with *pDR5:GUS* harvested under control (**A, C**) and 4 weeks after infection, WAI (**B, D**) with *Sinorhizobium meliloti* 2011-gfp. Scale bar, 500 μm.

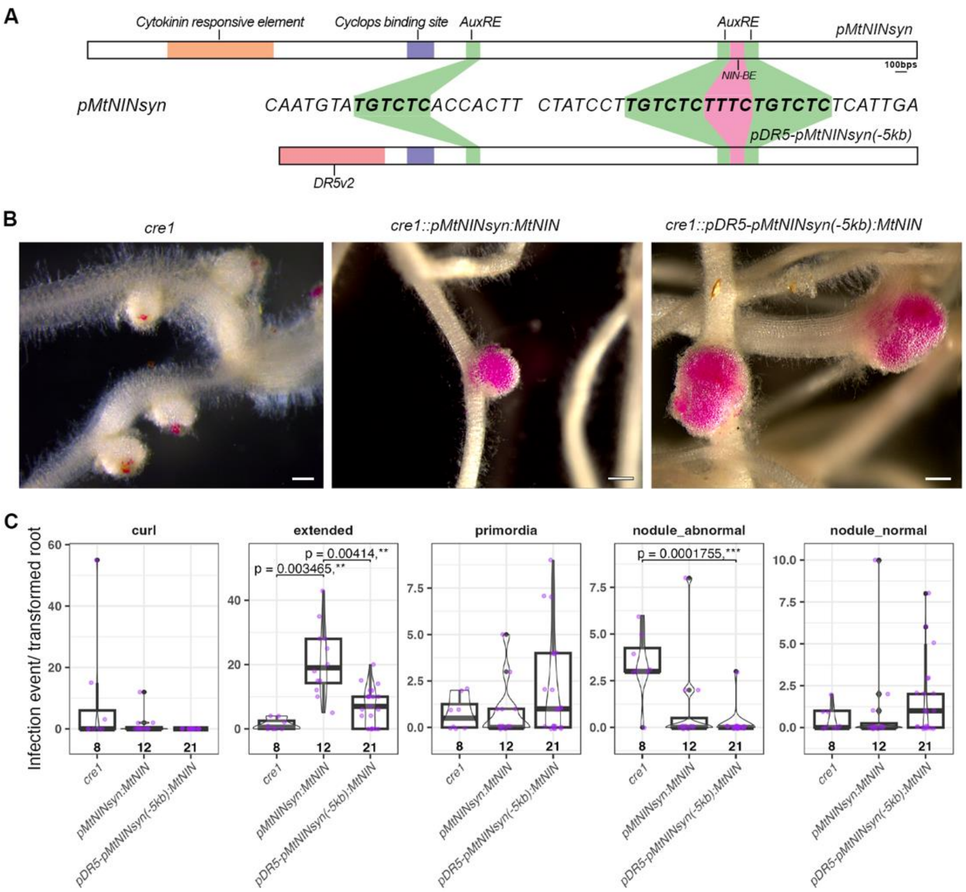


**Supplementary Fig. 18| *pDR5-pMtNINsyn(-5kb)* minimal promoter can induce cortical cell division in *Medicago truncatula cre1-1* mutant. (A)** Schematic representation of the synthetic promoters. Representative images of *M. truncatula cre1* mutant hairy-root transformed with (**B**) *pMtNINsyn:MtNIN* and *pDR5-pMtNINsyn(-5kb):MtNIN* harvested 4 weeks after infection (WAI) with *Sinorhizobium meliloti* 2011-gfp (purple) where *cre1* represent nodules formed in the untransformed roots. Scale bar, 100 µm. (**C**) The boxplot represents the number of infection events per transformed root where median (thick line), second to third quartiles (box) and independent transformation experiment (coloured dots). Wilcoxon test followed by Bonferroni correction represents statistical significance where the p value and the asterisk are represented over the brackets, n = number of transformed roots scored. nodule_abnormal, nodule with aberrant infection at the apex and nodule_normal, proper infection.

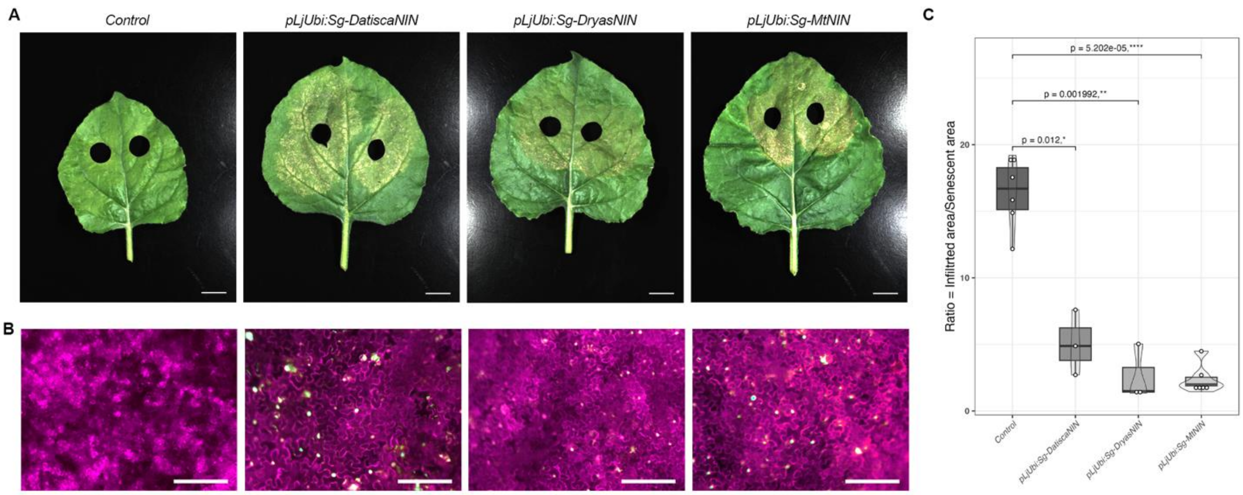


**Supplementary Fig. 19| Transient expression of NIN in *Nicotiana benthamiana* leads to early senescence.** *N. benthamiana* leaves infiltrated with Control (*pMtNINsyn:GUS*), *pLjUbi:Sg-MtNIN*, *pLjUbi:Sg-DatiscaNIN* and *pLjUbi:Sg-DryasNIN* under (**A**) bright-field and (**B**) fluorescence images showing mCherry (transformation marker, purple) and Sg (Staygold) fused NIN protein (green). Scale bar, 2.4 cm (A) and 250 μm (B). (**C**) Boxplot showing the Ratio = Infiltrated area/Senescent area where median (thick line), second to third quartiles (box). Student's t-test represents statistical significance where the p value and the asterisk are represented over the brackets.

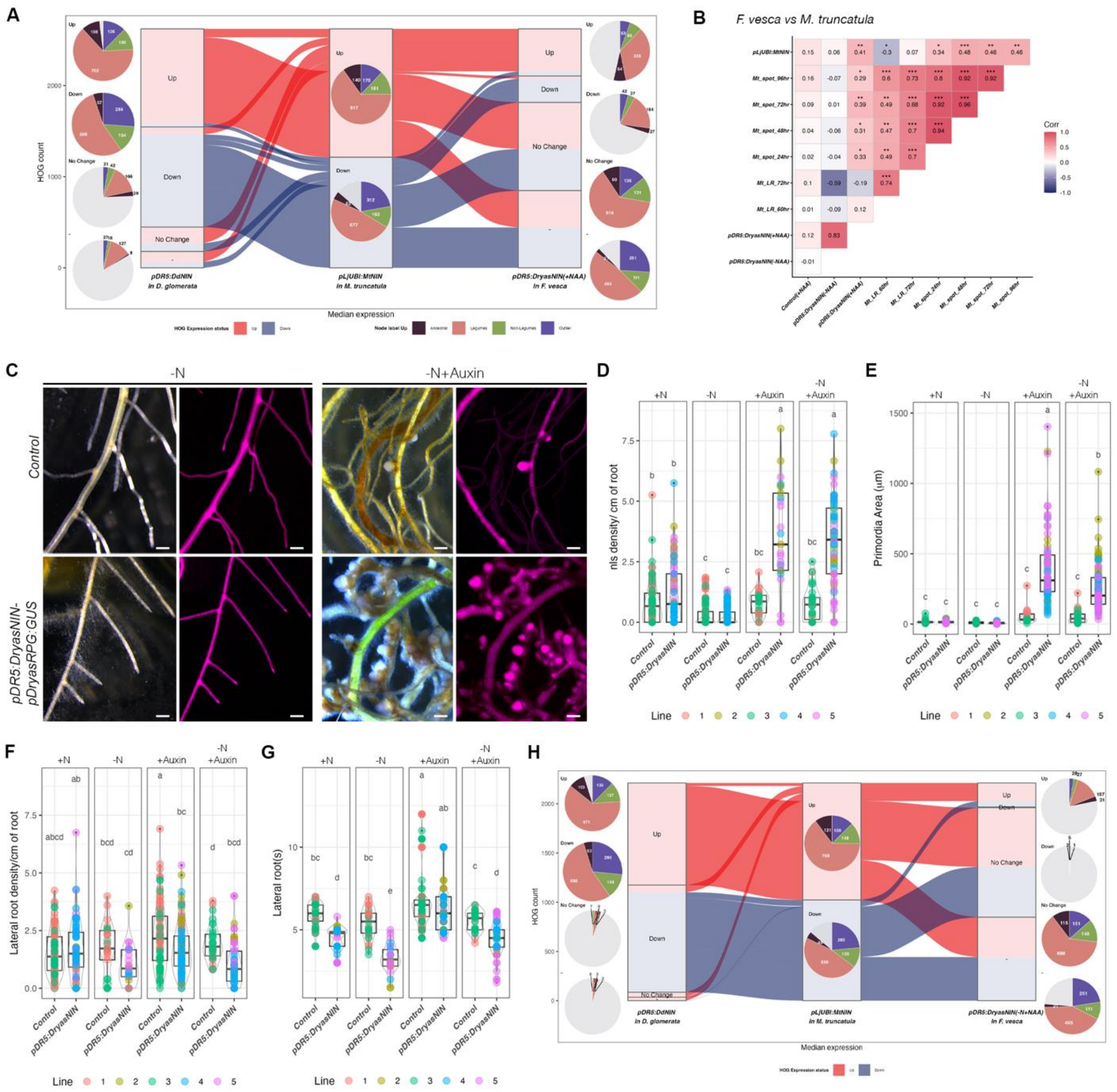


**Supplementary Fig. 20| Synthetic auxin responsive element driving NIN can induce spontaneous nodule-like structures (nls) in *Fragaria vesca* (*Fv*, strawberry) independent of Nitrogen (N) status. (A)** Alluvial-plot illustrating the relationship between median gene expression in each head orthogroup (HOG) for *pLjUBI:MtNIN* in *M. truncatula*, *pDR5:DryasNIN* in *D. glomerata* and *pDR5:DryasNIN (+NAA)* in *F. vesca* where the Up (>0) and Down (<0) HOGs in *M. truncatula* are compared against Up (>0) and Down (<0) HOGs in *D. glomerata* and *F. vesca pDR5:DryasNIN* datasets. (B) Polychoric correlation analysis for annotated Up (>0) and Down (<0) genes for *pDR5:DryasNIN (+NAA)* in *F. vesca* compared against *M. truncatula pLjUBI:MtNIN (4)*, spot and lateral root timepoint datasets *(3)* where statistical significance for each pairwise correlation was assessed using the asymptotic p-value with ***, p<0.0001. (**C**) Stereofluorescence images of *F. vesca* stable lines transformed with Control (*pDryasRPG:GUS*) and *pDR5:DryasNIN-pDryasRPG:GUS* treated under -N+P and -N+P+NAA (100 µM) harvested 4 weeks post treatment (4 WPT) where mCherry (transformation marker, purple). Scale bar, 500 µm. Boxplot represents the (**D**) nls density, (**E**) Primordia area per transformed root, (**F**) lateral root density and (**G**) lateral root number where median (thick line), second to third quartiles (box) and independent lines (coloured dots). Single Letters represent statistically significant groupings analyzed by two-way ANOVA followed by TukeyHSD, $p < 0.05$. (**H**) Alluvial-plot illustrating the relationship between median gene expression in each head orthogroup (HOG) for *pLjUBI:MtNIN* in *M. truncatula*, *pDR5:DryasNIN* in *D. glomerata* and *pDR5:DryasNIN (-N+NAA)* in *F. vesca* where the Up (>0) and Down (<0) HOGs in *M. truncatula* are compared against Up (>0) and Down (<0) HOGs in *D. glomerata* and *F. vesca pDR5:DryasNIN* datasets. (**A** and **H**) No Change (= 0) and – (Orthogroup not assigned). The pie charts represent the "Node label Up" from ancestral HOG identity based on Libourel et. al., 2023 where number represent the number of HOGs classified as Ancestral, Nodule, Non-nodule and Outlier. See Supplementary Dataset 5.

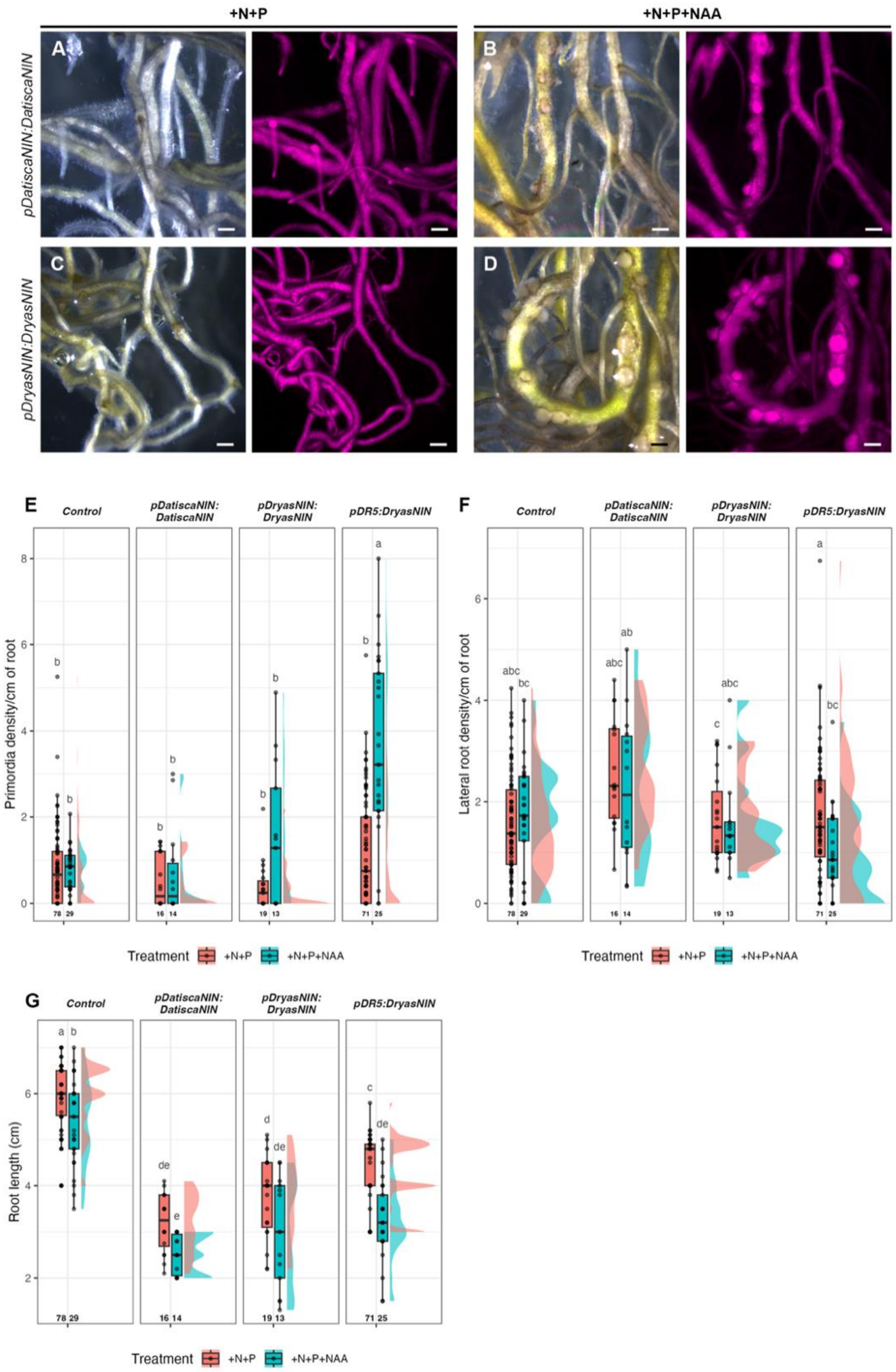


**Supplementary Fig. 21| Endogenous promoter driving NIN in *Fragaria vesca* (Fv, strawberry) is not efficient in inducing nodule like structures (nls).** Stereofluorescence images of *F. vesca* stable lines transformed with (**A-B**) *pDatiscaNIN:DatiscaNIN* and (**C-D**) *pDryasNIN:DryasNIN* treated under (**A, C**) +N+P and (**B, D**) +N+P+NAA (100 µM) harvested 4 weeks post treatment (4 WPT) where mCherry (transformation marker, purple). Scale bar, 500 µm. Boxplot represents the (**E**) primordia density, (**F**) lateral root density and (**G**) root length where median (thick line), second to third quartiles (box). Single

Letters represent statistically significant groupings analyzed by two-way ANOVA followed by TukeyHSD, $p < 0.05$.

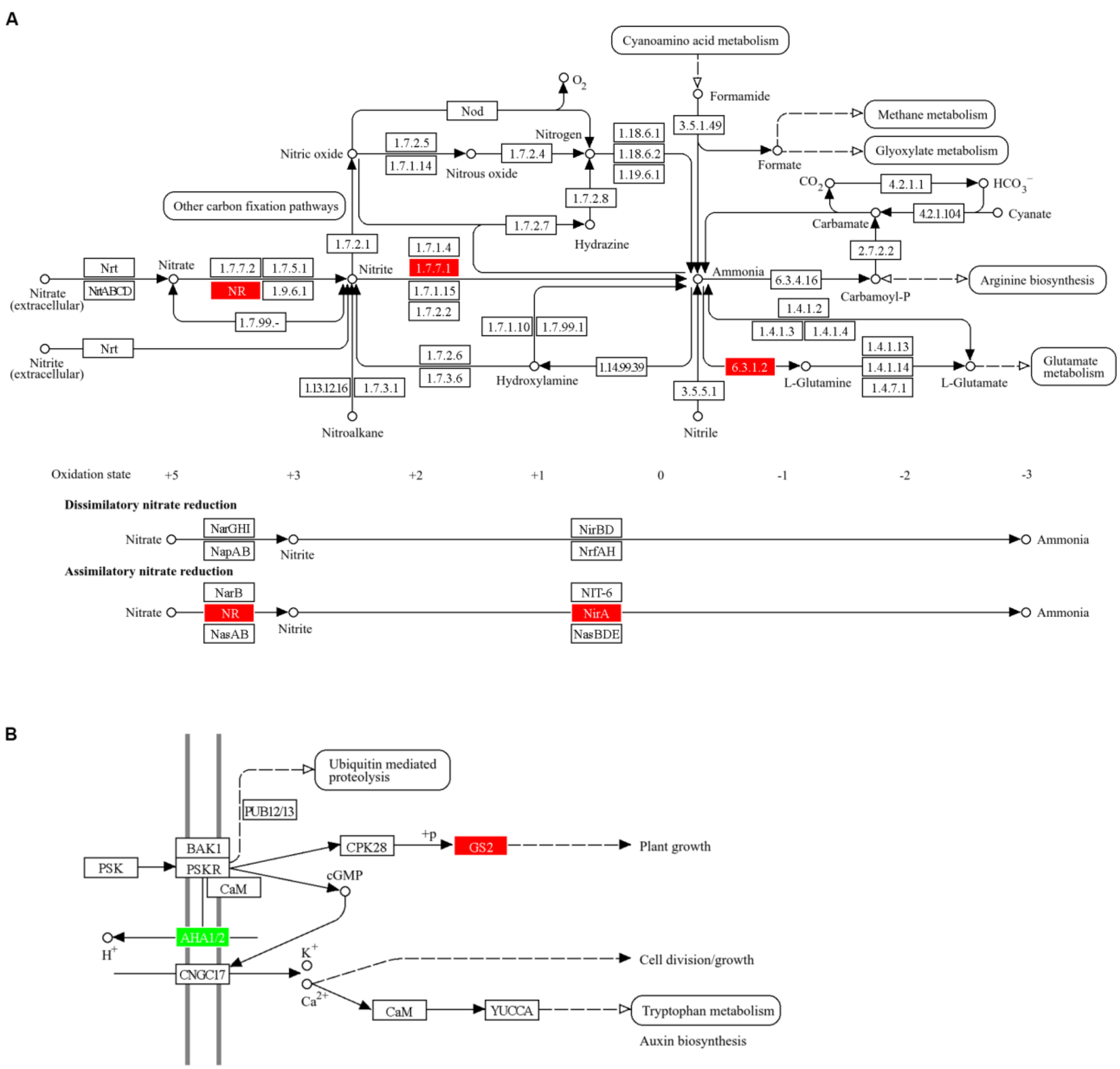


**Supplementary Fig. 22| KEGG pathway analysis.** Differentially expressed genes for *pDR5:DryasNIN (+NAA) in Fragaria vesca* where upregulated enzymes were highlighted in red and downregulated enzymes in green. The figure illustrates (**A**) Nitrogen metabolism and (**B**) Plant growth pathways affected due to differential expression of genes.

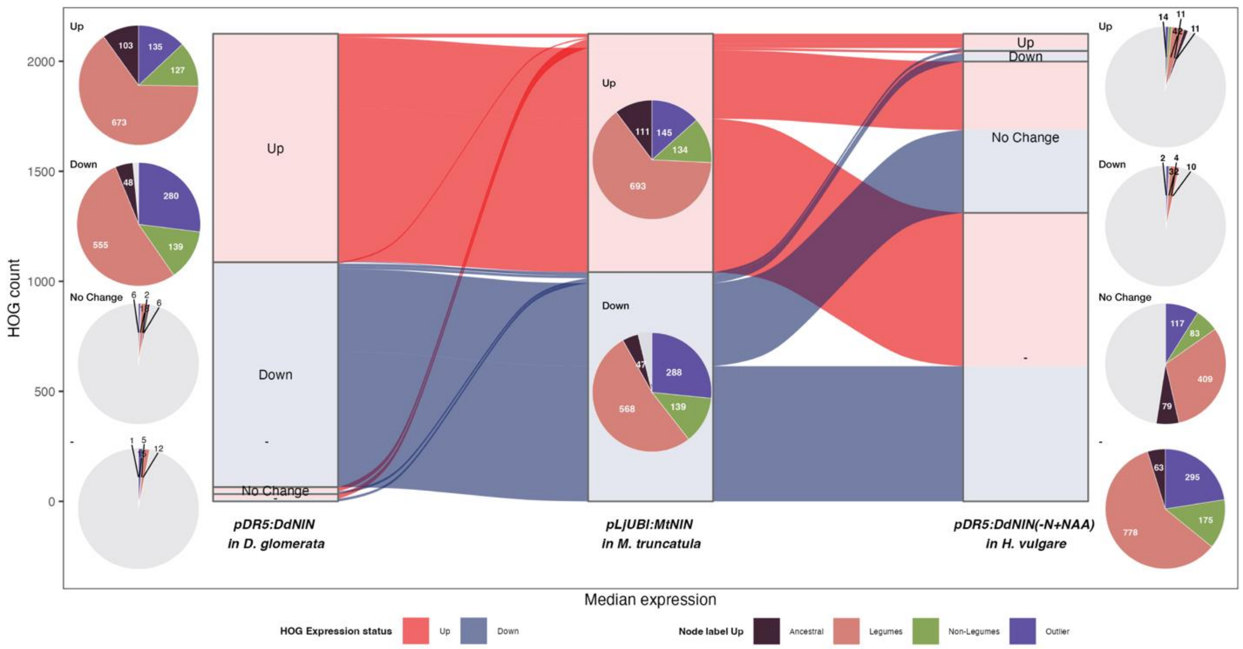


**Supplementary Fig. 23| Alluvial-plot illustrating the relationship between median gene expression in each head orthogroup (HOG)** for *pLjUBI:MtNIN* in *M. truncatula*, *pDR5:DryasNIN* in *D. glomerata* and *pDR5:DryasNIN* (-N+NAA) in *H. vulgare* where the Up (>0) and Down (<0) HOGs in *M. truncatula* are compared against Up (>0) and Down (<0) HOGs in *D. glomerata* and *H. vulgare pDR5:DryasNIN* datasets. No Change (= 0) and Orthologue absent (-). The pie charts represent the “Node label Up” from ancestral HOG identity based on Libourel et. al., 2023 where number represent the number of HOGs classified as Ancestral, Nodule, Non-nodule and Outlier. See Supplementary Dataset 6.

**Supplementary Table 1: Metadata of the Maximum-likelihood phylogegetic tree of NIN/NLP1 clade**

| **Node ID** | **Scientific name** | **Protein** | **Nodul ator** | **Family** | **Order** |
|---|---|---|---|---|---|
| *Bdis_Bradi1g76340.3.p* | *Brachypodium distachyon* | NLP1 | 0 | Poaceae | Poales |
| *Osat_LOC_Os03g03900.1* | *Oryza sativa* | NLP1 | 0 | Poaceae | Poales |
| *Bdis_Bradi4g37147.2.p* | *Brachypodium distachyon* | NLP1 | 0 | Poaceae | Poales |
| *Osat_LOC_Os09g37710.2* | *Oryza sativa* | NLP1 | 0 | Poaceae | Poales |
| *Mgut_Migut.B00011.1.p_NIN* | *Mimulus guttatus* | NIN | 0 | Phrymacea e | Lamiales |
| *Slyc_Solyc01g112190.2.1_NIN* | *Solanum lycopersicum* | NIN | 0 | Solanaceae | Solanales |
| *Ptrc_Potri.004G205500.1_NIN* | *Populus trichocarpa* | NIN | 0 | Salicaceae | Malpighiales |
| *Ptrc_Potri.009G166900.1_NIN* | *Populus trichocarpa* | NIN | 0 | Salicaceae | Malpighiales |
| *Mesc_Manes.11G110900.1.p_NIN* | *Manihot esculenta* | NIN | 0 | Euphorbiac eae | Malpighiales |
| *Rcom_29623.m000322_NIN* | *Ricinus communis* | NIN | 0 | Euphorbiac eae | Malpighiales |
| *Dglo_Datgl1955S23745_NIN* | *Datisca glomerata* | NIN | 1 | Datiscacea e | Cucurbitales |
| *Datcan_jg3638.t1_NIN* | *Datisca cannabina* | NIN | 1 | Datiscacea e | Cucurbitales |
| *Pper_ppa018195m_NIN* | *Prunus persica* | NIN | 0 | Rosaceae | Rosales |
| *Ddru_Drydr380S11191_NIN* | *Dryas drummmondii* | NIN | 1 | Rosaceae | Rosales |
| *Cercocarpus_NIN* | *Cercocarpus betuloides* | NIN | 1 | Rosaceae | Rosales |
| *Zjuj_XP_015893088.1_NIN* | *Ziziphus jujuba* | NIN | 0 | Rhamnacea e | Rosales |
| *Zjuj_XP_015893083.1_NIN* | *Ziziphus jujuba* | NIN | 0 | Rhamnacea e | Rosales |
| *Dtri_Distr1414S00745_NIN* | *Discaria trinervis* | NIN | 1 | Rhamnacea e | Rosales |
| *Cgla_Casgl993S23747_NIN* | *Casuarina glauca* | NIN | 1 | Casuarinac eae | Fagales |
| *Aglu_Alngl424567S27630_NIN* | *Alnus glutinosa* | NIN | 1 | Betulaceae | Fagales |
| *Jreg_WALNUT_00004000-RA_NIN* | *Juglans regia* | NIN | 0 | Juglandace ae | Fagales |
| *Vvin_GSVIVT01023886001_NIN* | *Vitis vinifera* | NIN | 0 | Vitaceae | Vitales |
| *Dglo_Datgl_scaffold169_1468791..1472708_NIN* | *Datisca glomerata* | NIN | 1 | Datiscacea e | Cucurbitales |
| *Datcan_jg4080.t1_NIN* | *Datisca cannabina* | NIN | 1 | Datiscacea e | Cucurbitales |

| | | | | | |
|---|---|---|---|---|---|
| *Hlup_HL.SW.v1.0.G010120.1_NIN* | *Humulus lupulus* | NIN | 0 | Cannabaceae | Rosales |
| *Aglu_Alngl764S35691_NLP1* | *Alnus glutinosa* | NLP1 | 1 | Betulaceae | Fagales |
| *Cgla_Casgl109S07006_NLP1* | *Casuarina glauca* | NLP1 | 1 | Casuarinaceae | Fagales |
| *Jreg_WALNUT_00018800-RA_NLP1* | *Juglans regia* | NLP1 | 0 | Juglandaceae | Fagales |
| *Jreg_WALNUT_00016305-RA_NLP1* | *Juglans regia* | NLP1 | 0 | Juglandaceae | Fagales |
| *Ddru_Drydr83S05805_NLP1* | *Dryas drummmondii* | NLP1 | 1 | Rosaceae | Rosales |
| *Pcom_PCP004399.1_NLP1* | *Pyrus communis* | NLP1 | 0 | Rosaceae | Rosales |
| *Pper_Prupe.5G011200.1.p_NLP1* | *Prunus persica* | NLP1 | 0 | Rosaceae | Rosales |
| *Fves_mrna10914.1-v1.0-hybrid_NLP1* | *Fragaria vesca* | NLP1 | 0 | Rosaceae | Rosales |
| *Rocc_Bras_G13903_NLP1* | *Rubus occidentalis* | NLP1 | 0 | Rosaceae | Rosales |
| *Dtri_Distr887S29018_NLP1* | *Discaria trinervis* | NLP1 | 1 | Rhamnaceae | Rosales |
| *Zjuj_XP_015884123.1_NLP1* | *Ziziphus jujuba* | NLP1 | 0 | Rhamnaceae | Rosales |
| *Hlup_HL.SW.v1.0.G017874.1_NLP1* | *Humulus lupulus* | NLP1 | 0 | Cannabaceae | Rosales |
| *Vvin_GSVIVT01013370001_NLP1* | *Vitis vinifera* | NLP1 | 0 | Vitaceae | Vitales |
| *Mesc_Manes.05G130800.1.p_NLP1* | *Manihot esculenta* | NLP1 | 0 | Euphorbiaceae | Malpighiales |
| *Mesc_Manes.18G004300.1.p_NLP1* | *Manihot esculenta* | NLP1 | 0 | Euphorbiaceae | Malpighiales |
| *Rcom_30170.m014365_NLP1* | *Ricinus communis* | NLP1 | 0 | Euphorbiaceae | Malpighiales |
| *Ptrc_Potri.005G251700.1_NLP1* | *Populus trichocarpa* | NLP1 | 0 | Salicaceae | Malpighiales |
| *Ptrc_Potri.002G009700.1_NLP1* | *Populus trichocarpa* | NLP1 | 0 | Salicaceae | Malpighiales |
| *Slyc_Solyc04g082480.2.1_NLP1* | *Solanum lycopersicum* | NLP1 | 0 | Solanaceae | Solanales |
| *Ccan_Cerca189S06512_NLP1* | *Cercis canadensis* | NLP1 | 0 | Fabaceae | Fabales |
| *Ccan_Cerca58S27147_NLP1* | *Cercis canadensis* | NLP1 | 0 | Fabaceae | Fabales |
| *Cari_Ca_14561.1_NLP1* | *Cicer arietinum* | NLP1 | 1 | Fabaceae | Fabales |
| *Mtru_Medtr3g115400.2_NLP1* | *Medicago truncatula* | NLP1 | 1 | Fabaceae | Fabales |
| *Ccaj_C.cajan_29749_NLP1* | *Cajanus cajan* | NLP1 | 1 | Fabaceae | Fabales |
| *Pvul_Phvul.009G011200.1_NLP1* | *Phaseolus vulgaris* | NLP1 | 1 | Fabaceae | Fabales |

| | | | | | |
|---|---|---|---|---|---|
| *Vrad_Vradi05g14430.1_NLP1* | *Vigna radiata* | NLP1 | 1 | Fabaceae | Fabales |
| *Gmax_Glyma.04G017400.1.p_NLP1* | *Glycine max* | NLP1 | 1 | Fabaceae | Fabales |
| *Gmax_Glyma.06G017800.1.p_NLP1* | *Glycine max* | NLP1 | 1 | Fabaceae | Fabales |
| *Caus_Castanospermum00843-PA_NLP1* | *Castanospermum australe* | NLP1 | 0 | Fabaceae | Fabales |
| *Ljap_Lj1g3v2295200.1_NLP1* | *Lotus japonicus* | NLP1 | 1 | Fabaceae | Fabales |
| *Nsch_Nissc337S21912_NLP1* | *Nissolia schottii* | NLP1 | 0 | Fabaceae | Fabales |
| *Aipa_Araip_Araip.B07_96852500..96856378_NLP1* | *Arachis ipaensis* | NLP1 | 1 | Fabaceae | Fabales |
| *Cfas_Chafa97S13327_NLP1* | *Chamaecrista fasciculata* | NLP1 | 1 | Fabaceae | Fabales |
| *Mpud_Mimpu_merged_scaffold9844+scaffold5711_NLP1* | *Mimosa pudica* | NLP1 | 1 | Fabaceae | Fabales |
| *Cfas_Chafa95053S28562_NLP1* | *Chamaecrista fasciculata* | NLP1 | 1 | Fabaceae | Fabales |
| *Dglo_Datgl151S21250_NLP1* | *Datisca glomerata* | NLP1 | 1 | Datiscaceae | Cucurbitales |
| *Bfuc_Begfu1351S18529_NLP* | *Begonia fuchsioides* | NLP1 | 0 | Begoniaceae | Cucurbitales |
| *Bfuc_Begfu1129S16439_NLP1* | *Begonia fuchsioides* | NLP1 | 0 | Begoniaceae | Cucurbitales |
| *Mgut_Migut.N02963.1.p_NIN* | *Mimulus guttatus* | NLP1 | 0 | Phrymaceae | Lamiales |
| *Lusi_Lus10016018_NLP1* | *Linum usitatissimum* | NLP1 | 0 | Linaceae | Malpighiales |
| *Lusi_Lus10012257_NLP1* | *Linum usitatissimum* | NLP1 | 0 | Linaceae | Malpighiales |
| *Csat_Csa4M658610.1_NLP1* | *Cucumis sativus* | NLP1 | 0 | Cucurbitaceae | Cucurbitales |
| *Clan_Cla023449_NLP1* | *Citrullus lanatus* | NLP1 | 0 | Cucurbitaceae | Cucurbitales |
| *Bfuc_Begfu1417S19338_NLP1* | *Begonia fuchsioides* | NLP1 | 0 | Begoniaceae | Cucurbitales |
| *Bfuc_Begfu745S40752_NLP1* | *Begonia fuchsioides* | NLP1 | 0 | Begoniaceae | Cucurbitales |
| *Bfuc_Begfu1777S22537_NLP1* | *Begonia fuchsioides* | NLP1 | 0 | Begoniaceae | Cucurbitales |
| *Bfuc_Begfu4799S04580_NLP1* | *Begonia fuchsioides* | NLP1 | 0 | Begoniaceae | Cucurbitales |
| *Lusi_Lus10023930_NIN* | *Linum usitatissimum* | NIN | 0 | Linaceae | Malpighiales |
| *Lusi_Lus10014428_NIN* | *Linum usitatissimum* | NIN | 0 | Linaceae | Malpighiales |
| *Mesc_Manes.07G059900.1.p_NIN* | *Manihot esculenta* | NIN | 0 | Euphorbiaceae | Malpighiales |

| | | | | | |
|---|---|---|---|---|---|
| *Ptrc_Potri.016G042900.1_NIN* | *Populus trichocarpa* | NIN | 0 | Salicaceae | Malpighiales |
| *Ptrc_Potri.009G166400.1_NIN* | *Populus trichocarpa* | NIN | 0 | Salicaceae | Malpighiales |
| *Cfas_Chafa4076S22354_NIN* | *Chamaecrista fasciculata* | NIN | 1 | Fabaceae | Fabales |
| *Ljap_Lj2g3v3373100.1_NIN* | *Lotus japonicus* | NIN | 1 | Fabaceae | Fabales |
| *Aipa_Araip.10007239.1_NIN* | *Arachis ipaensis* | NIN | 1 | Fabaceae | Fabales |
| *Mtru_Medtr5g099060.1_NIN* | *Medicago truncatula* | NIN | 1 | Fabaceae | Fabales |
| *Cari_Ca_03225.1_NIN* | *Cicer arietinum* | NIN | 1 | Fabaceae | Fabales |
| *Ccaj_C.cajan_33924_NIN* | *Cajanus cajan* | NIN | 1 | Fabaceae | Fabales |
| *Gmax_Glyma.04G000600.1.p_NIN* | *Glycine max* | NIN | 1 | Fabaceae | Fabales |
| *Vrad_Vigra_Vr05_24677902..24680507_NIN* | *Vigna radiata* | NIN | 1 | Fabaceae | Fabales |
| *Pvul_Phvul.009G115800.1_NIN* | *Phaseolus vulgaris* | NIN | 1 | Fabaceae | Fabales |
| *Mpud_Mimpu1984S19289_NIN* | *Mimosa pudica* | NIN | 1 | Fabaceae | Fabales |

* Nodulator = 1 and Non-nodulator = 0

**Supplementary Table 2: Metadata for the OrthoFinder analysis**

| **Gene_ID** | **Scientific name** | **References** | **order** | **family** | **RNS** |
|---|---|---|---|---|---|
| Allnan_v1r1_augustus.hints_longest_isoform | *Allocasuarina nana* | This study | Fagales | Casuarinaceae | Y |
| Alnglu_longest_isoform | *Alnus glutinosa* | [2] | | | |
| Aratha.TAIR10.pep.all | *Arabidopsis thaliana* | [3] | | | |
| Begfuc_longest_isoform | *Begonia fuchsioides* | [2] | | | |
| Carfan_longest_isoform | *Carpinus fangiana* | https://www.ncbi.nlm.nih.gov/datasets/genome/GCA_006937295.1/ | | | |
| Casgla_longest_isoform | *Casuarina glauca* | [2] | | | |
| Casmol_longest_isoform | *Castanea mollissima* | [4] | | | |
| Ceathy_v1r1_augustus.hints_longest_isoform | *Ceanothus thyrsiflorus* | This study | Rosales | Rhamnaceae | Y |
| Cermon_v1r1_augustus.hints_longest_isoform | *Cercocarpus montanus* | This study | Rosales | Rosaceae | Y |
| Colpar_v1r1_augustus.hints_longest_isoform | *Colletia paradoxa* | This study | Rosales | Rhamnaceae | Y |
| Coluli_v1r1_augustus.hints_longest_isoform | *Colletia ulicina* | This study | Rosales | Rhamnaceae | Y |
| Corjap_v1r1_augustus.hints_longest_isoform | *Coriaria japonica* | This study | Cucurbitales | Coriariaceae | Y |
| Corlae_v1r1_augustus.hints_longest_isoform | *Corynocarpus laevigatus* | This study | Cucurbitales | Corynocarpaceae | N |
| Cucmax_longest_isoform | *Cucurbita maxima* | [5] | | | |
| Cucsat_longest_isoform | *Cucumis sativus PI183967* | [6] | | | |
| Datcan_v1r1_augustus.hints_longest_isoform | *Datisca cannabina* | This study | Cucurbitales | Datiscaceae | Y |
| Datglo_longest_isoform | *Datisca glomerata* | [2] | | | |
| Datglo_v3_pep | *Datisca glomerata* | This study | | | |
| Distri_longest_isoform | *Discaria trinervis* | [2] | | | |
| Drydru_v2r1_augustus.hints_longest_isoform | *Dryas drummondii* | This study | Rosales | Rosaceae | Y |

| | | | | | |
|---|---|---|---|---|---|
| Dryoct_v2r1_augustus.hints_longest_isoform | *Dryas octopetala* | This study | Rosales | Rosaceae | N |
| Elaang_v1r1_augustus.hints_longest_isoform | *Elaeagnus angustifolia* | This study | Rosales | Elaeagnaceae | Y |
| Filulm_v1r1_augustus.hints_longest_isoform | *Filipendula ulmaria* | This study | Rosales | Rosaceae | N |
| Fraiin_longest_isoform | *Fragaria iinumae* | [7] | | | |
| Fraves_longest_isoform | *Fragaria vesca* | [8] | | | |
| Fvesca_677_v4.0.a2.protein_primaryTranscriptOnly | *Fragaria vesca* | [9] | | | |
| Glymax_longest_isoform | *Glycine max* | [10] | | | |
| Hiprha_longest_isoform | *Hippophae rhamnoides* | [11] | | | |
| Hipsal_v1r1_augustus.hints_longest_isoform | *Hippophae salicifolia* | This study | Rosales | Elaeagnaceae | Y |
| Horvul_Golden_Promise_v1r1_Apollo_300620_prot | *Hordeum vulgare* | [12] | | | |
| Jugreg_longest_isoform | *Juglans regia* | [13] | | | |
| Lotjap_longest_isoform | *Lotus japonicus* | [14] | | | |
| Maldom_GDDH13_1-1 | *Malus domestica* | https://www.rosaceae.org/species/malus/malus_x_domestica/genome_v3.0.a1 | | | |
| Medtru_longest_isoform | *Medicago truncatula* | [15] | | | |
| MtrunA17r5.0-ANR-EGN-r1.9.prot | *Medicago truncatula* | [16] | | | |
| Mtruncatula_285_Mt4.0v1.protein_primaryTranscriptOnly | *Medicago truncatula* | [17] | | | |
| Myrcer_v1r1_augustus.hints_longest_isoform | *Myrica cerifera* | This study | Fagales | Myricaceae | Y |
| Parand_longest_isoform | *Parasponia andersonii* | [18] | | | |
| Pomvac_v1r1_augustus.hints_longest_isoform | *Pomaderris vacciniifolia* | This study | Rosales | Rhamnaceae | N |
| Potmic_Potentilla_transcripts.translated_cds | *Potentilla micrantha* | [19] | | | |
| Prumum_longest_isoform | *Prunus mume* | [20] | | | |
| Pruper_longest_isoform | *Prunus persica* | [21] | | | |
| Purtri_v1r1_augustus.hints_longest_isoform | *Purshia tridentata* | This study | Rosales | Rhamnaceae | N |

| | | | | | |
|---|---|---|---|---|---|
| Pyrcom_longest_isoform | *Pyrus communis* | [22] | | | |
| Rharub_longest_isoform | *Rhamnella rubrinervis* | https://www.ncbi.nlm.nih.gov/datasets/genome/GCA_007844105.2/ | | | |
| Roschi_longest_isoform | *Rosa chinensis* | [23] | | | |
| Rubell_v1r1_augustus.hints_longest_isoform | *Rubus ellipticus* | This study | Rosales | Rhamnaceae | N |
| Rubocc_longest_isoform | *Rubus occidentalis* | [24] | | | |
| Sagthe_v1r1_augustus.hints_longest_isoform | *Sageretia theezans* | This study | Rosales | Rhamnaceae | N |
| Shearg_v1r1_augustus.hints_longest_isoform | *Shepherdia argentea* | This study | Rosales | Elaeagnaceae | Y |
| Zizjuj_longest_isoform | *Ziziphus jujuba cv. Dongzao* | [25] | | | |

**Supplementary Table 3: Transformation efficiency in *Fragaria vesca* cv. Hawaii4**

| **Construct** | **Number of transformed explant** | **Number of calli with shoots obtained** |
|---|---|---|
| *pLjUBI-GUS-tNOS-p35S-mCherry-t35S* | 50 | 40 |
| *pLjUBI-DdNIN-3xMyc-trbcs-p35S-mCherry-t35S* | 50 | 0 |
| *pDR5-DdNIN_cDNA-tNOS-p35S-mCherry-t35S* | 50 | 21 |

**Supplementary Table 4: Construct assembly**

| EC number | Level | Construct details |
|---|---|---|
| *EC37362* | 0 | *pL0M-PU-pDatiscaNIN_F1-5-37362* |
| *EC54188* | 0 | *pL0B-PU-pDatiscaNIN2_F1-5-54188* |
| *EC37363* | 0 | *pL0-PU-pDryasNIN_F1-5-37363* |
| *EC55328* | 0 | *pL0M-PU-pMtNIN(3C)+3XMtCYC+35S-55328* |
| *EC25116* | 0 | *pL0M-PU-pDR5* |
| *EC15057* | 0 | *pL0M-PU-pNOS-15057* |
| *EC51266* | 0 | *pL0M-PU-35S-TMV-1-51266* |
| *EC15251* | 0 | *pL0M-PU-LjUBI-15251* |
| *EC54611* | 0 | *pL0B-PU-pMtNINsyn-54611* |
| *EC54614* | 0 | *pL0B-PU-pMtNINsynAuxREmut-54614* |
| *EC54663* | 0 | *pL0-PU-pMtNINsynNIN-BEmut-54663* |
| *EC54638* | 0 | *pL0B-PU-pDR5_pMtNINsyn_F2-54638* |
| *EC37364* | 0 | *pL0-PU-pDryasRPG_F1-5-37364* |
| *EC54535* | 0 | *pL0M-S-n2-Staygold-CO-54535* |
| *EC75111* | 0 | *pL0M-SC-GUS-intron-75111* |
| *EC15068* | 0 | *pL0M-SC-Kan-15068* |
| *EC15071* | 0 | *pL0M-SC-mCherry-15071* |
| *EC61119* | 0 | *pL0M-SC-MtNIN(CDS)-61119* |
| *EC54016* | 0 | *pL0B-SC-DgNIN_cDNA-F1-3(CDS)-54016* |
| *EC54189* | 0 | *pL0B-SC-DatiscaNIN2-cDNA_F1-2-54189* |
| *EC54041* | 0 | *pL0B-SC-Dryas_drumNIN_cDNA_F1-3(CDS)-54041* |
| *EC15318* | 0 | *pL0M-T-Rbcs-15318* |
| *EC41414* | 0 | *pL0M-T-35S-1-41414* |
| *EC41421* | 0 | *pL0M-T-Nos-41421* |
| *EC41432* | 0 | *pL0M-T-ocs-1-41432* |
| *EC15069* | 0 | *pL0M-SC-Hyg-15069* |
| *EC54007* | 1 | *pL1B-R2-pDatiscaNIN-GUS-tocs-54007* |
| *EC15029* | 1 | *pL1M-R1-pNOS-Kan-tNOS-15029* |
| *EC54091* | 1 | *pL1B-R3-p35S-mcherry-t35S-54091* |
| *EC54198* | 1 | *pL1B-R2-pDgNIN2-GUS-tOcs-54198* |
| *EC54115* | 1 | *pL1B-R2-pDryasNIN-GUS-tNOS-54115* |
| *EC54121* | 1 | *pL1B-R2-pLjUBI-GUS-tNOS-54121* |
| *EC54019* | 1 | *pL1B-R2-pDR5-GUS-tNOS-54019* |
| *EC54197* | 1 | *pL1B-R2-pDR5-DdNIN_cDNA-tNOS-54197* |
| *EC15030* | 1 | *pL1M-R1-p35S-HYG-tNOS* |
| *EC54542* | 1 | *pL1B-R2-pDR5-MtNIN_CDS-tNOS-54542* |

| | | |
|---|---|---|
| *EC54118* | 1 | *pL1B-R3-pLjUBI-3xHA-DdNIN-trbcs-54118* |
| *EC54125* | 1 | *pL1B-R2-p35S-mcherry-35S-54125* |
| *EC54314* | 1 | *pL1B-R3-pDryasRPG-GUS-tNOS-54314* |
| *EC54562* | 1 | *pL1B-R3-pLjUBI-DgNIN-FLAG-tNOS-54562* |
| *EC54564* | 1 | *pL1B-R3-pLjUBI-DgNIN2-FLAG-tNOS-54564* |
| *EC54541* | 1 | *pL1B-R2-pMtNIN(3C)+3XMtCYC+35S-GUS-tNOS-54541* |
| *EC54543* | 1 | *pL1B-R2-pMtNIN(3C)+3XMtCYC+35S-MtNIN_CDS-tNOS-54543* |
| *EC54544* | 1 | *pL1B-R2-pMtNIN(3C)+3XMtCYC+35S-DgNIN_CDS-tNOS-54544* |
| *EC54545* | 1 | *pL1B-R2-pMtNIN(3C)+3XMtCYC+35S-DdNIN_CDS-tNOS-54545* |
| *EC54603* | 1 | *pL1B-R2-pMtNIN(3C)+3XMtCYC+35S-DgNIN2_CDS-tNOS-54603* |
| *EC54574* | 1 | *pL1B-R2-pLjUBI-Staygold-MtNIN-tNOS-54574* |
| *EC54656* | 1 | *pL1B-R2-pLjUBI-Staygold-DatiscaNIN-tNOS-54656* |
| *EC54657* | 1 | *pL1B-R2-pLjUBI-Staygold-DryasNIN-tNOS-54657* |
| *EC54027* | 1 | *pL1B-R2-pDgNIN-DgNIN_CDS-tNOS-54027* |
| *EC54327* | 1 | *pL1B-R2-pDdNIN-DdNIN_CDS-tNOS-54327* |
| *EC54264* | 1 | *pENTR3c-DgNIN-RNAi-54264* |
| *EC54621* | 1 | *pL1B-R2-pMtNINsyn-MtNIN-tNOS-54621* |
| *EC54622* | 1 | *pL1B-R2-pMtNINsynAuxREmut-MtNIN-tNOS-54622* |
| *EC54664* | 1 | *pL1B-R2-pMtNINsyn-NIN_BEmut-MtNIN-tNOS-54664* |
| *EC54629* | 1 | *pL1B-R2-pMtNINsyn-GUS-tNOS-54629* |
| *EC54630* | 1 | *pL1B-R2-pMtNINsynAuxREmut-GUS-tNOS-54630* |
| *EC54640* | 1 | *pL1B-R2-pDR5_pMtNINsyn_F2-MtNIN-tNOS-54640* |
| *EC54286* | 1 | *pL1B-R1-p35S-mcherry-t35S-54286* |
| *EC54010* | 2 | *pL2B-Kan-pDgNIN-GUS-tNOS-p35S-mCherry-t35S-54010* |
| *EC54203* | 2 | *pL2B-Kan-pDgNIN2-GUS-tOcs-p35S-mCherry-t35S-54203* |
| *EC54119* | 2 | *pL2B-Kan-pDdNIN-GUS-tNOS-p35S-mCherry-t35S-54119* |
| *EC54120* | 2 | *pL2B-Kan-pDdRPG-GUS-tNOS-p35S-mcherry-t35S-54120* |
| *EC54122* | 2 | *pL2B-Kan-pLjUBI-GUS-tNOS-p35S-mCherry-t35S-54122* |
| *EC54096* | 2 | *pL2B-Kan-pDR5-GUS-tNOS-p35S-mCherry-t35S-54096* |
| *EC54097* | 2 | *pL2B-Kan-pDR5-DgNIN_CDS-tNOS-p35S-mCherry-t35S-54097* |
| *EC54208* | 2 | *pL2B-Kan-pDR5-DdNIN_CDS-tNOS-p35S-mCherry-t35S-54208* |
| *EC54371* | 2 | *pL2B-Hyg-pDR5-DdNIN_cDNA-tNOS-p35S-mCherry-t35S-54371* |
| *EC54549* | 2 | *pL2B-Kan-pDR5-MtNIN_CDS-tNOS-p35S-mCherry-t35S-54549* |
| *EC54123* | 2 | *pL2B-Kan-pLjUBI-DdNIN-3xMyc-trbcs-p35S-mCherry-t35S-54123* |
| *EC54323* | 2 | *pL2B-Kan-pDR5-DdNIN-tNOS-pDryasRPG-GUS-tNOS-p35S-mCherry-t35S-54323* |
| *EC54570* | 2 | *pL2B-Kan-p35S-mCherry-t35S-pLjUBI-DgNIN-FLAG-tNOS-54570* |
| *EC54572* | 2 | *pL2B-Kan-p35S-mCherry-t35S-pLjUBI-DgNIN2-FLAG-tNOS-54572* |
| *EC54548* | 2 | *pL2B-Kan-pMtNIN(3C)+3XMtCYC+35S-GUS-tNOS-p35S-mCherry-t35S-54548* |

| | | |
|---|---|---|
| *EC54550* | 2 | *pL2B-Kan-pMtNIN(3C)+3XMtCYC+35S-MtNIN_CDS-tNOS-p35S-mCherry-t35S-54550* |
| *EC54551* | 2 | *pL2B-Kan-pMtNIN(3C)+3XMtCYC+35S-DgNIN_CDS-tNOS-p35S-mCherry-t35S-54551* |
| *EC54552* | 2 | *pL2B-Kan-pMtNIN(3C)+3XMtCYC+35S-DdNIN_CDS-tNOS-p35S-mCherry-t35S-54552* |
| *EC54617* | 2 | *pL2B-Kan-pMtNIN(3C)+3XMtCYC+35S-DgNIN2_CDS-tNOS-p35S-mCherry-t35S-54617* |
| *EC54578* | 2 | *pL2B-Kan-pLjUBI-Staygold-MtNIN-tNOS-p35S-mCherry-t35S-54578* |
| *EC54658* | 2 | *pL2B-Kan-pLjUBI-Staygold-DatiscaNIN-tNOS-p35S-mCherry-t35S-54658* |
| *EC54659* | 2 | *pL2B-Kan-pLjUBI-Staygold-DryasNIN-tNOS-p35S-mCherry-t35S-54659* |
| *EC54324* | 2 | *pL2B-Kan-pDgNIN-DgNIN_CDS-tNOS-p35S-mCherry-t35S-54324* |
| *EC54325* | 2 | *pL2B-Kan-pDdNIN-DdNIN_CDS-tNOS-p35S-mCherry-t35S-54325* |
| *EC54272* | 2 | *pK7GWIWG2-7F2.1-Kan-p35S-DgNIN-RNAi-t35S-p35S-eGFP-t35S-54272* |
| *EC54623* | 2 | *pL2B-Kan-pMtNINsyn-MtNIN-tNOS-p35S-mCherry-t35S-54623* |
| *EC54624* | 2 | *pL2B-Kan-pMtNINsynAuxREmut-MtNIN-tNOS-p35S-mCherry-t35S-54624* |
| *EC54665* | 2 | *pL2B-Kan-pMtNINsyn-NIN_BEmut-MtNIN-tNOS-p35S-mCherry-t35S-54665* |
| *EC54634* | 2 | *pL2B-p35S-mCherry-t35S-pMtNINsyn-MtNIN-tNOS-pDR5-GUS-trbcs-54634* |
| *EC54635* | 2 | *pL2B-p35S-mCherry-t35S-pMtNINsynAuxREmut-MtNIN-tNOS-pDR5-GUS-trbcs-54635* |
| *EC54631* | 2 | *pL2B-Kan-pMtNINsyn-GUS-tNOS-p35S-mCherry-t35S-54631* |
| *EC54632* | 2 | *pL2B-Kan-pMtNINsynAuxREmut-GUS-tNOS-p35S-mCherry-t35S-54632* |
| *EC54642* | 2 | *pL2B-p35S-mCherry-t35S-pDR5_pMtNINsyn_F2-MtNIN-tNOS-54642* |

**Supplementary Table 5: Primer list**

| **Primer name** | **Sequence** |
|---|---|
| *DgActin qRT-for* | *ggaatggaagctgctggaatc* |
| *DgActin qRT-rev* | *ggtctgcaatacctgggaac* |
| *DgNIN qRT-for* | *ggttagaggtaacagactcatccattc* |
| *DgNIN qRT-rev* | *caggatcttcatgatcttggtagattg* |
| *DgNIN2 qRT-for* | *caaggacgttggatggattcctcaaatgg* |
| *DgNIN2 qRT-rev* | *cgtgattggcctgatcttcttcgtag* |
| *DryasNIN qRT-for* | *aaactgatatcttctcccacag* |
| *DryasNIN qRT-rev* | *ctcttcctttggttcttctgtttg* |
| *DgNIN RNAi BamH1 for* | *ccgggatccgctatggtcagcaggttgatg* |
| *DgNIN RNAi Xho1 rev* | *cgcctcgagcagaatgcttgtaacaccag* |
| *FvHisH4 qRT-for* | *tcaagcgtatctccggtctc* |
| *FvHisH4 qRT-rev* | *agtgtccttccctgcctctt* |
| *MtNIN qRT-for* | *attgcaaggcgatttaacctaaca* |
| *MtNIN qRT-rev* | *gagaggggaagcttgaaaaagaga* |
| *MtEF1alpha-for* | *ctttgcttggtgctgtttagatgg* |
| *MtEF1alpha-rev* | *attccaaaggcggctgcata* |

**Supplementary Dataset 1.** Sequence information for the Maximum-likelihood phylogenetic tree of NIN/NLP1 clade

**Supplementary Dataset 2.** (**A**) *Datisca glomerata* DEGs for the hairy-root transformed plants (**B**) Expression trend between *Empty vector*, *DgNIN_RNAi* and *NIN* overexpression. (**C**) Expression trend between *DgNIN* and *DgNIN2* overexpression in *Datisca glomerata* versus *MtNIN* overexpression in *Medicago truncatula*.

**Supplementary Dataset 3.** OrthoFinder analysis Orthogroups

**Supplementary Dataset 4.** AuxRE analysis of the promoter region for NIN and NLPs. (**A**) Promoter sequences, (**B**) Promoter length, (**C**) MEME for *AuxRE* and (**D**) FIMO output.

**Supplementary Dataset 5.** (**A**) *Fragaria vesca* DEGs for *pDR5:DryasNIN* post treatment with auxin under Nitrogen (N) repleted and depleted condition. (**B**) HOG median expression trend between *pDR5:DryasNIN(+N+NAA)* in *F. vesca*, *pDR5:DdNIN (-Inf)* in *D. glomerata* and *MtNIN* overexpression in *M. truncatula*. (**C**) Heatmap dataset between *Fv* and *Mt*.

**Supplementary Dataset 6.** (**A**) Barley (*Hordeum vulgare*) DEGs for *pDR5:DryasNIN* post treatment with auxin under Nitrogen (N) repleted and depleted condition. (**B**) HOG median expression trend between *pDR5:DryasNIN(-N+NAA)* in *H. vulgare*, *pDR5:DdNIN (-Inf)* in *D. glomerata* and *MtNIN* overexpression in *M. truncatula*. (**C**) Heatmap dataset between *Hv* and *Mt*.

### References


1. Lee, T. *et al.* Light-sensitive short hypocotyl genes confer symbiotic nodule identity in the legume Medicago truncatula. *Current Biology* **34**, 825-840.e7 (2024).
2. Griesmann, M. *et al.* Phylogenomics reveals multiple losses of nitrogen-fixing root nodule symbiosis. *Science (1979).* **361**, (2018).
3. Lamesch, P. *et al.* The Arabidopsis Information Resource (TAIR): Improved gene annotation and new tools. *Nucleic Acids Res.* **40**, (2012).
4. Wang, J. *et al.* Construction of pseudomolecules for the chinese chestnut (Castanea mollissima) genome. *G3: Genes, Genomes, Genetics* **10**, 3565–3574 (2020).
5. Sun, H. *et al.* Karyotype Stability and Unbiased Fractionation in the Paleo-Allotetraploid Cucurbita Genomes. *Mol. Plant* **10**, 1293–1306 (2017).
6. Li, Q. *et al.* A chromosome-scale genome assembly of cucumber (Cucumis sativus L.). *Gigascience* **8**, (2019).
7. Edger, P. P. *et al.* Origin and evolution of the octoploid strawberry genome. *Nat. Genet.* **51**, 541–547 (2019).
8. Edger, P. P. *et al.* Single-molecule sequencing and optical mapping yields an improved genome of woodland strawberry (Fragaria vesca) with chromosome-scale contiguity. *GigaScience* vol. 7 Preprint at https://doi.org/10.1093/gigascience/gix124 (2018).
9. Li, Y., Pi, M., Gao, Q., Liu, Z. & Kang, C. Updated annotation of the wild strawberry Fragaria vesca V4 genome. *Hortic. Res.* **6**, (2019).
10. Valliyodan, B. *et al.* Construction and comparison of three reference-quality genome assemblies for soybean. *Plant Journal* **100**, 1066–1082 (2019).
11. Wu, Z. *et al.* Genome of Hippophae rhamnoides provides insights into a conserved molecular mechanism in actinorhizal and rhizobial symbioses. *New Phytologist* **235**, 276–291 (2022).
12. Schreiber, M. *et al.* A genome assembly of the barley 'transformation reference' cultivar golden promise. *G3: Genes, Genomes, Genetics* **10**, 1823–1827 (2020).
13. Peng, S., Yang, G., Liu, C., Yu, Z. & Zhai, M. The complete chloroplast genome of the Juglans regia (Juglandales: Julandaceae). *Mitochondrial DNA Part A* **28**, 407–408 (2017).

14. Li, H., Jiang, F., Wu, P., Wang, K. & Cao, Y. A high-quality genome sequence of model legume lotus japonicus (Mg-20) provides insights into the evolution of root nodule symbiosis. *Genes (Basel).* **11**, (2020).
15. Young, N. D. *et al.* The Medicago genome provides insight into the evolution of rhizobial symbioses. *Nature* **480**, 520–524 (2011).
16. Pecrix, Y. *et al.* Whole-genome landscape of Medicago truncatula symbiotic genes. *Nature Plants* vol. 4 1017–1025 Preprint at https://doi.org/10.1038/s41477-018-0286-7 (2018).
17. Tang, H. *et al.* An improved genome release (version Mt4.0) for the model legume Medicago truncatula. *BMC Genomics* **15**, 312 (2014).
18. van Velzen, R. *et al.* Comparative genomics of the nonlegume *Parasponia* reveals insights into evolution of nitrogen-fixing rhizobium symbioses. *Proceedings of the National Academy of Sciences* **115**, (2018).
19. Buti, M. *et al.* The genome sequence and transcriptome of Potentilla micrantha and their comparison to Fragaria vesca (the woodland strawberry). *Gigascience* **7**, 1–14 (2018).
20. Zhang, Q. *et al.* The genome of Prunus mume. *Nat. Commun.* **3**, (2012).
21. Verde, I. *et al.* The Peach v2.0 release: High-resolution linkage mapping and deep resequencing improve chromosome-scale assembly and contiguity. *BMC Genomics* **18**, (2017).
22. Chagné, D. *et al.* The draft genome sequence of European pear (Pyrus communis L. 'Bartlett'). *PLoS One* **9**, (2014).
23. Raymond, O. *et al.* The Rosa genome provides new insights into the domestication of modern roses. *Nat. Genet.* **50**, 772–777 (2018).
24. VanBuren, R. *et al.* The genome of black raspberry (Rubus occidentalis). *Plant J.* **87**, 535–547 (2016).
25. Liu, M. J. *et al.* The complex jujube genome provides insights into fruit tree biology. *Nat. Commun.* **5**, (2014).
